# Hyperconnectivity of two separate long-range cholinergic systems contributes to the reorganization of the brain functional connectivity during nicotine withdrawal in male mice

**DOI:** 10.1101/2023.03.29.534836

**Authors:** Lieselot L.G. Carrette, Adam Kimbrough, Pasha A. Davoudian, Alex C. Kwan, Andres Collazo, Olivier George

**Affiliations:** Department of Psychiatry, UC San Diego, La Jolla, CA, 92032, United States; Medical Scientist Training Program, Yale University School of Medicine, New Haven, CT, 06511, United States; Interdepartmental Neuroscience Program, Yale University School of Medicine, New Haven, CT, 06511, United States; Meinig School of Biomedical Engineering, Cornell University, Ithaca, NY, 14853, United States; Beckman Institute, CalTech, Pasadena, CA, 91125, United States

**Keywords:** Single-cell whole-brain imaging, FOS reactivity, stimulant, addiction

## Abstract

Chronic nicotine results in dependence with withdrawal symptoms upon discontinuation of use, through desensitization of nicotinic acetylcholine receptors and altered cholinergic neurotransmission. Nicotine withdrawal is associated with increased whole-brain functional connectivity and decreased network modularity, however, the role of cholinergic neurons in those changes is unknown. To identify the contribution of nicotinic receptors and cholinergic regions to changes in the functional network, we analyzed the contribution of the main cholinergic regions to brain-wide activation of the immediate early-gene FOS during withdrawal in male mice and correlated these changes with the expression of nicotinic receptor mRNA throughout the brain. We show that the main functional connectivity modules included the main long-range cholinergic regions, which were highly synchronized with the rest of the brain. However, despite this hyperconnectivity they were organized into two anticorrelated networks that were separated into basal forebrain projecting and brainstem-thalamic projecting cholinergic regions, validating a long-standing hypothesis of the organization of the brain cholinergic systems. Moreover, baseline (without nicotine) expression of *Chrna2*, *Chrna3*, *Chrna10*, and *Chrnd* mRNA of each brain region correlated with withdrawal-induced changes in FOS expression. Finally, by mining the Allen Brain mRNA expression database, we were able to identify 1755 gene candidates and three pathways (Sox2-Oct4-Nanog, JAK-STAT, and MeCP2-GABA) that may contribute to nicotine withdrawal-induced FOS expression. These results identify the dual contribution of the basal forebrain and brainstem-thalamic cholinergic systems to whole-brain functional connectivity during withdrawal; and identify nicotinic receptors and novel cellular pathways that may be critical for the transition to nicotine dependence.

**Significance Statement:** Discontinuation of nicotine use in dependent users is associated with increased whole-brain activation and functional connectivity and leads to withdrawal symptoms. Here we investigated the contribution of the nicotinic cholinergic receptors and main cholinergic projecting brain areas in the whole-brain changes associated with withdrawal. This not only allowed us to visualize and confirm the previously described duality of the cholinergic brain system using this novel methodology, but also identify nicotinic receptors together with 1751 other genes that contribute, and could thus be targets for treatments against, nicotine withdrawal and dependence.

## Introduction

Chronic nicotine use causes adaptive changes throughout the brain that lead to drug dependence (Markou, 2008; Martin-Soelch, 2013; Fowler et al., 2020), the emergence of a withdrawal state following cessation, and long-lasting somatic and motivational symptoms (Le Foll and Goldberg, 2009) that contribute to relapse (Allen et al., 2008; Zhou et al., 2009). Brain states, like dependence and withdrawal, have been described through patterns of synchronous neural firing (Brown, 2006). Changes in the patterns of neuronal co-reactivity, also called the functional connectome, can be observed in humans and rodents during withdrawal from nicotine (Hobkirk et al., 2018; Cheng et al., 2019; Kimbrough et al., 2021). Whole-brain imaging with single-cell resolution using light-sheet microscopy on cleared brains (Renier et al., 2014; Renier et al., 2016; Ueda et al., 2020) has made the study of brain-wide functional networks at single-cell resolution possible by looking at the expression of the immediate-early gene FOS (Wheeler et al., 2013; Vetere et al., 2017; Kimbrough et al., 2020; Kimbrough et al., 2021; Smith et al., 2021; Roland et al., 2022), a marker of neuronal reactivity, which integrates neuronal activation during a period of 1-2h, an ideal temporal window to characterize nicotine withdrawal. Using this approach, we have found that mice in withdrawal exhibit a pronounced increase in coactivation patterns throughout the brain (Kimbrough et al., 2020; Kimbrough et al., 2021). This increased correlation between the brain regions caused the regions to cluster closer together and thus led to a significant decrease in whole-brain modularity, as measured by the decrease in the number of main clusters (after setting the hierarchical cluster dendrograms at half height). Increased functional connectivity throughout the network also resulted in a reduction of brain regions identified as hubs. Hub regions are regions with the highest intramodular and intermodular connectivity as measured using graph theory (participation coefficient, within-module degree). These hub regions are hypothesized to be the biggest drivers of neuronal activity within the network. For instance, during nicotine withdrawal, the main hub regions shifted from cortical (e.g. sensory, motor) to subcortical (e.g. amygdalar, thalamic, hypothalamic, and midbrain) regions (Kimbrough et al., 2020; Kimbrough et al., 2021). However, the role of cholinergic neurons and cholinergic receptors in the whole-brain functional hyperconnectivity observed during withdrawal is unknown.

Desensitization and upregulation of nAChRs_(Benwell et al., 1988; Balfour and Fagerstrom, 1996; Dani and Heinemann, 1996; Fowler et al., 2020) contributes to the emergence of nicotine withdrawal symptoms by altering cholinergic neurotransmission in brain regions critical to sensory processing (Gil and Metherate, 2019), attention (Hahn, 2015), emotion, and motivation (Leslie et al., 2013). nAChRs form pentameric structures assembled from a family of subunits composed of α_2_–α_10_ and β_2_–β_4_. Alpha4 and beta2 are the most prevalent, but all subunits are expressed throughout the brain. A large number of brain regions (40+) have cholinergic neurons, characterized by the expression of choline acetyltransferase (ChAT), however, most of them are interneurons and only eight brain regions have long-range projecting cholinergic neurons (Mesulam et al., 1983). The long-range cholinergic regions include Ch1 (medial septal nucleus; MS), Ch2 (vertical nucleus of the diagonal band; NDB), Ch3 (horizontal limb of the diagonal band nucleus; NDB), Ch4 (nucleus basalis of Meynert that consists of the magnocellular nucleus; MA and substantia innominata; SI), Ch5 (pedunculopontine nucleus; PPN), Ch6 (laterodorsal tegmental nucleus), Ch7 (medial habenula, MH), and Ch8 (parabigeminal nucleus) (Mesulam et al., 1983; Woolf, 1991). We hypothesized that following chronic nicotine administration, most cholinergic regions that are rich in nicotinic receptors would have a synchronized correlated activity due in part to the brain-wide upregulation of nicotinic receptors (Govind et al., 2009; Fowler et al., 2020) and the increase in cholinergic transmission during nicotine withdrawal (Carcoba et al., 2014). The increased correlation would lead to larger modules and decreased modularity. Furthermore, since cholinergic receptor signaling is critical for nicotine-induced FOS activation (Pang et al., 2016; Simmons et al., 2016), a subhypothesis was that the regional expression level of cholinergic-related genes would be correlated to regional differential FOS expression under withdrawal in nicotine dependent animals.

To test these hypotheses, we reanalyzed the previously published whole-brain nicotine withdrawal network (Kimbrough et al., 2021) focusing on the cholinergic regions using hierarchical clustering and graph-theory analysis, and investigated the relationship between baseline gene expression levels and FOS reactivity using the whole-brain in-situ Allen Brain expression database, which contains the regional whole-brain expression of 19,413 genes in the mouse genome (Lein et al., 2007; Davoudian et al., 2023). Contrary to our hypothesis, we found that during nicotine withdrawal, the cholinergic regions did not cluster together in a single module but were instead represented in each of the main modules and organized into two anticorrelated networks that were separated into basal forebrain projecting and brainstem-thalamic projecting cholinergic regions. Moreover, while mRNA expression of a few nicotinic receptors correlated with FOS activation, we identified a list of over 1000 candidate genes and three intracellular pathways, that may contribute to the reorganization of the whole-brain functional connectome during nicotine withdrawal.

## Materials and methods

This report includes a reanalysis of a previously acquired and published dataset (Kimbrough et al., 2021) consisting of FOS counts per brain region (175; available in the extended data Table 1-1) for two groups of male C57BL/6J mice (60 days old at the start of the experiment), 8h after removal from minipumps (Alzet; model 1002) that were implanted in the lower back to deliver nicotine (N=5, 24 mg/kg/d) or saline (N=4) for 7 days. This dose was chosen based on previous studies that indicated rewarding effects during use, resulting in withdrawal-like symptoms after the cessation of chronic use (Johnson et al., 2008; Stoker et al., 2012). The brains were harvested following perfusion (phosphate buffered saline, followed by 4% formaldehyde), postfixed overnight immunolabelled for FOS (primary: 1:2000, Synaptic Systems catalog #226003 and secondary: 1:500; Invitrogen, catalog #A31573, donkey anti-rabbit Alexa Fluor 647), cleared according to the iDISCO+ protocol, imaged using light-sheet microscopy (Ultramicroscope II; LaVision Biotec at 1.6x effective magnification at resolution of 4 x 4 μm and a 3 μm step size), and analyzed using the ClearMap package (Renier et al., 2016). Three brain regions with low-to-no FOS counts were excluded based on quality control of the original data: the dorsal premammilary nucleus, parabigeminal nucleus, and suprachiasmatic nucleus. These experiments had been conducted in strict adherence to the National Institutes of Health Guide for the Care and Use of Laboratory Animals and approved by The Scripps Research Institute Institutional Animal Care and Use Committee and by the Institutional Animal Care and Use Committee of the University of California. No new experimental procedures were performed for this manuscript. The data was processed similarly as previously published (Kimbrough et al., 2020; Kimbrough et al., 2021) using GraphPad Prism and R (used R functions are given between square brackets []), as described in more detail below.

### Functional connectome construction

The FOS counts per region obtained from the published dataset, were all increased by 1 and normalized to a log10 value to reduce variability, before calculating Pearson correlations between regions [cor(data, method = ‘pearson’)]. The matrix was then hierarchically clustered, based on the Euclidean distances calculated from the correlations [Heatmap.2(data)] Modules were obtained by cutting the clustering dendrogram at half height [cutree()].

### Average correlation calculations

Average R values were calculated for each treatment (saline or nicotine) within the basal forebrain cholinergic regions (n=3, excluding self-correlations), withing the brainstem-thalamic cholinergic regions (n=6, excluding self-correlations), and between both cholinergic subgroups (n=12). Average R values were also calculated per treatment for the interaction of all cholinergic regions with the major anatomical groups in the brain. Two-way ANOVA was then performed to examine the effect of treatment condition on the average R value for each comparison.

### Network analysis

Networks were analyzed for centrality [degree(graph)] or [betweenness(graph)] with the R package iGraph [graph_from_data_frame()]. Participation coefficient was obtained using a customized version of the bctpy Python package (https://github.com/aestrivex/bctpy), derived from the MATLAB implementation of Brain Connectivity Toolbox (Rubinov and Sporns, 2010).

### Analysis of expression data

Structure and gene expression data was extracted from the In Situ Hybridization gene expression database and Allen Brain Atlas (Lein et al., 2007) in Python, as published and described before (Fulcher and Fornito, 2016; Fulcher et al., 2019; Fulcher, 2020; Davoudian et al., 2023) and intersected with the 175 brain regions from the FOS dataset. The expression density of all nicotinic cholinergic receptors in the gene expression atlas was averaged across the experimental sets following centering and scaling per experiment. Next, the correlation of the baseline expression level (%-pixel) for every gene in every experiment was correlated with the FOS expression change in withdrawal compared to control (log-fold change) per brain region for every gene. The frontal pole cerebral cortex was excluded as an outlier.

### Reactome analysis

The gene set was analyzed using the Reactome pathway database on reactome.org by inserting the gene set, projected to human, and analyze without interactors (Joshi-Tope et al., 2005).

### Statistical analysis

Statistical analysis was performed as indicated in GraphPad Prism software or with R in R Studio, using twoway ANOVA with Tukey corrected post-hoc test or a Mann-Whitney U-test [wilcox.test()]. Results were considered significant with p < 0.05 (limit 5% in tests as false positives). In cases of multiple comparisons, p-values were corrected using Benjamini-Hochberg (BH) false discovery rate (FDR) with q = 0.05 (False Discovery Rate = 5%; limit 5% of significant results as potential false positives) as Bonferroni multiple comparison correction of p-values was considered too stringent (limit 5% of false positives in the results) [p.adjust(method = ‘BH’)]. Significance of Pearson correlations [cor(method = ‘pearson’)] was obtained with [cor.test(method = ‘pearson’)].

**Table 1.** Statistical table.

|  | **Data Structure**<br>**(Shapiro-Wilk test)** | **Type of test** | **Power (95% CI between**<br>**saline and nicotine mean)** |
| --- | --- | --- | --- |
| **a** | Normal | 2-way ANOVA | -0.88 to -0.14 |
| **b** | Normal | 2-way ANOVA | -0.40 to -0.27 |
| **c** | Non-normal | Mann-Whitney U-test | 36.00 to 50.00 |
| **d** | Non-normal | Mann-Whitney U-test | -63.03 to -27.80 |
| **e** | Non-normal | Mann-Whitney U-test | -0.20 to -0.11 |

### Data Visualization

Graphs were constructed using the R package ggplot2 included in the tidyverse package or the ggpubr package for barplots. Heatmaps were constructed using the R packages gplots and ComplexHeatmap. Networks were visualized by plotting [plot()] in the R package igraph or using Gephi software. Pathway illustrations were created with BioRender.com. Figures were combined and edited with Adobe Illustrator.

## Results

### Cholinergic groups are distributed throughout the nicotine withdrawal network

Using the methods summarized in **Figure 1A**, the nicotine and a saline control functional connectomes were obtained. Immunolabeling of immediate-early gene expression like *c-Fos* captured the neuronal reactivity over a period of 1-2 h during nicotine withdrawal. Automated registration onto an anatomical reference atlas using the ClearMap pipeline (Renier et al., 2016) then allowed for unbiased quantification of neuronal activation throughout the brain. Finally, based on synchronous reactivity between functionally connected brain regions, correlation analysis of the FOS counts allowed calculation of functional distances (**Fig. 1B-C, Ext. Fig. 1-1**) and construction of a whole-brain functional network that can be further analyzed using graph theory (**Fig. 1D-E**; thresholded for Pearson correlation > 0.75 and Ext. Fig. 1-2, 1-3). The nicotine withdrawal network consisted of 175 brain regions (nodes) with 4738 functional connections (edges), which was a 50% increase from the saline control that had 3019 functional connections. Hierarchical clustering of the correlation matrices with division of the dendrogram at half-height revealed nine modules, which was a clear decrease from the thirteen modules in the control network. The five main nicotine modules included both cortical and subcortical regions, and were named based on the regions with the most significant within-module influence based on the within-module degree z-score, which measures the intramodule connectivity or relative importance of a region within its own module (Kimbrough et al., 2021). The long-projection cholinergic groups did not cluster together in a single module as originally hypothesized, rather they were spread between the modules. All five main modules contained at least one of the 8 main long-projection cholinergic groups (Ch1-8, Circled in red in **Fig. 1D**): The largest cortico-mid-hindbrain module contains cholinergic group 5 – pedunculopontine nucleus (PPN). Next, the cortico-hypothalamic module contains cholinergic group 4 - magnocellular nucleus (MA). The intermediate cortico-hypothalamic module contains cholinergic group 1 - medial septum (MS). The smaller orbitofrontal-extended amygdalar module contains cholinergic groups 2-3 - diagonal band nucleus (NDB) and cholinergic group 4: substantia innominate (SI) and the midbrain-thalamo-habenular module cholinergic group 7 - habenula: medial (MH) and lateral (LH). The smallest modules did not have any cholinergic regions. Note that cholinergic groups 6 and 8 were omitted as they were too posterior for the imaging.

**Figure 1.**
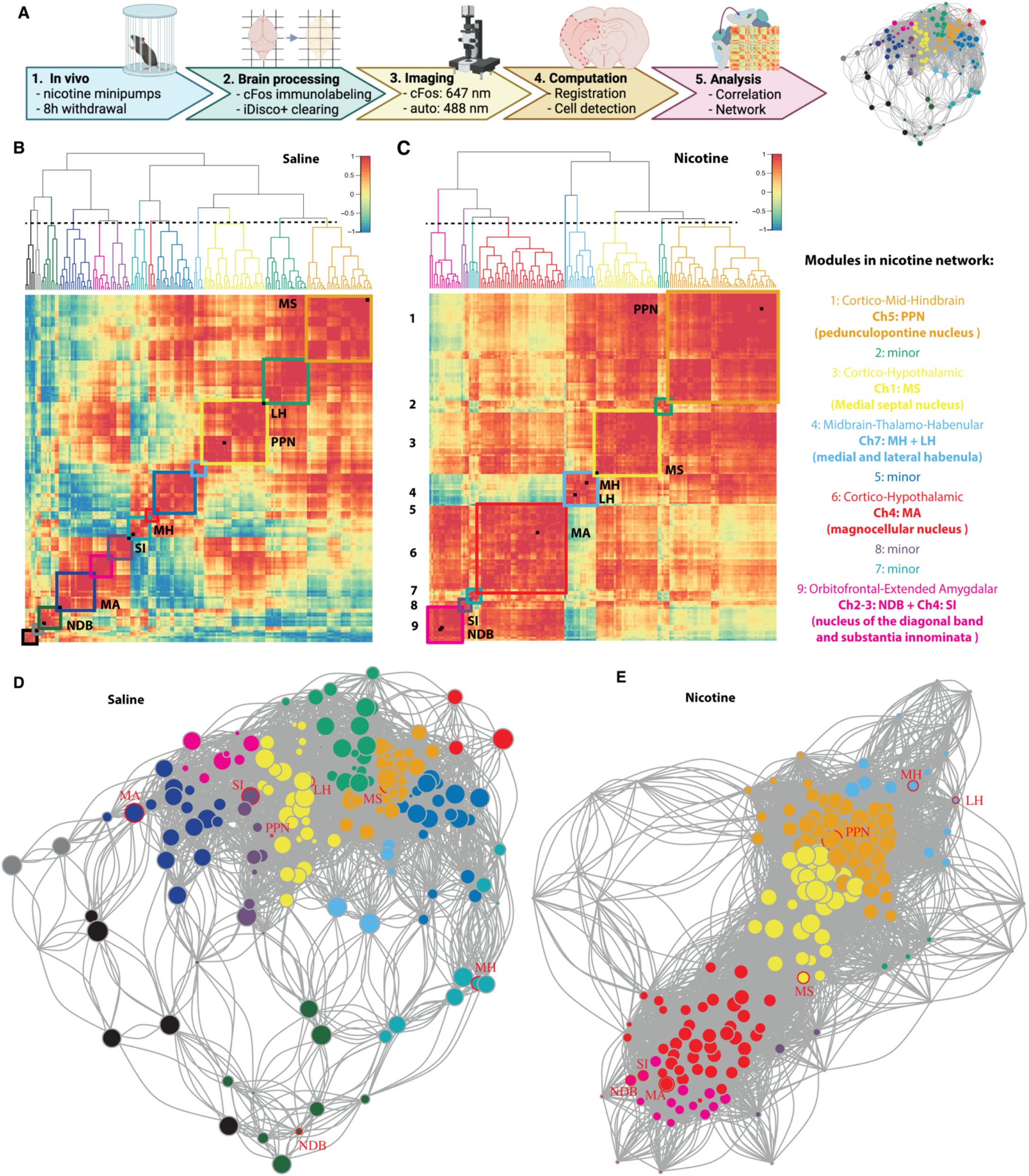
Functional network of nicotine withdrawal in mice. (**A)** Experimental timeline for obtaining the nicotine withdrawal functional connectome. 1) Mice were implanted subcutaneously with osmotic minipumps that delivered nicotine for 1 week, 8h after removal brains were harvested using perfusion. 2) The brains were immunolabeled for FOS and cleared using the iDISCO+ protocol. 3) Next the brains were imaged using light sheet imaging at 647 nm for FOS and 488 nm for autofluorescence. 4) Images were automatically registered to the Allen Brain Atlas and active cells counted per brain region using ClearMap. 5) The FOS cell counts of each region were correlated per group to obtain distances between the regions to create a network of the brain regions, that could be further analyzed. (**B-C)** Hierarchically clustered distance heatmaps of the resting state functional connectome (control, B) or nicotine-withdrawal (C), with 13 and 9 modules respectively depicted with colored squares and cholinergic regions labeled as black squares on the diagonal. The order of the brain regions in the heatmaps is available in the Extended Figure 1-1 (**D-E)** Network graph for saline (D) and nicotine withdrawal (E) with indication of cholinergic long-range regions (red circles). The node colors represent the different modules and the size the degree (number of connections). A larger image of the network with labeled nodes is available as Extended Figures 1-1 and 1-2. Abbreviations: medial septal nucleus (MS), nucleus of the diagonal band (NDB), magnocellular nucleus (MA), substantia innominata (SI), pedunculopontine nucleus (PPN), medial habenula (MH), and lateral habenula (LH).

### Increased interaction of the long-range cholinergic groups throughout the brain in 2 subsystems

To investigate the role of the cholinergic regions in the organization of the whole-brain network, we first tested whether the long-range cholinergic groups (**Fig. 2A**) were significantly correlated with each other (**Fig. 2B**). The correlation heatmap focusing on these regions showed the emergence of 2 anticorrelated cholinergic subsystems during nicotine withdrawal; one in the basal forebrain consisting of MA, DBN, and SI and one in the brainstem-thalamic area consisting of PPN, MH and LH, with the MS in between. Under control conditions (saline) there was no such organization. The average correlation between the cholinergic regions within the basal forebrain or within the brainstem-thalamic network was significantly higher in the nicotine group compared to the saline group (two-way ANOVA F(1,36)=7.85, p=0.008), with a significant difference between regions (two-way ANOVA F(2,36)=4.62, p=0.016), and a significant interaction (two-way ANOVA F(2,36)=3.36, p=0.046) (**Fig. 2C**)^a^. Post-hoc analysis confirmed that under nicotine withdrawal the average correlation within the subgroups was significantly higher than their interaction (p<0.04).

**Figure 2.**
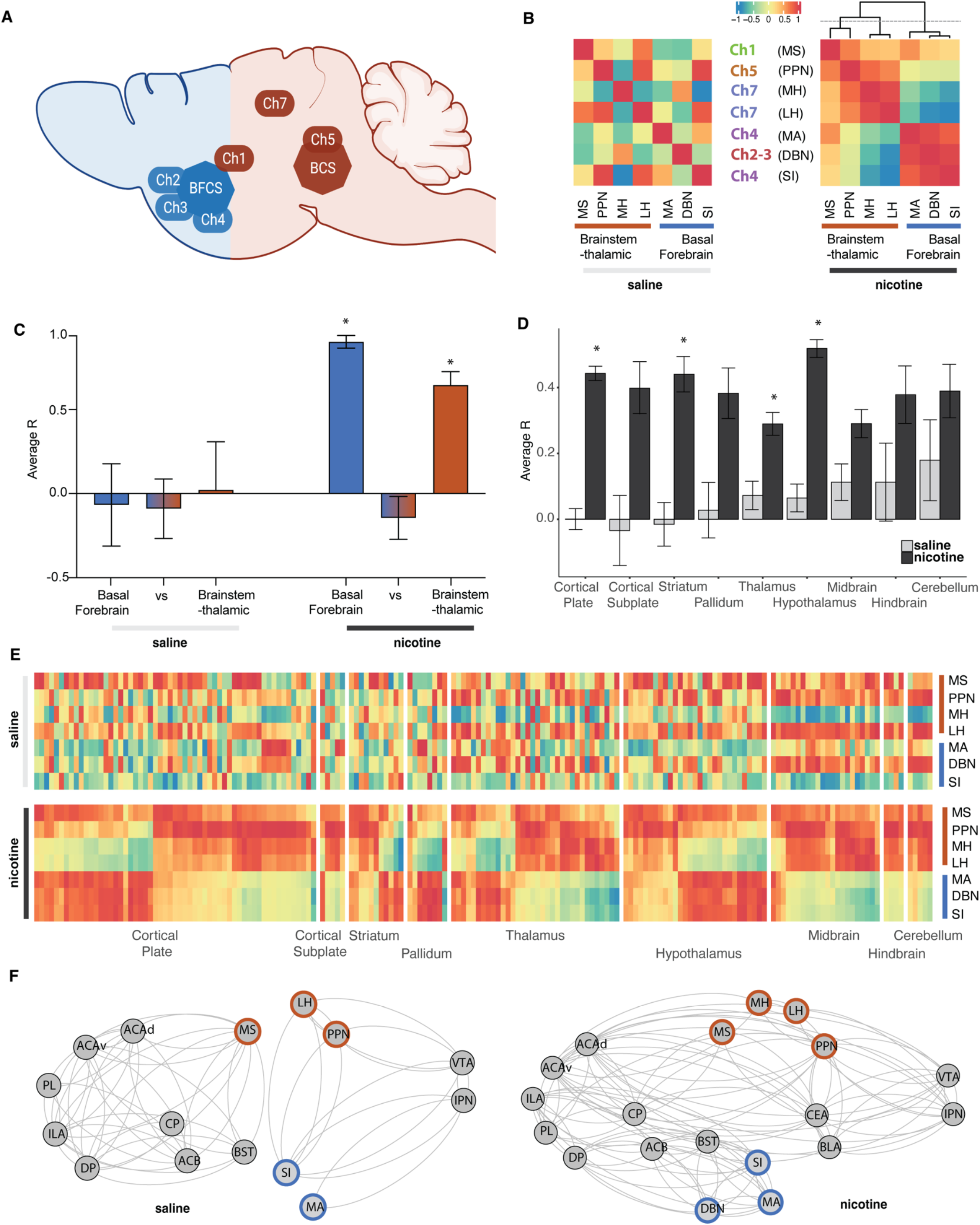
**Interactions amongst the long-range cholinergic groups and with the whole-brain** for saline and nicotine. (**A**) Brain schematic showing the localization and circuitry of the long-range cholinergic groups divided in basal forebrain cholinergic system (BFCS; blue) and brainstem-thalamic cholinergic system (BCS; brown). (**B**) Heatmap representation of the correlation of cholinergic long-range groups. (**C**) Average correlation (R) between the cholinergic long-range groups within the basal forebrain cholinergic system (blue), brainstem-thalamic cholinergic system (brown), or between (*p<0.05)^a^. (**D)** Average correlation (R) of cholinergic long-range groups with the rest of the brain organized into anatomical groups (*p<0.05)^b^. (**E**) Heatmap representation of D with the correlation to all individual regions in the brain (order of the regions see Ext. Fig. 2-1). (**F**) Integration of the long-range cholinergic groups in representative minimal addiction networks. Abbreviations: anterior cingulate area (ACA), infralimbic area (ILA), prelimbic area (PL), dorsal peduncular area (DP), caudoputamen (CP), nucleus accumbens (ACB), bed nucleus of the stria terminalis (BST), basolateral amygdala (BLA), central amygdala (CEA), ventral tegmental area (VTA), and interpeduncular nucleus (IPN).

We then looked at how these long-range cholinergic groups correlated with the different anatomical groups throughout the brain using hierarchical clustering within the main anatomical structures (cortical plate, cortical subplate, striatum, pallidum, thalamus, hypothalamus, midbrain, hindbrain, or cerebellum) (**Fig. 2D-E**). There was a significant increase in correlation throughout the brain in the nicotine group compared to the saline group (two-way ANOVA F(1,2082)=100.7, p<0.0001), without significant difference between regions (two-way ANOVA F(8,2082)=1.32, p=0.23), but with significant interaction (two-way ANOVA F(8,2082)=3.21, p=0.0012)^b^. In line with the overall increase in correlation, the posthoc test showed an increase in the correlation of the cholinergic groups with the cortical plate (p<0.0001), striatum (p<0.0001), thalamus (p=0.0035), and hypothalamus (p<0.0001).

Finally, we looked at the interaction of the long-range cholinergic regions with brain regions that have been shown to be critical to nicotine addiction and nicotine withdrawal including the anterior cingulate area (ACA) (Hong et al., 2009; Wang et al., 2019; Abulseoud et al., 2020), infralimbic area (ILA) (George and Koob, 2010; Huang et al., 2015; Kutlu et al., 2016), prelimbic area (PL) (George and Koob, 2010; Semenova et al., 2018), dorsal peduncular area (DP) (George and Koob, 2010), caudoputamen (CP) (Muskens et al., 2012; Huang et al., 2015), nucleus accumbens (ACB) (Rada et al., 2001; Schmidt et al., 2001; Huang et al., 2015), bed nucleus of the stria terminalis (BST) (Reisiger et al., 2014; Qi et al., 2016), basolateral amygdala (BLA) (Bergstrom et al., 2010; Sharp, 2019) and central amygdala (CEA) (Baiamonte et al., 2014; Huang et al., 2015; Funk et al., 2016), ventral tegmental area (VTA) (Grieder et al., 2014; Huang et al., 2015; Wills and Kenny, 2021), and interpeduncular nucleus (IPN) (Molas et al., 2017; Wills and Kenny, 2021; Klenowski et al., 2022). Here too, the minimal networks showed an overall increased functional connectivity during nicotine withdrawal, particularly between the cortex, subcortical regions, and key cholinergic regions including cholinergic groups 2, 3, 4, and 7 (PPN, MH, LH, SI, MA, and BDN), which were separated in basal forebrain and brainstem-thalamic groups (**Fig. 2F**).

### Non-long-range cholinergic regions function as connector hubs in the nicotine network

To better understand the role of the cholinergic groups within the whole-brain network and validate their role as hub regions or identify others, we calculated the network centrality measures: degree (number of connections a region has) and betweenness (number of shortest paths between two regions that go through a region). Nicotine withdrawal significantly increased the average degree (p < 2.2e-16^c^, **Fig. 3A**) and decreased the average betweenness of the network (p = 2.0e-08^d^, **Fig. 3B**). Regions that were in the top 20 of both degree and betweenness were considered hub regions (Wheeler et al., 2013). For the saline network, 2 hubs with both high degree and betweenness were identified: the hypothalamic parastrial area (PS) and the midbrain cuneiform nucleus (CUN). For the nicotine network, 4 hubs with those criteria were identified: the fundus of the striatum (FS), paraventricular hypothalamic nucleus (PVH), gustatory areas (GU), and posterolateral visual areas (VISpl). The fundus of the striatum and caudoputamen stood out for having significantly higher betweenness scores than other regions and thus having a central role in the network involving most shortest paths during nicotine withdrawal.

**Figure 3.**
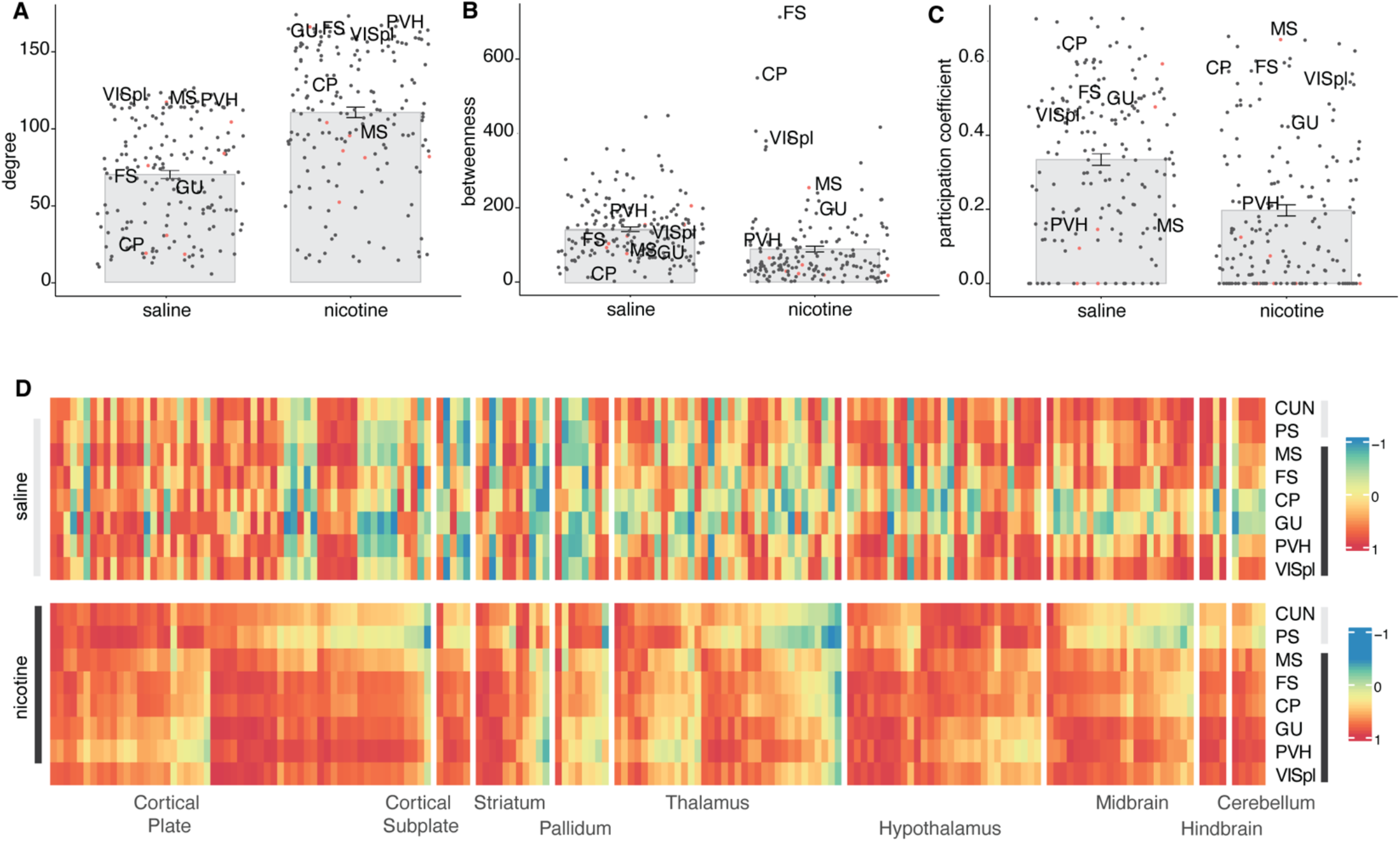
Centrality measurements for hub regions in the saline and nicotine networks. ***(A)** degree (*p < 2.2e-16^c^), *(**B**) betweenness* (p = 2.0e-08^d^)*, and (**C**) participation coefficient (*p =3.5e-9^e^*), with the long-range cholinergic regions identifiable by a red dot red and the hub regions for nicotine withdrawal labeled. (**D**) Heatmap representation of the correlation of the hub regions to all individual regions in the brain (order of the regions same as Ext. Fig. 2-1). Abbreviations: the fundus of the striatum (FS), paraventricular hypothalamic nucleus (PVH), gustatory areas (GU), and posterolateral visual areas (VISpl), caudoputamen (CP), and medial septum (MS)*.

Because the networks are modular, an important role of hub regions is to act as connectors between modules, which is captured through a high participation coefficient that measures the intermodule connectivity or the extent to which a region connects to multiple other modules. Regions with a high participation coefficient were therefore also considered as hubs. Nicotine withdrawal significantly decreased the participation coefficient (p =3.5e-9^e^, **Fig. 3C**). The cholinergic group 1 MS had the highest participation coefficient of the network and thus functions as a top connector between the network modules. The hubs fundus of the striatum and caudoputamen also scored high for this measure. The central role of these regions in the network is confirmed by looking at their correlation with all brain regions (**Fig. 3D**), which showed strongly increased correlation during nicotine withdrawal.

### Identification of novel gene targets that correlates with brain-wide FOS activation

To examine the contribution of the regional expression level of cholinergic-related genes like the nAChRs to the organization of the functional connectome, the basal expression level of *Chrna1-10, Chrnb1-3, Chrnd, and ChAT* throughout the brain was extracted from the in-situ Allen Brain database (Lein et al., 2007) and examined. While *Chrna1, Chrnb1*, and *Chrnd* are generally considered muscle-type subunits, expression in the brain has been observed (Aishah et al., 2017). No clear pattern could be observed differentiating expression in the different modules of the functional connectome (**Fig. 4A** left; expression within the long-range cholinergic groups is highlighted on the right). When organizing the brain regions in anatomical order on the other hand, expression patterns could be observed with *Chrna4* and *Chrnab2* being expressed mostly in the thalamus, *Chrna3, Chrna6*, and *Chrnb3* mostly in the midbrain, and *Chrna1, Chrna2, Chrna7, Chrna9, Chrna10, Chrnb1*, and *Chrnd* mostly in the cortical plate (**Fig. 4A** middle).

**Figure 4.**
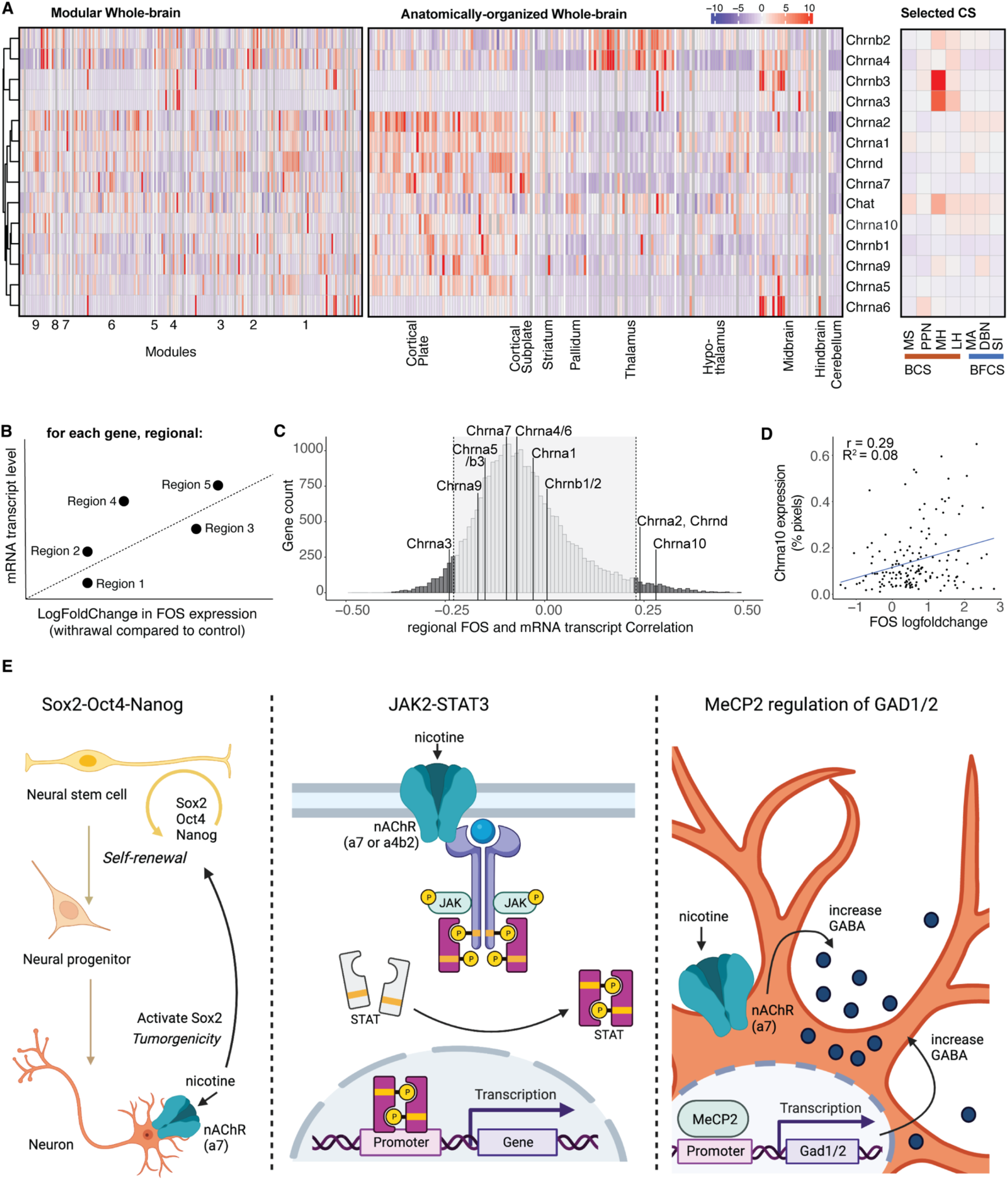
Whole-brain expression distribution of cholinergic receptor subunits, cholinergic transferase, and other proteins. (**A**) Expression density throughout the whole brain hierarchically clustered by row and split into the modules of the nicotine-withdrawal network (left; regions ordered as in Fig. 1C, list in Ext. Fig. 1-1), split into anatomical groups (middle; regions ordered as in Fig. 2E, list in Ext. Fig. 2-1), and the selected long-range cholinergic groups. (**B**) Diagram of the correlation analysis for each gene between basal mRNA expression level and nicotine withdrawal induced FOS transcriptional change in each region. (**C**) Histogram of the number of genes (count) for all found correlations. (**D**) Example correlation graph for the most correlated cholinergic-related gene Chrna10. (**E**) Schematic representations of the pathways identified by the Reactome analysis for the identified significantly correlated genes: Oct4, Sox2 and Nanog activating genes related to proliferation (left), gene expression by JAK-STAT signaling (middle), and the MeCP2 pathway for regulating the transcription for genes involved in GABA signaling through GAD1 and GAD2 (right), with the potential involvement of nicotine.

Next, we wanted to compare the contribution of these cholinergic-related genes to the changes in whole-brain FOS activation and compare it to the other 19,413 genes of which the in-situ Allen Brain database contains the region-specific gene expression (Lein et al., 2007). For every gene, we looked at the correlation between the baseline gene expression level (%-pixel) and the FOS expression change in nicotine withdrawal vs saline control (log-fold change) for every region (Davoudian et al., 2023) (**Fig.4B**). Significance was obtained for genes with a correlation coefficient higher than |0.23| (False Discovery Rate <5%) (**Fig. 4C**), which included *Chrna2, Chrna3, Chrna10* (**Fig. 4D**), and *Chrnd*. The expression of the other cholinergic-related genes did not significantly correlate with the increased induction of FOS expression during nicotine withdrawal. However, we identified 1755 genes that were significantly correlated with FOS expression during withdrawal (False Discovery Rate < 5%, see Ext, Table 4-1).

To investigate the obtained gene list, it was inserted into Reactome, the free, open-source, curated, and peer-reviewed pathway database (Joshi-Tope et al., 2005), which returned 3 top hits: 1) Octamer-binding transcription factor 4 (OCT4), sex determining region Y-box 2 (SOX2), nanog homeobox (NANOG) activate genes related to proliferation (p=5.12E-3), 2) Gene and protein expression by janus kinase (JAK) - signal transducer and activator of transcription (STAT) signaling after Interleukin-12 stimulation (p=1.08-2), and methyl CpG binding protein 2 (MeCP2) regulation of transcription of genes involved in gamma-aminobutyric acid (GABA) signaling (p=6.91-3) (**Fig. 4E**).

## Discussion

This work follows up on the published established whole-brain nicotine withdrawal network obtained through single-cell whole-brain imaging of the immediate early gene *c-Fos* compared to controls (Kimbrough et al., 2021), focusing on the long-range cholinergic regions to help interpret and understand specific functional connectome changes. Contrary to our hypothesis, the well-defined long-range cholinergic groups (Ch1-7) were not found to cluster together, but rather were distributed throughout the nicotine withdrawal network. Cholinergic regions showed increased functional connectivity with all regions of the brain through two anticorrelated sub-networks separated into basal forebrain projecting and brainstem-thalamic projecting cholinergic regions, validating a long-standing hypothesis of the organization of the brain cholinergic systems. Most of the cholinergic-related genes were found to have whole-brain expression profiles that correlated poorly with the nicotine withdrawal-induced FOS changes except for *Chrna2, Chrna3, Chrna10*, and *Chrnd* mRNA. Finally, we identified a list of over 1700 genes for which the baseline expression correlated significantly with the altered brain reactivity in the nicotine withdrawal state and identified cellular pathways that may contribute to neuronal activation during nicotine withdrawal.

This report demonstrates that each module in the nicotine withdrawal network includes at least one of the well-defined long-range cholinergic groups (Ch1-7), and that the localization of each group within each module was consistent with known anatomical and functional connections for these groups (**Fig. 1)**. Cholinergic group 1 is the primary cholinergic input to the hippocampus (Teles-Grilo Ruivo and Mellor, 2013; Muller and Remy, 2018) and was found in the cortico-hypothalamus module. Cholinergic group 5 is the primary cholinergic input for the brainstem (Grofova and Keane, 1991; Mena-Segovia and Bolam, 2017) and was part of the cortico-mid-hind-brain module. Cholinergic groups 2, 3, and 4 are the primary projections to the isocortex, striatum, and amygdala (Mesulam et al., 1983; Luiten et al., 1987; Alheid and Heimer, 1988) and were found in the orbitofrontal-extended amygdalar and cortico-hypothalamic modules. Finally, cholinergic group 7, projects to the brainstem and was found in the midbrain-thalamo-habenular module. Cholinergic neurons have been described to act and project globally rather than modular, which helps in communication throughout the whole brain (Mesulam et al., 1983; Woolf, 1991).

Nicotine withdrawal had strong effects on the functional connectome. First, the functional connectivity was increased between the long-range cholinergic regions and the rest of the brain (**Fig. 2**), particularly with the regions that had lower functional connectivity under control conditions such as the cortical plate, striatum, thalamus, and hypothalamus. The increased synchronization between the long-range cholinergic regions and the rest of the brain may contribute to the synchronization of FOS activity throughout the brain, resulting in decreased modularity (Kimbrough et al., 2021). A possible mechanism underlying this brain-wide synchronization is a global increase in acetylcholine release during withdrawal (Rada et al., 2001; Carcoba et al., 2014) leading to activation of nAChRs, intracellular cation influx, and activation of multiple intracellular cascades activating *c-Fos* transcription (Merlo Pich et al., 1999; Hu et al., 2002; Changeux, 2010; Mizuno et al., 2015). These results are in line with human fMRI data, where increases in resting state connectivity during nicotine withdrawal have also been observed (Fedota and Stein, 2015). Moreover, increased local connectivity within specific network nodes correlate with subjective measures of nicotine craving and measures of nicotine dependence (Claus et al., 2013; Janes et al., 2014; Moran-Santa Maria et al., 2015).

Second, nicotine withdrawal caused a functional reorganization of the long-range cholinergic network composed of the MA, DBN, and SI on one side and the MS, PPN, MH, and LH on the other side, which are correlated within, anticorrelated between, and connect to mostly non-overlapping regions in the brain. Also, when looking at a minimal network containing the long-range cholinergic regions and key regions known to be involved in addiction, the same findings were illustrated: increased functional connectivity between the cholinergic groups 2, 3, 4, and 7 and the anterior cingulate, infralimbic, prelimbic, dorsal peduncular, and ventral tegmental area, caudoputamen, nucleus accumbens, basolateral and central amygdala, bed nucleus of the stria terminalis, and interpeduncular nucleus and the increased subdivision of the circuitry in basal forebrain and brainstem-thalamic cholinergic systems. Indeed, several cholinergic regions (MH, MA, DBN) were not incorporated in this minimal network under control conditions. The organization of a basal forebrain and a brainstem cholinergic system resembles the original anatomical descriptions of the basal forebrain and brainstem cholinergic systems by Woolf and Mesulam (Mesulam et al., 1983; Woolf, 1991; George et al., 2006), which not only validates the approach of single-cell whole-brain imaging for functional connectome analysis, but has profound implications from a theoretical point of view. Indeed, it suggests that the different cholinergic regions throughout the brain are not independent from each other, but instead are functionally connected through two opposite systems, a basal forebrain cholinergic system and a brainstem cholinergic system. Nicotine withdrawal then emerges with the dysregulation of these two systems that become anticorrelated. Whether one cholinergic system inhibits the other or whether they are anticorrelated though the action of a third system remains to be tested.

Third, the central hubs of the network changed. In the saline control network, the cuneiform nucleus and parastrial area acted as hub regions with high degree and betweenness, measures of network centrality (**Fig. 3**) (Wheeler et al., 2013). Following nicotine withdrawal, the fundus of the striatum and caudoputamen were identified as hub regions with high network centrality measures degree and betweenness. The long-range cholinergic region MS (cholinergic group 1), which is the main input of the hippocampus and has been associated with the anxiogenic effects of nicotine (Zarrindast et al., 2013) had a high participation coefficient and therefore also participated as hub, especially in the connection between the different modules. It’s connector role between the basal forebrain and brainstem cholinergic systems was already illustrated in Fig. 2.B. While the fundus of the striatum is typically not recognized as a long-range cholinergic region, it is a transition zone between the ventral part of the caudoputamen and the substantia innominate (cholinergic group 4) that expresses high levels of acetylcholine esterase and where dopamine release is under a particular tight cholinergic control (O’Connor et al., 1995). The caudoputamen is the brain region with the highest basal acetylcholine level due to a dense cholinergic arborization originating from cholinergic interneurons (Zhou et al., 2002; Gonzales and Smith, 2015; Abudukeyoumu et al., 2019). Caudoputamen cholinergic interneurons are critical to dopamine release, reinforcement learning and the formation of habit (Knowlton et al., 1996; Matsumoto et al., 1999; Kitabatake et al., 2003). These hub regions all play central roles in orchestrating the negative emotional state under nicotine withdrawal and were all found together in the intermediate size module 3 (Ext. Fig. 1-1 and Ext. Fig. 1-3). The medial septum - fundus of the striatum - caudoputamen module might thus function as the main nicotine responsive module that orchestrates the whole-brain response.

All nicotinic receptor genes, except the muscle-type CHRNB1, including eight genomic regions containing 11 neuronal CHRN genes and three genomic regions containing four muscle-type CHRN genes, have been significantly associated with nicotine dependence and/or alcohol dependence (Zuo et al., 2016). Analysis of the correlation between baseline mRNA expression of the nAChRs in all brain regions with the withdrawal-induced change in FOS expression (**Fig. 4B**), showed significant correlation for *Chrna2, Chrna3, Chrna10*, and *Chrnd* (**Fig. 4C-D**). *Chrna2* has been identified in human GWAS studies in association with smoking related behaviors, like smoking status, smoking initiation, cigarettes smoked per day, and smoking cessation (Liu et al., 2019; Xu et al., 2020). *Chrna3* is part of a locus on chromosome 15q25 with *Chrna5* and *Chrnb4*, which has also been identified in human GWAS studies to be associated with smoking related behaviors and nicotine dependence. One SNP is specifically localized in *Chrna3* (Spitz et al., 2008; Liu et al., 2010). *Chrna10* was identified through linkage analysis in sibling pairs for nicotine withdrawal (Pergadia et al., 2009) and found together with *Chrnd* to increase the risk for nicotine dependence in an African-American population subset (Saccone et al., 2010). It is important to note that these correlations were obtained using baseline gene expression with no exposure to nicotine, suggesting that these genes may be predisposing factors to nicotine dependence. However, further studies are required to examine the correlation between withdrawal-induced gene expression and withdrawal-induced FOS activity. Looking at the mRNA expression was a first attempt to link nAChRs levels to Fos activation. A limitation is that mRNA expression does not necessarily correlate with protein levels or even functional activity of the protein, therefore, while we observed significant correlations between some nAChRs and differential Fos expression, it is possible that negative results for other subunits may be due to a lack of correlation between mRNA levels and protein levels for instance due to posttranscriptional events (Mousavi et al., 2003). Finally, another limitation is that both the original whole-brain reactivity (Kimbrough et al., 2021) and whole-brain gene expression (Lein et al., 2007) studies only incorporated male subjects in their study and database precluding any analysis of sex differences. Follow up studies are needed to evaluate if these effects also exist in females.

We then extended the gene analysis to include the mRNA transcript levels of all 19,413 genes of the In Situ Hybridization gene expression Allen Brain Atlas database, which resulted in a list of 1755 genes that had significant correlations between their expression and the nicotine withdrawal-induced FOS changes. Through pathway analysis (Joshi-Tope et al., 2005), we identified potentially promising gene and pathways that may contribute to the FOS expression during nicotine withdrawal. The first pathway contained the transcription factors *Sox2, Oct4*, and *Nanog*, which are highly expressed in proliferative adult neurogenesis precursor cells in discrete brain regions (Suh et al., 2007; Bennett et al., 2009; Ahlfeld et al., 2017; Stevanovic et al., 2021) that are associated with FOS expression (Velazquez et al., 2015) and have been reported to be affected by nicotine (Brooks and Henderson, 2021). The second pathway was part of the immune response by JAK2-STAT3, which has been shown to be activated by nicotine through complex formation with Chrna7 or Crna4/Chrnab2 to provide neuroprotective effects (Shaw et al., 2002; Wang et al., 2022). The third pathway brings up another very relevant transcription factor, MeCP2, which is mutated in the neurodevelopmental disorder Rett syndrome (Amir et al., 1999). MeCP2 knock-out mice have reduced ChAT-positive cells, reduced Chrna4 and Chrna6 expression, an attenuated behavior response to nicotine (Leung et al., 2017), dysfunctional reduced GABA signaling (Chao et al., 2010), and some reversed deficits following nicotine administration (Zhang et al., 2016). MeCP2 has been shown to modulate the effects of drugs of abuse in preclinical models (Deng et al., 2010; Im et al., 2010; Repunte-Canonigo et al., 2014; Xu et al., 2021). GABA signaling, regulated by MeCP2 through GAD1 and GAD2 that were both in the gene list, has a well-established role in nicotine dependence and withdrawal (Markou, 2008; D’Souza and Markou, 2013; Klenowski et al., 2022). Despite relatively low significance for these pathways (loss of significance with correction for multiple comparisons, False Discovery Rate 80%), the top 3 identified pathways have been shown to be affected by or associated with nicotine, providing validity to this exploratory approach, which might be a way to further process and investigate the obtained whole-brain functional connectome datasets. Finally, these results demonstrate the power of using single-cell whole-brain imaging combined with whole-brain transcriptomics to identify new brain regions, new gene targets, and new cellular pathways that may contribute to nicotine dependence and substance use disorder in general.

In conclusion, these results demonstrate that cholinergic regions increased functional connectivity with the rest of the brain through two anticorrelated sub-networks separated into basal forebrain projecting and brainstem-thalamic projecting cholinergic regions. The expression level of *Chrna2, Chrna3, Chrna10*, and *Chrnd* mRNA throughout the brain was correlated with the nicotine withdrawal-induced FOS changes. Finally, we’ve identified a list of over ~1700 genes for which the baseline expression correlates significantly with the altered brain reactivity in the nicotine withdrawal state and identified cellular pathways that may contribute to neuronal activation during nicotine withdrawal.

## Author contributions

LLGC and OG analyzed the dataset and wrote the manuscript. AC collected the microscopy images. AK analyzed the microscopy images to generate the Fos counts dataset. PAD and ACK contributed code for analyzing the ISH data. All authors read and approved the submitted version.

## Acknowledgements

Support from the Preclinical Addiction Research Consortium is acknowledged. This work was supported by funding from the National Institute on Drug Abuse 1R21DA057694, the Tobacco-Related Disease Research Program Grants 27IR0047 and 32IR5384 to OG, and an innovation Award from the International Rett Syndrome Foundation 3901 to LC. Publication fees were contributed by the UCSD library. The authors declare no competing financial interests. Current address for AK: Department of Basic Medical Sciences, College of Veterinary Medicine, Purdue University, West Lafayette, IN, 47906, United States

