## Extended data for "Hyperconnectivity of two separate long-range cholinergic systems contributes to the reorganization of the brain functional connectivity during nicotine withdrawal in male mice"

**saline**

| Region | Module |
|---|---|
| Posterior limiting nucleus of the thalamus | 13 |
| Endopiriform nucleus | |
| Olfactory tubercle | |
| Central amygdalar nucleus | |
| Basolateral amygdalar nucleus | |
| Globus pallidus external segment | 12 |
| Posterior complex of the thalamus | |
| Ventral anterior–lateral complex of the thalamus | |
| Piriform–amygdalar area | 11 |
| Diagonal band nucleus | |
| Ventrolateral preoptic nucleus | |
| Cortical amygdalar area posterior part | |
| Anteroventral preoptic nucleus | |
| Nucleus of the lateral olfactory tract | |
| Bed nucleus of the accessory olfactory tract | |
| Parataenial nucleus | |
| Interanteromedial nucleus of the thalamus | |
| Magnocellular nucleus | 10 |
| Globus pallidus internal segment | |
| Temporal association areas | |
| Ventral auditory area | |
| Posterior auditory area | |
| Lateral amygdalar nucleus | |
| Dorsal part of the lateral geniculate complex | |
| Primary auditory area | |
| Postpiriform transition area | |
| Mammillary body | |
| Ventral posterolateral nucleus of the thalamus | |
| Field CA2 | |
| Dorsal auditory area | |
| Entorhinal area lateral part | |
| Ansiform lobule | |
| Entorhinal area medial part | |
| Intergeniculate leaflet of the lateral geniculate complex | |
| Ectorhinal area | 9 |
| Perirhinal area | |
| Reticular nucleus of the thalamus | |
| Induseum griseum | |
| Interpeduncular nucleus | |
| Ventral posterior complex of the thalamus | |
| Red nucleus | |
| Preparasubthalamic nucleus | |
| Nucleus of the brachium of the inferior colliculus | |
| Medial geniculate complex | 8 |
| Ventral tegmental area | |
| Substantia innominata | |
| Medial amygdalar nucleus | |
| Pons | |
| Retrochiasmatic area | |
| Field CA3 | |
| Ventral part of the lateral geniculate complex | |
| Substantia nigra reticular part | |
| Zona incerta | |
| Paraventricular nucleus of the thalamus | 7 |
| Medial habenula | |
| Frontal pole cerebral cortex | |
| Supplemental somatosensory area | |
| Anterior olfactory nucleus | |
| Central medial nucleus of the thalamus | |
| Intercalated amygdalar nucleus | |
| Medial preoptic area | |
| Intermediodorsal nucleus of the thalamus | |
| Visceral area | 6 |
| Caudoputamen | |
| Orbital area ventrolateral part | |
| Anterior amygdalar area | |
| Primary somatosensory area nose | 5 |
| Primary somatosensory area mouth | |
| Claustrum | |
| Anteroventral nucleus of thalamus | |
| Supraoptic nucleus | |
| Gustatory areas | |
| Taenia tecta | |
| Prelimbic area | |
| Piriform area | |
| Agranular insular area ventral part | |
| Dorsal peduncular area | |
| Primary somatosensory area upper limb | |
| Supramammillary nucleus | |
| Anterior cingulate area dorsal part | |
| Orbital area medial part | |
| Infralimbic area | |
| Median preoptic nucleus | |
| Lateral preoptic area | |
| Periventricular hypothalamic nucleus preoptic part | |
| Paraventricular hypothalamic nucleus descending division | 4 |
| Medial pretectal area | |
| Anterior group of the dorsal thalamus | |
| Dorsomedial nucleus of the hypothalamus | |
| Nucleus accumbens | |
| Triangular nucleus of septum | 3 |
| Lateral hypothalamic area | |
| Tuberal nucleus | |
| Posterior amygdalar nucleus | |
| Periventricular hypothalamic nucleus posterior part | |
| Ventral medial nucleus of the thalamus | |
| Subthalamic nucleus | |
| Arcuate hypothalamic nucleus | |
| Ventromedial hypothalamic nucleus | |
| Posterior hypothalamic nucleus | |
| Pedunculopontine nucleus | |
| Culmen | |
| Paracentral nucleus | |
| Substantia nigra compact part | |
| Anterior pretectal nucleus | |
| Nucleus of Darkschewitsch | |
| Field CA1 | |
| Nucleus of the posterior commissure | |
| Submedial nucleus of the thalamus | |
| Subparafascicular nucleus | |
| Simple lobule | |
| Paraflocculus | |
| Thalamus sensory–motor cortex related | |
| Subiculum | |
| Midbrain reticular nucleus | |
| Central lobule | |
| Parafascicular nucleus | |
| Peripeduncular nucleus | |
| Nucleus of reuniens | |
| Posterior pretectal nucleus | |
| Lateral habenula | |
| Nucleus of the optic tract | 2 |
| Postsubiculum | |
| Parasubiculum | |
| Lateral posterior nucleus of the thalamus | |
| Midbrain reticular nucleus retrorubral area | |
| Presubiculum | |
| Dentate gyrus | |
| Fundus of striatum | |
| Cuneiform nucleus | |
| Inferior colliculus | |
| Superior colliculus motor related | |
| Vestibular nuclei | |
| Superior colliculus sensory related | |
| Parasubthalamic nucleus | |
| Primary visual area | |
| Pons motor related | |
| Precommissural nucleus | |
| Retrosplenial area ventral part | |
| Interanterodorsal nucleus of the thalamus | |
| Anterior hypothalamic nucleus | |
| Posterolateral visual area | |
| Primary somatosensory area barrel field | 1 |
| Agranular insular area posterior part | |
| Anteromedial visual area | |
| Lateral dorsal nucleus of thalamus | |
| Primary somatosensory area trunk | |
| Lateral septal complex | |
| Primary somatosensory area lower limb | |
| Primary motor area | |
| Medial preoptic nucleus | |
| Anterodorsal preoptic nucleus | |
| Fasciola cinerea | |
| Anterodorsal nucleus | |
| Periaqueductal gray | |
| Parastrial nucleus | |
| Pontine reticular nucleus | |
| Lateral visual area | |
| Anterolateral visual area | |
| Retrosplenial area lateral agranular part | |
| Bed nuclei of the stria terminalis | |
| Anteroventral periventricular nucleus | |
| Posteromedial visual area | |
| Retrosplenial area dorsal part | |
| Mediodorsal nucleus of thalamus | |
| Orbital area lateral part | |
| Paraventricular hypothalamic nucleus | |
| Septofimbrial nucleus | |
| Periventricular zone | |
| Secondary motor area | |
| Olivary pretectal nucleus | |
| Central lateral nucleus of the thalamus | |
| Medial septal nucleus | |
| Subparaventricular zone | |
| Anterior cingulate area ventral part | |

**nicotine**

| Region | Module |
|---|---|
| Bed nucleus of the accessory olfactory tract | 9 |
| Frontal pole cerebral cortex | |
| Ventral posterior complex of the thalamus | |
| Medial amygdalar nucleus | |
| Bed nuclei of the stria terminalis | |
| Diagonal band nucleus | |
| Substantia innominata | |
| Anterior olfactory nucleus | |
| Medial preoptic nucleus | |
| Primary somatosensory area barrel field | |
| Anterior hypothalamic nucleus | |
| Posterior complex of the thalamus | |
| Ventral anterior–lateral complex of the thalamus | |
| Orbital area lateral part | |
| Lateral dorsal nucleus of thalamus | |
| Orbital area ventrolateral part | |
| Lateral posterior nucleus of the thalamus | 8 |
| Nucleus of the lateral olfactory tract | |
| Anterior group of the dorsal thalamus | |
| Primary somatosensory area mouth | |
| Supraoptic nucleus | 7 |
| Dorsal part of the lateral geniculate complex | |
| Olfactory tubercle | |
| Orbital area medial part | |
| Globus pallidus external segment | |
| Preparasubthalamic nucleus | 6 |
| Primary somatosensory area nose | |
| Taenia tecta | |
| Anterior amygdalar area | |
| Subparaventricular zone | |
| Secondary motor area | |
| Submedial nucleus of the thalamus | |
| Primary somatosensory area trunk | |
| Substantia nigra reticular part | |
| Paracentral nucleus | |
| Subthalamic nucleus | |
| Entorhinal area medial part | |
| Parasubiculum | |
| Cuneiform nucleus | |
| Posterior amygdalar nucleus | |
| Tuberal nucleus | |
| Median preoptic nucleus | |
| Reticular nucleus of the thalamus | |
| Periventricular hypothalamic nucleus preoptic part | |
| Anteroventral preoptic nucleus | |
| Field CA1 | |
| Field CA2 | |
| Agranular insular area ventral part | |
| Retrochiasmatic area | |
| Infralimbic area | |
| Anteroventral periventricular nucleus | |
| Lateral hypothalamic area | |
| Ventral medial nucleus of the thalamus | |
| Globus pallidus internal segment | |
| Magnocellular nucleus | |
| Nucleus accumbens | |
| Medial preoptic area | |
| Lateral preoptic area | |
| Primary somatosensory area upper limb | |
| Dorsal peduncular area | |
| Primary motor area | |
| Zona incerta | |
| Field CA3 | |
| Ventromedial hypothalamic nucleus | |
| Anterodorsal preoptic nucleus | |
| Primary somatosensory area lower limb | |
| Parastrial nucleus | |
| Primary visual area | |
| Interanterodorsal nucleus of the thalamus | 5 |
| Paraventricular nucleus of the thalamus | 4 |
| Parataenial nucleus | |
| Induseum griseum | |
| Triangular nucleus of septum | |
| Lateral habenula | |
| Intergeniculate leaflet of the lateral geniculate complex | |
| Precommissural nucleus | |
| Nucleus of the posterior commissure | |
| Olivary pretectal nucleus | |
| Nucleus of the optic tract | |
| Medial habenula | |
| Medial pretectal area | |
| Interanteromedial nucleus of the thalamus | |
| Thalamus sensory–motor cortex related | |
| Septofimbrial nucleus | |
| Medial septal nucleus | 3 |
| Dentate gyrus | |
| Posterolateral visual area | |
| Anteromedial visual area | |
| Substantia nigra compact part | |
| Paraventricular hypothalamic nucleus descending division | |
| Nucleus of reuniens | |
| Dorsomedial nucleus of the hypothalamus | |
| Subiculum | |
| Parasubthalamic nucleus | |
| Fundus of striatum | |
| Caudoputamen | |
| Arcuate hypothalamic nucleus | |
| Posterior limiting nucleus of the thalamus | |
| Periventricular zone | |
| Paraflocculus | |
| Anterior cingulate area dorsal part | |
| Paraventricular hypothalamic nucleus | |
| Supplemental somatosensory area | |
| Periventricular hypothalamic nucleus posterior part | |
| Ansiform lobule | |
| Presubiculum | |
| Posteromedial visual area | |
| Intercalated amygdalar nucleus | |
| Central amygdalar nucleus | |
| Mammillary body | |
| Gustatory areas | |
| Postsubiculum | |
| Anterolateral visual area | |
| Supramammillary nucleus | |
| Lateral visual area | |
| Anterodorsal nucleus | 2 |
| Posterior pretectal nucleus | |
| Nucleus of Darkschewitsch | |
| Central lateral nucleus of the thalamus | |
| Anterior pretectal nucleus | |
| Parafascicular nucleus | |
| Ventral posterolateral nucleus of the thalamus | 1 |
| Piriform–amygdalar area | |
| Ventrolateral preoptic nucleus | |
| Lateral septal complex | |
| Postpiriform transition area | |
| Dorsal auditory area | |
| Entorhinal area lateral part | |
| Endopiriform nucleus | |
| Cortical amygdalar area posterior part | |
| Central medial nucleus of the thalamus | |
| Prelimbic area | |
| Intermediodorsal nucleus of the thalamus | |
| Agranular insular area posterior part | |
| Piriform area | |
| Ectorhinal area | |
| Perirhinal area | |
| Primary auditory area | |
| Ventral auditory area | |
| Visceral area | |
| Posterior auditory area | |
| Temporal association areas | |
| Anteroventral nucleus of thalamus | |
| Vestibular nuclei | |
| Peripeduncular nucleus | |
| Simple lobule | |
| Fasciola cinerea | |
| Inferior colliculus | |
| Periaqueductal gray | |
| Culmen | |
| Ventral part of the lateral geniculate complex | |
| Mediodorsal nucleus of thalamus | |
| Nucleus of the brachium of the inferior colliculus | |
| Retrosplenial area ventral part | |
| Lateral amygdalar nucleus | |
| Retrosplenial area dorsal part | |
| Interpeduncular nucleus | |
| Red nucleus | |
| Central lobule | |
| Retrosplenial area lateral agranular part | |
| Midbrain reticular nucleus retrorubral area | |
| Ventral tegmental area | |
| Midbrain reticular nucleus | |
| Superior colliculus motor related | |
| Basolateral amygdalar nucleus | |
| Pons motor related | |
| Anterior cingulate area ventral part | |
| Pedunculopontine nucleus | |
| Subparafascicular nucleus | |
| Claustrum | |
| Pons | |
| Superior colliculus sensory related | |
| Posterior hypothalamic nucleus | |
| Pontine reticular nucleus | |
| Medial geniculate complex | |

Figure 1-1: Organization of the regions in the clustered correlation matrices for the saline and nicotine groups (Fig. 1B and C)

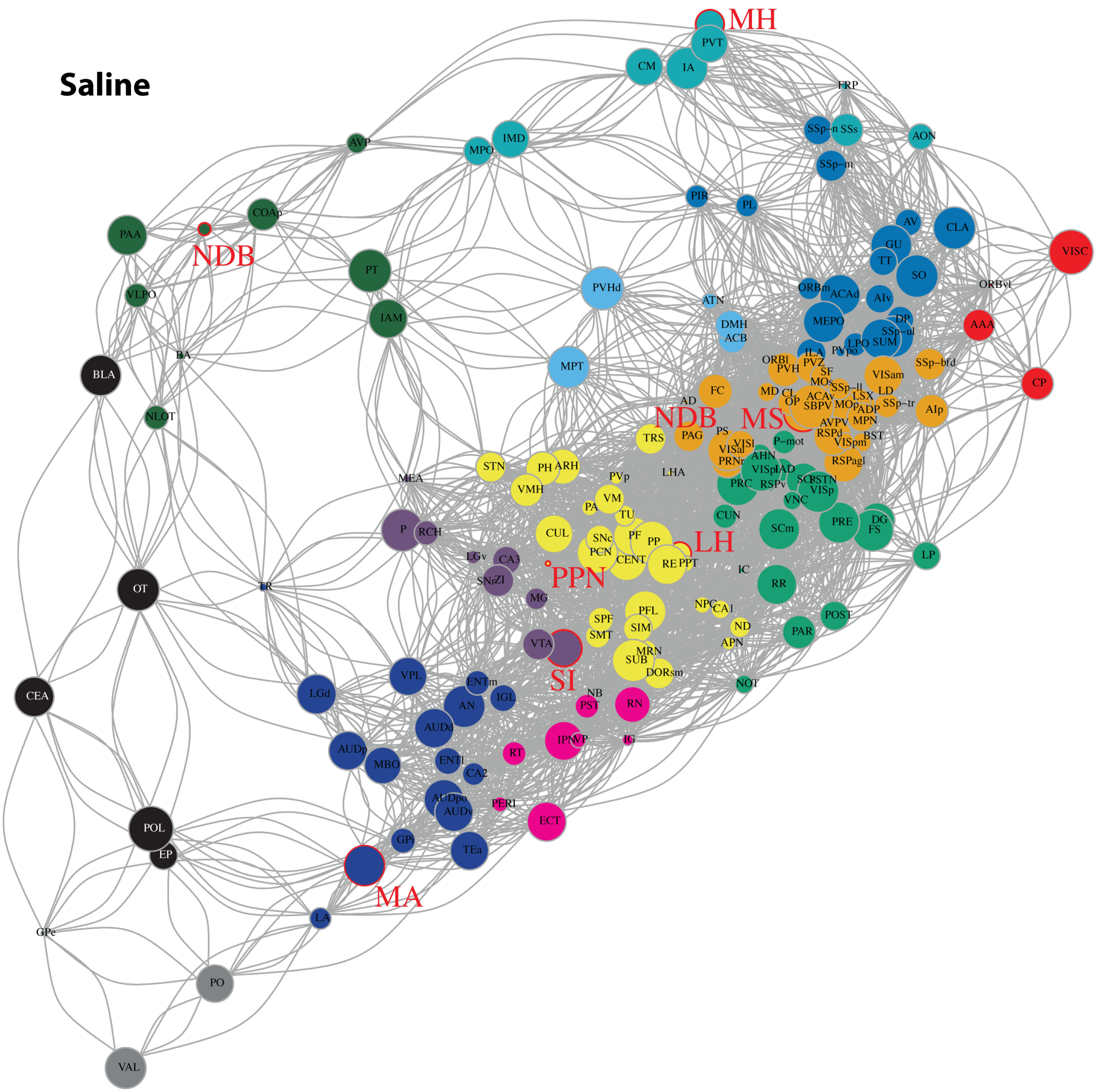


Figure 1-2: zoom and node label of Fig 1D

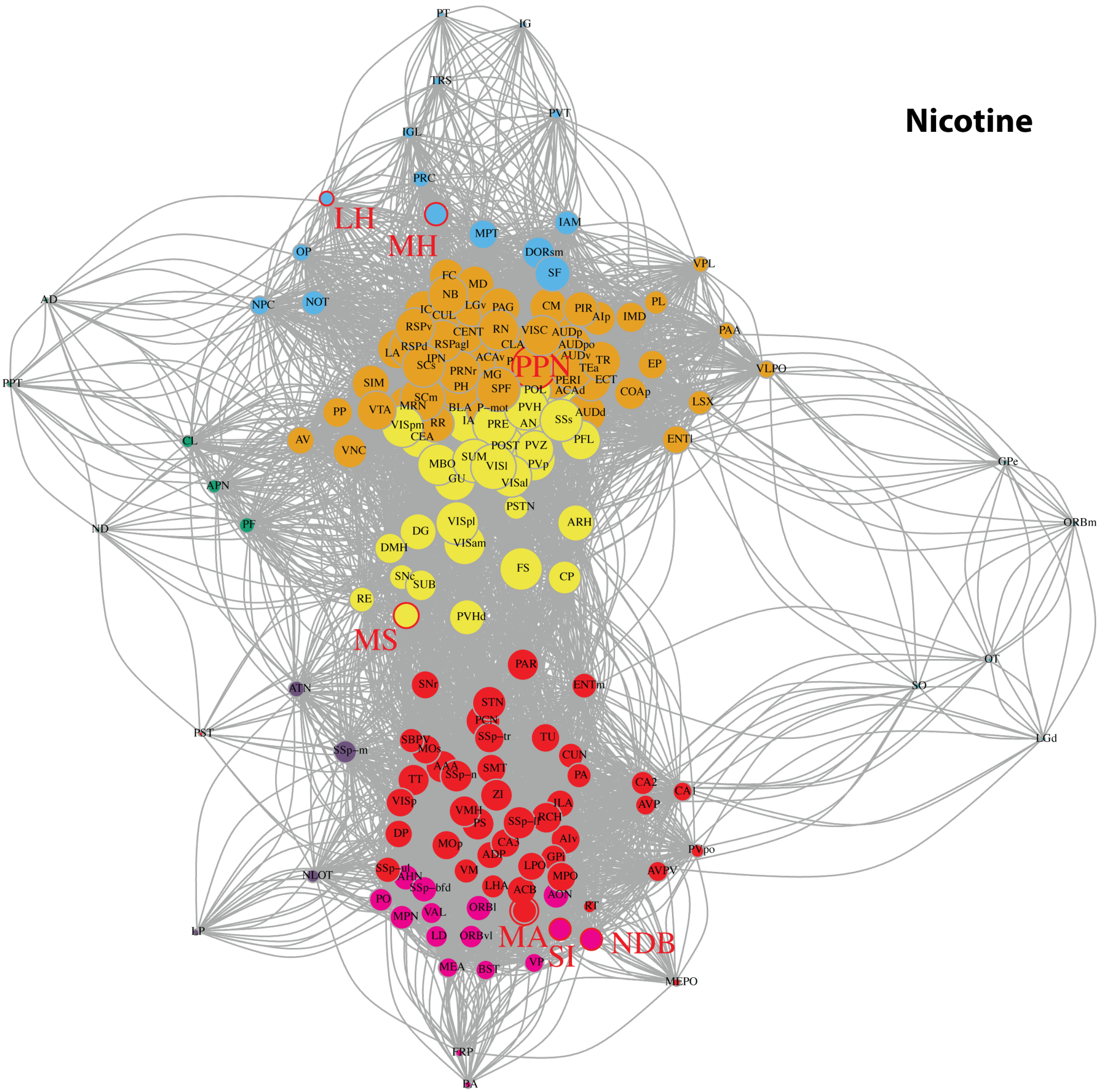


Figure 1-3: zoom and node labels of Fig 1E

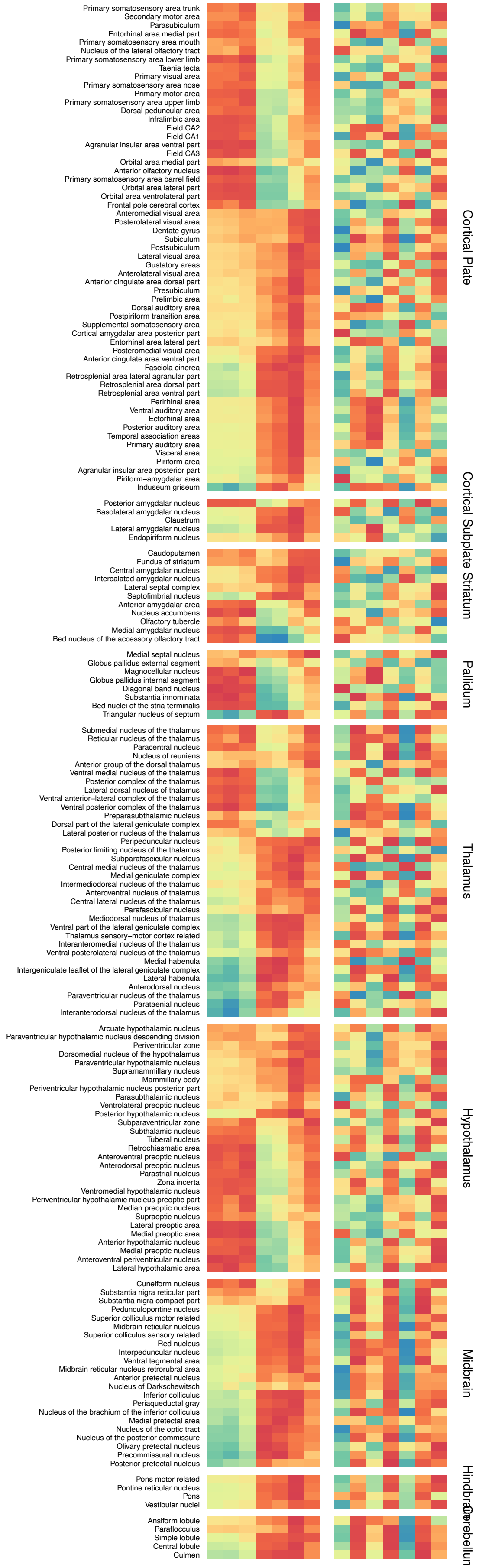


Figure 2-1: Organization of the regions in the correlation matrix separated by anatomical groups (Fig. 2E and 3D)

Table 1-1: Published Fos counts during nicotine withdrawal and saline controls

| Region | Abb | Sal1 | Sal2 | Sal3 | Sal4 | Nic1 | Nic2 | Nic3 | Nic4 | Nic5 |
|---|---|---|---|---|---|---|---|---|---|---|
| Agranular insular area posterior part | AIp | 496 | 1083 | 1364 | 991 | 1195 | 278 | 549 | 1191 | 991 |
| Agranular insular area ventral part | AIv | 155 | 502 | 408 | 221 | 284 | 494 | 306 | 738 | 550 |
| Ansiform lobule | AN | 47 | 16 | 17 | 75 | 20 | 0 | 17 | 381 | 95 |
| Anterior amygdalar area | AAA | 66 | 88 | 102 | 79 | 92 | 154 | 154 | 286 | 181 |
| Anterior cingulate area dorsal part | ACAd | 294 | 2954 | 1008 | 506 | 798 | 176 | 553 | 2649 | 2315 |
| Anterior cingulate area ventral part | ACAv | 210 | 1947 | 816 | 1159 | 1040 | 95 | 1177 | 3819 | 4540 |
| Anterior group of the dorsal thalamus | ATN | 12 | 23 | 11 | 14 | 3 | 13 | 66 | 69 | 31 |
| Anterior hypothalamic nucleus | AHN | 100 | 407 | 187 | 445 | 53 | 1231 | 500 | 3262 | 544 |
| Anterior olfactory nucleus | AON | 96 | 308 | 587 | 47 | 212 | 892 | 245 | 1427 | 546 |
| Anterior pretectal nucleus | APN | 54 | 83 | 131 | 361 | 44 | 39 | 582 | 371 | 143 |
| Anterodorsal nucleus | AD | 19 | 126 | 17 | 64 | 13 | 4 | 213 | 25 | 25 |
| Anterodorsal preoptic nucleus | ADP | 1 | 12 | 7 | 7 | 8 | 17 | 13 | 24 | 23 |
| Anterolateral visual area | VISal | 38 | 215 | 57 | 210 | 94 | 33 | 99 | 479 | 157 |
| Anteromedial visual area | VISam | 51 | 1388 | 795 | 445 | 290 | 206 | 776 | 4010 | 1444 |
| Anteroventral nucleus of thalamus | AV | 36 | 64 | 60 | 37 | 36 | 23 | 160 | 146 | 91 |
| Anteroventral periventricular nucleus | AVPV | 0 | 46 | 12 | 22 | 20 | 120 | 9 | 284 | 113 |
| Anteroventral preoptic nucleus | AVP | 51 | 22 | 16 | 3 | 18 | 51 | 14 | 153 | 148 |
| Arcuate hypothalamic nucleus | ARH | 44 | 137 | 9 | 131 | 44 | 17 | 27 | 668 | 391 |
| Basolateral amygdalar nucleus | BLA | 700 | 354 | 458 | 266 | 393 | 130 | 403 | 771 | 481 |
| Bed nuclei of the stria terminalis | BST | 125 | 374 | 325 | 344 | 154 | 3701 | 449 | 3124 | 837 |
| Bed nucleus of the accessory olfactory tract | BA | 40 | 13 | 6 | 10 | 7 | 96 | 9 | 42 | 9 |
| Caudoputamen | CP | 5517 | 8131 | 12885 | 7919 | 10055 | 9691 | 10145 | 15966 | 15386 |
| Central amygdalar nucleus | CEA | 565 | 301 | 485 | 291 | 369 | 261 | 475 | 650 | 582 |
| Central lateral nucleus of the thalamus | CL | 7 | 19 | 11 | 14 | 8 | 5 | 217 | 78 | 44 |
| Central lobule | CENT | 74 | 129 | 52 | 340 | 115 | 0 | 111 | 494 | 230 |
| Central medial nucleus of the thalamus | CM | 21 | 29 | 19 | 9 | 16 | 0 | 5 | 22 | 86 |
| Claustrum | CLA | 255 | 568 | 654 | 271 | 454 | 142 | 402 | 683 | 618 |
| Cortical amygdalar area posterior part | COAp | 245 | 159 | 54 | 49 | 244 | 45 | 83 | 501 | 195 |
| Culmen | CUL | 78 | 180 | 16 | 518 | 95 | 0 | 116 | 176 | 290 |
| Cuneiform nucleus | CUN | 2 | 13 | 5 | 27 | 3 | 9 | 6 | 31 | 32 |
| Dentate gyrus | DG | 1305 | 2285 | 3007 | 3272 | 1420 | 1275 | 3347 | 6816 | 3709 |
| Diagonal band nucleus | NDB | 273 | 100 | 41 | 29 | 138 | 658 | 117 | 734 | 475 |
| Dorsal auditory area | AUDd | 2095 | 1282 | 1423 | 2322 | 2239 | 534 | 1004 | 5255 | 3344 |
| Dorsal part of the lateral geniculate complex | LGd | 116 | 9 | 15 | 62 | 89 | 78 | 12 | 146 | 76 |
| Dorsal peduncular area | DP | 16 | 169 | 123 | 42 | 19 | 115 | 89 | 388 | 79 |
| Dorsal premammillary nucleus | PMd | 34 | 10 | 1 | 7 | 0 | 0 | 2 | 149 | 9 |
| Dorsomedial nucleus of the hypothalamus | DMH | 116 | 454 | 118 | 184 | 33 | 31 | 285 | 859 | 213 |
| Ectorhinal area | ECT | 907 | 532 | 1188 | 1734 | 2011 | 262 | 796 | 3920 | 2407 |
| Endopiriform nucleus | EP | 811 | 657 | 793 | 735 | 1240 | 411 | 541 | 1623 | 1150 |
| Entorhinal area lateral part | ENTl | 1009 | 329 | 449 | 1367 | 1110 | 172 | 206 | 2762 | 1061 |
| Entorhinal area medial part | ENTm | 37 | 11 | 9 | 88 | 29 | 34 | 19 | 308 | 84 |

| | | | | | | | | | | |
|---|---|---|---|---|---|---|---|---|---|---|
| Fasciola cinerea | FC | 5 | 83 | 6 | 28 | 15 | 0 | 24 | 23 | 93 |
| Field CA1 | CA1 | 1505 | 1937 | 1992 | 3435 | 2148 | 3146 | 1295 | 8391 | 3982 |
| Field CA2 | CA2 | 323 | 218 | 257 | 363 | 310 | 421 | 254 | 780 | 508 |
| Field CA3 | CA3 | 750 | 733 | 587 | 842 | 792 | 1703 | 1111 | 3420 | 1557 |
| Frontal pole cerebral cortex | FRP | 6 | 13 | 13 | 2 | 4 | 1081 | 86 | 151 | 457 |
| Fundus of striatum | FS | 16 | 45 | 73 | 84 | 40 | 33 | 51 | 211 | 115 |
| Globus pallidus external segment | GPe | 139 | 56 | 282 | 72 | 259 | 104 | 42 | 353 | 144 |
| Globus pallidus internal segment | GPi | 28 | 6 | 25 | 36 | 29 | 101 | 31 | 330 | 66 |
| Gustatory areas | GU | 426 | 1312 | 1030 | 471 | 651 | 501 | 858 | 1345 | 978 |
| Induseum griseum | IG | 6 | 3 | 11 | 43 | 47 | 0 | 11 | 1 | 55 |
| Inferior colliculus | IC | 119 | 502 | 398 | 2183 | 234 | 3 | 482 | 663 | 1742 |
| Infralimbic area | ILA | 121 | 1211 | 362 | 385 | 571 | 812 | 545 | 1451 | 965 |
| Interanterodorsal nucleus of the thalamus | IAD | 6 | 21 | 12 | 25 | 7 | 3 | 38 | 0 | 15 |
| Interanteromedial nucleus of the thalamus | IAM | 18 | 20 | 2 | 7 | 9 | 0 | 3 | 4 | 24 |
| Intercalated amygdalar nucleus | IA | 66 | 89 | 59 | 41 | 51 | 25 | 73 | 130 | 109 |
| Intergeniculate leaflet of the lateral geniculate complex | IGL | 41 | 23 | 22 | 64 | 22 | 0 | 33 | 4 | 88 |
| Intermediodorsal nucleus of the thalamus | IMD | 9 | 12 | 6 | 5 | 9 | 0 | 1 | 12 | 33 |
| Interpeduncular nucleus | IPN | 67 | 35 | 76 | 265 | 42 | 0 | 71 | 233 | 50 |
| Lateral amygdalar nucleus | LA | 594 | 328 | 702 | 649 | 747 | 213 | 946 | 1156 | 944 |
| Lateral dorsal nucleus of thalamus | LD | 59 | 97 | 91 | 85 | 48 | 537 | 160 | 830 | 177 |
| Lateral habenula | LH | 46 | 105 | 51 | 256 | 42 | 0 | 247 | 19 | 79 |
| Lateral hypothalamic area | LHA | 608 | 1240 | 556 | 1371 | 977 | 2807 | 923 | 6692 | 1552 |
| Lateral posterior nucleus of the thalamus | LP | 59 | 101 | 202 | 171 | 47 | 359 | 630 | 422 | 187 |
| Lateral preoptic area | LPO | 62 | 312 | 172 | 140 | 117 | 771 | 221 | 1717 | 745 |
| Lateral septal complex | LSX | 260 | 798 | 623 | 628 | 1303 | 584 | 607 | 1796 | 2138 |
| Lateral visual area | VISl | 64 | 233 | 89 | 222 | 33 | 10 | 50 | 307 | 110 |
| Magnocellular nucleus | MA | 22 | 16 | 22 | 23 | 1 | 78 | 6 | 112 | 109 |
| Mammillary body | MBO | 35 | 9 | 12 | 36 | 9 | 0 | 43 | 860 | 53 |
| Medial amygdalar nucleus | MEA | 591 | 409 | 144 | 518 | 167 | 947 | 328 | 873 | 388 |
| Medial geniculate complex | MG | 155 | 126 | 68 | 337 | 85 | 1 | 80 | 584 | 174 |
| Medial habenula | MH | 277 | 410 | 339 | 75 | 117 | 0 | 143 | 31 | 536 |
| Medial preoptic area | MPO | 296 | 359 | 202 | 180 | 156 | 1028 | 276 | 1969 | 981 |
| Medial preoptic nucleus | MPN | 1 | 157 | 49 | 49 | 11 | 349 | 97 | 470 | 181 |
| Medial pretectal area | MPT | 48 | 82 | 29 | 54 | 17 | 0 | 13 | 9 | 69 |
| Medial septal nucleus | MS | 7 | 69 | 25 | 42 | 36 | 50 | 108 | 166 | 166 |
| Median preoptic nucleus | MEPO | 4 | 136 | 26 | 19 | 16 | 66 | 24 | 62 | 193 |
| Mediodorsal nucleus of thalamus | MD | 60 | 189 | 80 | 127 | 161 | 6 | 149 | 173 | 313 |
| Midbrain reticular nucleus | MRN | 324 | 350 | 345 | 864 | 227 | 33 | 510 | 1200 | 561 |
| Midbrain reticular nucleus retrorubral area | RR | 17 | 50 | 77 | 172 | 43 | 4 | 46 | 168 | 34 |
| Nucleus accumbens | ACB | 288 | 954 | 302 | 454 | 214 | 1222 | 445 | 2066 | 1784 |
| Nucleus of Darkschewitsch | ND | 2 | 3 | 4 | 8 | 1 | 0 | 2 | 4 | 0 |
| Nucleus of reunions | RE | 42 | 65 | 42 | 130 | 29 | 32 | 132 | 401 | 68 |
| Nucleus of the brachium of the inferior colliculus | NB | 79 | 64 | 80 | 206 | 78 | 1 | 89 | 105 | 192 |
| Nucleus of the lateral olfactory tract | NLOT | 260 | 30 | 13 | 30 | 3 | 60 | 108 | 107 | 134 |

| | | | | | | | | | | |
|---|---|---|---|---|---|---|---|---|---|---|
| Nucleus of the optic tract | NOT | 21 | 32 | 84 | 161 | 21 | 0 | 167 | 35 | 87 |
| Nucleus of the posterior commissure | NPC | 28 | 55 | 50 | 253 | 10 | 0 | 213 | 27 | 41 |
| Olfactory tubercle | OT | 1125 | 49 | 126 | 115 | 1021 | 565 | 142 | 2816 | 474 |
| Olivary pretectal nucleus | OP | 10 | 169 | 38 | 68 | 15 | 0 | 142 | 15 | 68 |
| Orbital area lateral part | ORBl | 22 | 791 | 59 | 139 | 44 | 852 | 167 | 1553 | 306 |
| Orbital area medial part | ORBm | 65 | 719 | 147 | 102 | 314 | 188 | 66 | 313 | 351 |
| Orbital area ventrolateral part | ORBvl | 25 | 371 | 1480 | 109 | 67 | 1068 | 252 | 1371 | 415 |
| Parabigeminal nucleus | PBG | 12 | 10 | 0 | 17 | 9 | 0 | 27 | 11 | 10 |
| Paracentral nucleus | PCN | 5 | 9 | 2 | 23 | 6 | 12 | 9 | 124 | 18 |
| Parafascicular nucleus | PF | 6 | 18 | 3 | 52 | 7 | 7 | 73 | 67 | 21 |
| Paraflocculus | PFL | 10 | 14 | 10 | 45 | 13 | 0 | 2 | 155 | 30 |
| Parastrial nucleus | PS | 3 | 20 | 4 | 15 | 1 | 21 | 11 | 111 | 29 |
| Parasubiculum | PAR | 10 | 18 | 41 | 69 | 8 | 10 | 8 | 440 | 38 |
| Parasubthalamic nucleus | PSTN | 18 | 41 | 34 | 47 | 17 | 5 | 10 | 70 | 11 |
| Parataenial nucleus | PT | 267 | 267 | 139 | 186 | 182 | 11 | 113 | 16 | 221 |
| Paraventricular hypothalamic nucleus | PVH | 38 | 405 | 85 | 135 | 101 | 21 | 98 | 467 | 399 |
| Paraventricular hypothalamic nucleus descending division | PVHd | 22 | 64 | 12 | 22 | 18 | 19 | 32 | 104 | 38 |
| Paraventricular nucleus of the thalamus | PVT | 600 | 884 | 864 | 95 | 689 | 0 | 31 | 9 | 1019 |
| Pedunculopontine nucleus | PPN | 22 | 23 | 16 | 31 | 11 | 0 | 8 | 47 | 32 |
| Periaqueductal gray | PAG | 172 | 636 | 171 | 503 | 333 | 0 | 195 | 857 | 595 |
| Peripeduncular nucleus | PP | 7 | 12 | 6 | 22 | 1 | 0 | 15 | 15 | 38 |
| Perirhinal area | PERI | 1122 | 691 | 1114 | 1648 | 1295 | 259 | 769 | 2652 | 1793 |
| Periventricular hypothalamic nucleus posterior part | PVp | 12 | 40 | 5 | 49 | 11 | 0 | 6 | 189 | 22 |
| Periventricular hypothalamic nucleus preoptic part | PVpo | 2 | 112 | 26 | 18 | 8 | 38 | 13 | 80 | 203 |
| Periventricular zone | PVZ | 7 | 208 | 41 | 50 | 37 | 19 | 46 | 171 | 226 |
| Piriform area | PIR | 3522 | 4826 | 3418 | 3316 | 5806 | 1158 | 2281 | 5829 | 5116 |
| Piriform-amygdalar area | PAA | 341 | 74 | 32 | 24 | 852 | 12 | 28 | 657 | 173 |
| Pons | P | 369 | 312 | 217 | 358 | 439 | 2 | 219 | 2845 | 1220 |
| Pons motor related | P-mot | 39 | 226 | 92 | 221 | 62 | 0 | 77 | 1433 | 273 |
| Pontine reticular nucleus | PRNr | 51 | 119 | 61 | 124 | 65 | 0 | 76 | 510 | 195 |
| Posterior amygdalar nucleus | PA | 21 | 43 | 9 | 61 | 22 | 36 | 19 | 133 | 64 |
| Posterior auditory area | AUDpo | 88 | 7 | 36 | 150 | 193 | 2 | 47 | 591 | 378 |
| Posterior complex of the thalamus | PO | 20 | 5 | 93 | 30 | 3 | 245 | 66 | 773 | 40 |
| Posterior hypothalamic nucleus | PH | 303 | 441 | 164 | 450 | 77 | 0 | 123 | 1122 | 384 |
| Posterior limiting nucleus of the thalamus | POL | 12 | 7 | 11 | 9 | 7 | 3 | 8 | 14 | 24 |
| Posterior pretectal nucleus | PPT | 3 | 10 | 4 | 38 | 3 | 0 | 89 | 14 | 2 |
| Posterolateral visual area | VISpl | 6 | 37 | 14 | 45 | 1 | 0 | 9 | 117 | 20 |
| posteromedial visual area | VISpm | 9 | 1431 | 442 | 760 | 156 | 22 | 852 | 3071 | 1680 |
| Postpiriform transition area | TR | 223 | 28 | 18 | 69 | 89 | 8 | 22 | 206 | 168 |
| Postsubiculum | POST | 33 | 94 | 342 | 584 | 139 | 59 | 154 | 494 | 308 |
| Precommissural nucleus | PRC | 29 | 72 | 41 | 85 | 26 | 0 | 23 | 6 | 44 |
| Prelimbic area | PL | 279 | 1451 | 368 | 211 | 559 | 250 | 308 | 595 | 1010 |
| Preparasubthalamic nucleus | PST | 6 | 5 | 6 | 12 | 3 | 3 | 3 | 22 | 1 |
| Presubiculum | PRE | 61 | 282 | 371 | 588 | 152 | 23 | 160 | 1080 | 423 |

| | | | | | | | | | | |
|---|---|---|---|---|---|---|---|---|---|---|
| Primary auditory area | AUDp | 1664 | 647 | 819 | 1486 | 2230 | 202 | 771 | 3565 | 2952 |
| Primary motor area | MOp | 716 | 14331 | 5991 | 6321 | 2814 | 11439 | 7902 | 23397 | 11076 |
| Primary somatosensory area barrel field | SSp-bfd | 1286 | 3242 | 3670 | 2285 | 1966 | 7040 | 4277 | 10541 | 4580 |
| Primary somatosensory area lower limb | SSp-ll | 295 | 5333 | 2162 | 2104 | 1110 | 2750 | 1775 | 4944 | 3859 |
| Primary somatosensory area mouth | SSp-m | 1073 | 4813 | 2147 | 711 | 850 | 1896 | 2933 | 4947 | 2214 |
| Primary somatosensory area nose | SSp-n | 1465 | 3286 | 2158 | 948 | 1341 | 2354 | 2290 | 5771 | 2622 |
| Primary somatosensory area trunk | SSp-tr | 56 | 2415 | 1502 | 958 | 477 | 1125 | 1283 | 3297 | 2578 |
| Primary somatosensory area upper limb | SSp-ul | 688 | 5664 | 3453 | 1863 | 927 | 4609 | 3311 | 7481 | 4302 |
| Primary visual area | VISp | 103 | 1793 | 918 | 3118 | 196 | 1672 | 1656 | 7180 | 2810 |
| Red nucleus | RN | 24 | 19 | 30 | 109 | 48 | 0 | 35 | 151 | 60 |
| Reticular nucleus of the thalamus | RT | 90 | 22 | 65 | 401 | 45 | 150 | 126 | 198 | 381 |
| Retrochiasmatic area | RCH | 190 | 81 | 10 | 176 | 52 | 250 | 89 | 1099 | 550 |
| Retrosplenial area dorsal part | RSPd | 107 | 3384 | 1282 | 1977 | 505 | 0 | 2341 | 5927 | 2977 |
| Retrosplenial area lateral agranular part | RSPagl | 47 | 1668 | 829 | 2169 | 547 | 0 | 954 | 5342 | 1827 |
| Retrosplenial area ventral part | RSPv | 228 | 2604 | 901 | 4150 | 1273 | 6 | 2906 | 5708 | 3226 |
| Secondary motor area | MOs | 645 | 7208 | 2528 | 2831 | 1740 | 4008 | 5508 | 10852 | 6794 |
| Septofimbrial nucleus | SF | 13 | 186 | 54 | 75 | 97 | 19 | 53 | 78 | 139 |
| Simple lobule | SIM | 20 | 40 | 22 | 728 | 15 | 1 | 156 | 218 | 239 |
| Subiculum | SUB | 252 | 270 | 279 | 1280 | 244 | 277 | 849 | 1986 | 811 |
| Submedial nucleus of the thalamus | SMT | 7 | 7 | 5 | 28 | 2 | 11 | 10 | 41 | 33 |
| Subparafascicular nucleus | SPF | 73 | 81 | 62 | 180 | 59 | 6 | 60 | 269 | 197 |
| Subparaventricular zone | SBPV | 5 | 248 | 56 | 90 | 7 | 40 | 76 | 222 | 156 |
| Substantia innominata | SI | 177 | 170 | 163 | 208 | 176 | 941 | 148 | 1496 | 397 |
| Substantia nigra compact part | SNc | 21 | 34 | 11 | 66 | 12 | 8 | 31 | 246 | 20 |
| Substantia nigra reticular part | SNr | 86 | 59 | 26 | 117 | 22 | 45 | 106 | 1022 | 97 |
| Subthalamic nucleus | STN | 30 | 36 | 14 | 36 | 9 | 18 | 20 | 152 | 42 |
| Superior colliculus motor related | SCm | 253 | 1169 | 890 | 2795 | 414 | 44 | 995 | 2742 | 1486 |
| Superior colliculus sensory related | SCs | 48 | 1520 | 602 | 2794 | 244 | 3 | 387 | 1633 | 706 |
| Supplemental somatosensory area | SSs | 3479 | 5752 | 5353 | 2603 | 3489 | 1380 | 2529 | 6370 | 3864 |
| Suprachiasmatic nucleus | SCH | 0 | 184 | 14 | 23 | 81 | 80 | 79 | 157 | 493 |
| Supramammillary nucleus | SUM | 8 | 32 | 22 | 16 | 6 | 0 | 12 | 218 | 28 |
| Supraoptic nucleus | SO | 2 | 33 | 30 | 4 | 40 | 44 | 4 | 97 | 248 |
| Taenia tecta | TT | 151 | 755 | 474 | 177 | 194 | 478 | 513 | 1159 | 601 |
| Temporal association areas | TEa | 1914 | 788 | 2007 | 2818 | 3272 | 336 | 1457 | 6394 | 4439 |
| Thalamus sensory-motor cortex related | DORsm | 4 | 5 | 6 | 45 | 15 | 0 | 5 | 9 | 40 |
| Triangular nucleus of septum | TRS | 14 | 207 | 5 | 143 | 48 | 0 | 28 | 2 | 93 |
| Tuberal nucleus | TU | 78 | 167 | 50 | 257 | 94 | 171 | 94 | 1184 | 372 |
| Ventral anterior-lateral complex of the thalamus | VAL | 25 | 21 | 42 | 30 | 19 | 95 | 44 | 185 | 39 |
| Ventral auditory area | AUDv | 1953 | 1330 | 1815 | 2191 | 2663 | 381 | 1079 | 4339 | 3884 |
| Ventral medial nucleus of the thalamus | VM | 79 | 127 | 55 | 152 | 86 | 320 | 92 | 1094 | 166 |
| Ventral part of the lateral geniculate complex | LGv | 219 | 194 | 132 | 239 | 174 | 3 | 137 | 210 | 289 |
| Ventral posterior complex of the thalamus | VP | 38 | 24 | 39 | 102 | 36 | 435 | 44 | 634 | 81 |
| Ventral posterolateral nucleus of the thalamus | VPL | 104 | 31 | 30 | 126 | 64 | 17 | 21 | 44 | 48 |
| Ventral tegmental area | VTA | 199 | 135 | 94 | 495 | 83 | 1 | 438 | 1789 | 168 |

| | | | | | | | | | | |
|---|---|---|---|---|---|---|---|---|---|---|
| Ventrolateral preoptic nucleus | VLPO | 67 | 8 | 4 | 1 | 76 | 22 | 18 | 84 | 93 |
| Ventromedial hypothalamic nucleus | VMH | 40 | 71 | 13 | 75 | 30 | 194 | 93 | 1267 | 159 |
| Vestibular nuclei | VNC | 20 | 90 | 57 | 137 | 7 | 0 | 152 | 428 | 32 |
| Visceral area | VISC | 866 | 1114 | 1762 | 974 | 1073 | 248 | 655 | 1578 | 1124 |
| Zona incerta | ZI | 348 | 277 | 151 | 464 | 227 | 520 | 360 | 1296 | 559 |

Table 4-1: List of significantly correlated genes (Fig. 4C with false discovery rate. (FDR) at 5%)

| | set | n | cor | p | FDR | Bonferroni |
|---|---|---|---|---|---|---|
| 175 | Arl6 | 156 | 0.49504278 | 5.05E-11 | 7.41E-07 | 1.29E-06 |
| 1001 | Mpo | 156 | 0.49366662 | 5.82E-11 | 7.41E-07 | 1.48E-06 |
| 1066 | Nek7 | 160 | -0.4641837 | 6.31E-10 | 4.22E-06 | 1.61E-05 |
| 990 | Mmp15 | 158 | 0.46347895 | 8.65E-10 | 4.22E-06 | 2.20E-05 |
| 299 | Cd8b1 | 154 | 0.46750657 | 9.79E-10 | 4.22E-06 | 2.49E-05 |
| 1519 | Sycn | 156 | 0.46365193 | 1.09E-09 | 4.22E-06 | 2.78E-05 |
| 773 | Igf2bp1 | 156 | 0.46233287 | 1.23E-09 | 4.22E-06 | 3.14E-05 |
| 564 | Fam53a | 156 | 0.46153767 | 1.33E-09 | 4.22E-06 | 3.38E-05 |
| 1036 | Mypn | 155 | 0.45112842 | 3.84E-09 | 1.09E-05 | 9.78E-05 |
| 1664 | Vars | 155 | 0.4499021 | 4.28E-09 | 1.09E-05 | 0.00010898 |
| 382 | Col2a1 | 159 | 0.44360592 | 4.73E-09 | 1.10E-05 | 0.00012049 |
| 1135 | Orm1 | 152 | 0.44975281 | 6.14E-09 | 1.30E-05 | 0.00015645 |
| 1508 | Styxl1 | 156 | 0.44113478 | 8.22E-09 | 1.52E-05 | 0.00020935 |
| 580 | Fbxl4 | 151 | 0.44748325 | 8.38E-09 | 1.52E-05 | 0.0002135 |
| 1275 | Rab13 | 155 | 0.44131491 | 9.05E-09 | 1.54E-05 | 0.0002304 |
| 876 | Lhx8 | 159 | 0.43277691 | 1.22E-08 | 1.94E-05 | 0.00031011 |
| 541 | Etfa | 160 | -0.4281883 | 1.62E-08 | 2.43E-05 | 0.00041296 |
| 1516 | Suv420h2 | 155 | 0.43266103 | 1.88E-08 | 2.65E-05 | 0.00047993 |
| 153 | Appbp2 | 156 | 0.43080749 | 1.98E-08 | 2.65E-05 | 0.00050392 |
| 369 | Cmah | 158 | 0.42577597 | 2.44E-08 | 3.09E-05 | 0.00062249 |
| 1025 | Mucl1 | 156 | 0.42778783 | 2.54E-08 | 3.09E-05 | 0.00064791 |
| 1426 | Slc2a1 | 158 | -0.4215244 | 3.48E-08 | 4.02E-05 | 0.000885 |
| 283 | Ccdc88b | 156 | 0.42201815 | 4.09E-08 | 4.52E-05 | 0.00104025 |
| 1218 | Pmaip1 | 158 | 0.41892355 | 4.30E-08 | 4.56E-05 | 0.00109494 |
| 1675 | Vps37c | 151 | 0.42695566 | 4.58E-08 | 4.66E-05 | 0.00116599 |
| 797 | Isl1 | 160 | 0.41514068 | 4.81E-08 | 4.71E-05 | 0.00122377 |
| 1039 | N4bp1 | 158 | 0.41698715 | 5.03E-08 | 4.75E-05 | 0.00128148 |
| 1072 | Ngfr | 158 | 0.41632383 | 5.31E-08 | 4.83E-05 | 0.00135213 |
| 871 | Lgals8 | 156 | 0.4176704 | 5.80E-08 | 5.10E-05 | 0.0014776 |
| 1229 | Ppih | 154 | 0.41898946 | 6.37E-08 | 5.13E-05 | 0.00162203 |
| 1198 | Pik3r3 | 160 | -0.4116379 | 6.38E-08 | 5.13E-05 | 0.00162549 |
| 1413 | Slc17a9 | 158 | 0.41392053 | 6.44E-08 | 5.13E-05 | 0.00164072 |
| 653 | Gm13420 | 160 | 0.4053327 | 1.06E-07 | 8.14E-05 | 0.0026877 |
| 58 | 9530077C05Rik | 160 | 0.40358046 | 1.21E-07 | 8.85E-05 | 0.00308517 |
| 1700 | Xbp1 | 160 | -0.4035334 | 1.22E-07 | 8.85E-05 | 0.00309659 |
| 1407 | Slc15a2 | 154 | 0.40998692 | 1.29E-07 | 9.12E-05 | 0.0032832 |
| 475 | Dph6 | 156 | 0.40541197 | 1.52E-07 | 0.00010463 | 0.00387137 |
| 863 | Lect1 | 156 | -0.4044905 | 1.63E-07 | 0.00010936 | 0.00415564 |
| 1626 | Ttll12 | 152 | 0.40508863 | 2.26E-07 | 0.00014582 | 0.00575088 |
| 448 | Depdc7 | 155 | 0.40119397 | 2.30E-07 | 0.00014582 | 0.00585341 |
| 1242 | Pramel7 | 155 | 0.40091328 | 2.35E-07 | 0.00014582 | 0.00597869 |

| | | | | | | |
|---|---|---|---|---|---|---|
| 1455 | Smpd2 | 156 | 0.39917528 | 2.45E-07 | 0.00014828 | 0.00622794 |
| 1080 | Nlrx1 | 156 | 0.39792642 | 2.69E-07 | 0.00015762 | 0.00684204 |
| 932 | Luc7l | 152 | 0.40215621 | 2.81E-07 | 0.00015762 | 0.00715779 |
| 873 | Lhx6 | 156 | 0.39719322 | 2.84E-07 | 0.00015762 | 0.00722912 |
| 1240 | Ppp4c | 156 | 0.39715417 | 2.85E-07 | 0.00015762 | 0.00725031 |
| 1363 | Sass6 | 155 | 0.39790404 | 2.94E-07 | 0.00015943 | 0.00749328 |
| 704 | Gstk1 | 160 | -0.3904065 | 3.33E-07 | 0.00017165 | 0.00848676 |
| 213 | Baat | 152 | 0.39983062 | 3.34E-07 | 0.00017165 | 0.00850178 |
| 1537 | Tap1 | 155 | 0.39608111 | 3.37E-07 | 0.00017165 | 0.0085824 |
| 527 | Epas1 | 158 | -0.3918164 | 3.56E-07 | 0.00017784 | 0.00906969 |
| 469 | Dnajb4 | 152 | 0.39827977 | 3.74E-07 | 0.00018324 | 0.00952862 |
| 1601 | Tnfrsf11b | 156 | 0.39288985 | 3.91E-07 | 0.00018657 | 0.00995878 |
| 953 | Mapkap1 | 156 | 0.39261405 | 3.99E-07 | 0.00018657 | 0.01016377 |
| 726 | Hbb-b2 | 160 | -0.3878695 | 4.03E-07 | 0.00018657 | 0.01026138 |
| 101 | Adm | 153 | 0.39481773 | 4.41E-07 | 0.0002006 | 0.01123368 |
| 521 | Enox2 | 154 | 0.39218022 | 4.90E-07 | 0.00021891 | 0.01247804 |
| 786 | Insl5 | 156 | 0.38927721 | 5.10E-07 | 0.0002239 | 0.01298625 |
| 1405 | Slc14a1 | 157 | 0.38775626 | 5.24E-07 | 0.00022597 | 0.0133325 |
| 1053 | Ncdn | 160 | -0.3814801 | 6.46E-07 | 0.00027396 | 0.0164378 |
| 858 | Lap3 | 156 | 0.38364031 | 7.67E-07 | 0.0003192 | 0.01952683 |
| 1216 | Pltp | 160 | -0.3789255 | 7.77E-07 | 0.0003192 | 0.01979039 |
| 1589 | Tmem183a | 156 | 0.38279934 | 8.14E-07 | 0.00032918 | 0.02073855 |
| 1131 | Oog3 | 155 | 0.38358576 | 8.36E-07 | 0.00033268 | 0.02129177 |
| 182 | Art4 | 156 | 0.3820081 | 8.62E-07 | 0.0003376 | 0.02194382 |
| 969 | Med27 | 125 | 0.42265788 | 9.10E-07 | 0.00035122 | 0.0231808 |
| 473 | Dnase2a | 155 | 0.38207512 | 9.31E-07 | 0.00035379 | 0.02370411 |
| 780 | Il17rb | 155 | 0.38145582 | 9.73E-07 | 0.00035493 | 0.02476652 |
| 1073 | Nhp2l1 | 152 | 0.38496076 | 9.73E-07 | 0.00035493 | 0.02477913 |
| 1111 | Oas1c | 155 | 0.38141091 | 9.76E-07 | 0.00035493 | 0.0248453 |
| 791 | Irs4 | 160 | 0.37506123 | 1.03E-06 | 0.00036289 | 0.02612755 |
| 14 | 2310022A10Rik | 155 | 0.38033535 | 1.05E-06 | 0.00036289 | 0.02680506 |
| 897 | LOC433005 | 159 | -0.3756483 | 1.06E-06 | 0.00036289 | 0.02711357 |
| 117 | Akr1b8 | 151 | 0.38481094 | 1.07E-06 | 0.00036289 | 0.02721494 |
| 1442 | Slc6a20b | 160 | -0.3744887 | 1.07E-06 | 0.00036289 | 0.02721698 |
| 711 | H2-D1 | 159 | -0.375018 | 1.11E-06 | 0.0003731 | 0.02835558 |
| 271 | Cbx5 | 154 | 0.37986645 | 1.18E-06 | 0.00038605 | 0.03004144 |
| 1237 | Ppp1r3d | 156 | -0.3775222 | 1.18E-06 | 0.00038605 | 0.03014353 |
| 1334 | Rnpc3 | 150 | 0.3842794 | 1.21E-06 | 0.00038605 | 0.03068622 |
| 1340 | Rps13 | 154 | 0.37947035 | 1.21E-06 | 0.00038605 | 0.03088437 |
| 1403 | Slamf7 | 152 | 0.38136536 | 1.25E-06 | 0.00039315 | 0.03184478 |
| 144 | Ap4m1 | 156 | 0.37589037 | 1.33E-06 | 0.00041137 | 0.03379438 |
| 511 | Eif6 | 156 | 0.37574311 | 1.34E-06 | 0.00041137 | 0.03414381 |
| 364 | Clk1 | 156 | -0.3753641 | 1.38E-06 | 0.00041737 | 0.0350589 |
| 1553 | Tcl1b1 | 156 | 0.37428457 | 1.48E-06 | 0.00043999 | 0.03779545 |
| 1307 | Rgs5 | 158 | -0.3720293 | 1.49E-06 | 0.00043999 | 0.0378395 |

| | | | | | | |
|---|---|---|---|---|---|---|
| 1287 | Rbbp6 | 155 | 0.37494314 | 1.53E-06 | 0.00044547 | 0.03906215 |
| 746 | Hook3 | 156 | 0.37375864 | 1.54E-06 | 0.00044547 | 0.03920115 |
| 108 | Ahsg | 155 | 0.37465626 | 1.56E-06 | 0.0004477 | 0.03984509 |
| 841 | Klhl15 | 156 | 0.37333569 | 1.59E-06 | 0.0004477 | 0.04036755 |
| 427 | Cyp26a1 | 156 | 0.37320293 | 1.60E-06 | 0.0004477 | 0.04074044 |
| 45 | 5031414D18Rik | 158 | 0.37072944 | 1.63E-06 | 0.0004503 | 0.04142729 |
| 1691 | Wisp1 | 159 | 0.36743558 | 1.89E-06 | 0.00051873 | 0.0482423 |
| 461 | Dlx3 | 160 | -0.3658268 | 1.96E-06 | 0.00053216 | 0.05002259 |
| 120 | Aldh1l1 | 160 | -0.365581 | 2.00E-06 | 0.00053559 | 0.05088107 |
| 614 | Gab1 | 159 | -0.3653867 | 2.18E-06 | 0.00057628 | 0.05556359 |
| 1402 | Six3os1 | 160 | 0.36421849 | 2.20E-06 | 0.00057628 | 0.05589916 |
| 949 | Map1lc3b | 160 | -0.363035 | 2.38E-06 | 0.00061632 | 0.0606366 |
| 1138 | Ostc | 158 | 0.36510853 | 2.40E-06 | 0.00061632 | 0.06101611 |
| 380 | Cntrl | 159 | -0.3633043 | 2.52E-06 | 0.00064081 | 0.06408054 |
| 1090 | Nrg4 | 158 | 0.36412586 | 2.56E-06 | 0.00064595 | 0.06524058 |
| 994 | Mmp9 | 156 | 0.36605425 | 2.61E-06 | 0.00065141 | 0.06644378 |
| 484 | Duoxa1 | 156 | 0.36588997 | 2.64E-06 | 0.00065229 | 0.06718564 |
| 1714 | Zbbx | 158 | -0.3632251 | 2.72E-06 | 0.00065428 | 0.06935612 |
| 1078 | Nit2 | 158 | 0.36321758 | 2.73E-06 | 0.00065428 | 0.06939133 |
| 1141 | Otx2 | 154 | -0.3673893 | 2.77E-06 | 0.00065428 | 0.07059471 |
| 501 | Efhd1 | 155 | 0.36623403 | 2.78E-06 | 0.00065428 | 0.0707554 |
| 358 | Cldn5 | 160 | -0.3607449 | 2.78E-06 | 0.00065428 | 0.07090911 |
| 1178 | Peg10 | 150 | 0.37167401 | 2.83E-06 | 0.00065428 | 0.07204334 |
| 1179 | Peg10 | 160 | 0.36050174 | 2.83E-06 | 0.00065428 | 0.07209217 |
| 988 | Mmp11 | 151 | 0.37039008 | 2.85E-06 | 0.00065428 | 0.07262464 |
| 1397 | Shroom1 | 158 | 0.3621379 | 2.93E-06 | 0.00066653 | 0.07465192 |
| 958 | Matn4 | 155 | 0.36440426 | 3.14E-06 | 0.00070784 | 0.07998631 |
| 396 | Cpa1 | 156 | 0.3627855 | 3.25E-06 | 0.00072321 | 0.08277586 |
| 388 | Colec11 | 160 | 0.35810725 | 3.33E-06 | 0.00072321 | 0.08478805 |
| 794 | Irx2 | 160 | -0.3580114 | 3.35E-06 | 0.00072321 | 0.08533807 |
| 1647 | Uck2 | 158 | 0.36015014 | 3.35E-06 | 0.00072321 | 0.08534285 |
| 1021 | Mttp | 156 | 0.36220816 | 3.38E-06 | 0.00072321 | 0.08603074 |
| 761 | Htr3b | 156 | 0.36220269 | 3.38E-06 | 0.00072321 | 0.08606216 |
| 634 | Gatsl3 | 160 | 0.35763661 | 3.44E-06 | 0.00072934 | 0.08752138 |
| 225 | Bcs1l | 155 | 0.3626769 | 3.52E-06 | 0.00073944 | 0.08974111 |
| 30 | 3830406C13Rik | 160 | -0.3571868 | 3.54E-06 | 0.00073944 | 0.09021192 |
| 1378 | Serpina1d | 158 | 0.35900162 | 3.62E-06 | 0.00074639 | 0.09216682 |
| 659 | Gm4912 | 159 | -0.3578494 | 3.64E-06 | 0.00074639 | 0.09266571 |
| 865 | Lefty1 | 155 | 0.36209099 | 3.66E-06 | 0.00074639 | 0.093299 |
| 494 | Ebf4 | 160 | 0.3564076 | 3.73E-06 | 0.00075338 | 0.09505909 |
| 751 | Hsd17b13 | 159 | 0.35737131 | 3.76E-06 | 0.00075338 | 0.0956796 |
| 325 | Celf6 | 155 | 0.36153085 | 3.80E-06 | 0.00075645 | 0.09682525 |
| 408 | Crisp1 | 155 | 0.36056209 | 4.05E-06 | 0.00080019 | 0.1032244 |
| 414 | Ctage5 | 156 | 0.35773751 | 4.54E-06 | 0.00088187 | 0.11568011 |
| 735 | Hiatl1 | 158 | -0.3555484 | 4.55E-06 | 0.00088187 | 0.1159421 |

| | | | | | | |
|---|---|---|---|---|---|---|
| 1532 | Tacr2 | 153 | -0.3609332 | 4.58E-06 | 0.00088187 | 0.11652135 |
| 146 | Apbb1 | 160 | -0.3532583 | 4.61E-06 | 0.00088187 | 0.11728832 |
| 723 | Hbb | 160 | -0.3530075 | 4.68E-06 | 0.00088956 | 0.11925596 |
| 1003 | Mpzl2 | 155 | 0.35825853 | 4.72E-06 | 0.00088956 | 0.12009022 |
| 609 | Fzd7 | 157 | -0.3559162 | 4.77E-06 | 0.00089125 | 0.12144656 |
| 1583 | Tmem125 | 158 | 0.35476403 | 4.80E-06 | 0.00089125 | 0.12210087 |
| 1341 | Rps15 | 155 | 0.3576056 | 4.92E-06 | 0.00090817 | 0.12532683 |
| 478 | Drd3 | 155 | 0.35727622 | 5.03E-06 | 0.00091939 | 0.12804997 |
| 852 | Kti12 | 158 | 0.35396245 | 5.05E-06 | 0.00091939 | 0.12871425 |
| 618 | Gabrq | 160 | 0.35154967 | 5.16E-06 | 0.00092568 | 0.13132846 |
| 160 | Aqp4 | 160 | -0.3515361 | 5.16E-06 | 0.00092568 | 0.13144605 |
| 1384 | Setx | 159 | -0.3523856 | 5.23E-06 | 0.00093132 | 0.13317901 |
| 1679 | Vtn | 160 | -0.3510838 | 5.32E-06 | 0.00094045 | 0.13542447 |
| 1172 | Pdha1 | 160 | -0.3509534 | 5.36E-06 | 0.00094202 | 0.13659284 |
| 532 | Epha10 | 160 | -0.3507756 | 5.43E-06 | 0.00094593 | 0.13820122 |
| 316 | Cdk20 | 152 | 0.35932499 | 5.46E-06 | 0.00094593 | 0.13905154 |
| 810 | Jmjd8 | 155 | 0.35584601 | 5.52E-06 | 0.00094958 | 0.14053847 |
| 276 | Ccdc155 | 152 | 0.35899978 | 5.58E-06 | 0.00095298 | 0.14199457 |
| 700 | Grwd1 | 152 | 0.35873069 | 5.67E-06 | 0.00096316 | 0.14447428 |
| 159 | Aqp4 | 160 | -0.3499692 | 5.72E-06 | 0.00096505 | 0.14572278 |
| 70 | Abcc4 | 159 | -0.3500914 | 6.08E-06 | 0.00101277 | 0.15477739 |
| 269 | Cbwd1 | 158 | 0.35105851 | 6.11E-06 | 0.00101277 | 0.15563099 |
| 627 | Galnt10 | 149 | 0.36093301 | 6.12E-06 | 0.00101277 | 0.15596668 |
| 1631 | Tubb2b | 146 | -0.3638483 | 6.35E-06 | 0.00103883 | 0.16174735 |
| 1484 | Spc25 | 155 | 0.35364286 | 6.36E-06 | 0.00103883 | 0.16205816 |
| 1689 | Wee2 | 152 | 0.35603084 | 6.74E-06 | 0.00109383 | 0.17173165 |
| 1367 | Scara5 | 155 | 0.35250753 | 6.85E-06 | 0.00110337 | 0.17433182 |
| 1254 | Prss58 | 158 | -0.3490415 | 6.97E-06 | 0.00111558 | 0.17737771 |
| 118 | Akr1c13 | 153 | 0.35421376 | 7.06E-06 | 0.00112311 | 0.17969724 |
| 970 | Med7 | 155 | 0.35160905 | 7.25E-06 | 0.00114698 | 0.18466369 |
| 1505 | Stoml2 | 160 | -0.3459877 | 7.42E-06 | 0.00116607 | 0.18890274 |
| 116 | Aknaos | 156 | 0.34981853 | 7.59E-06 | 0.00118617 | 0.19334652 |
| 1467 | Snx16 | 144 | -0.3629588 | 7.79E-06 | 0.00120853 | 0.19826617 |
| 1599 | Tmem9 | 158 | 0.34722429 | 7.83E-06 | 0.00120853 | 0.19940777 |
| 1678 | Vtn | 160 | -0.3449702 | 7.92E-06 | 0.00121532 | 0.20174268 |
| 1116 | Odf3b | 159 | -0.345688 | 8.08E-06 | 0.00122769 | 0.20585793 |
| 552 | Fam122c | 156 | 0.34871698 | 8.15E-06 | 0.00122769 | 0.2074406 |
| 1269 | Pwp1 | 155 | 0.34978269 | 8.15E-06 | 0.00122769 | 0.2074792 |
| 548 | F2 | 158 | 0.346335 | 8.29E-06 | 0.00124182 | 0.21110973 |
| 138 | Anxa13 | 156 | 0.34819259 | 8.42E-06 | 0.00125431 | 0.21448662 |
| 1358 | S100b | 158 | -0.3455673 | 8.71E-06 | 0.00127887 | 0.22173184 |
| 829 | Kctd9 | 152 | 0.35195746 | 8.73E-06 | 0.00127887 | 0.22223196 |
| 1721 | Zcchc5 | 158 | 0.34551142 | 8.74E-06 | 0.00127887 | 0.22252421 |
| 190 | Atf3 | 155 | 0.3483847 | 8.90E-06 | 0.00129554 | 0.2267194 |
| 747 | Hoxa10 | 160 | 0.34299594 | 8.99E-06 | 0.00130141 | 0.22904771 |

| | | | | | | |
|---|---|---|---|---|---|---|
| 790 | Iqub | 160 | -0.3428216 | 9.10E-06 | 0.00130859 | 0.23162004 |
| 918 | LOC545854 | 160 | -0.342656 | 9.19E-06 | 0.00131511 | 0.23408897 |
| 720 | Hba-a1 | 155 | -0.3477245 | 9.28E-06 | 0.00132057 | 0.23638126 |
| 1257 | Psg16 | 159 | -0.3432965 | 9.42E-06 | 0.00133183 | 0.23991398 |
| 694 | Grin3b | 159 | 0.34321895 | 9.47E-06 | 0.00133183 | 0.24110292 |
| 1336 | Rpgrip1 | 160 | -0.3420913 | 9.53E-06 | 0.00133183 | 0.24269639 |
| 1228 | Ppargc1b | 156 | -0.3461776 | 9.57E-06 | 0.00133183 | 0.24372501 |
| 1289 | Rbm17 | 156 | 0.34609022 | 9.62E-06 | 0.00133193 | 0.245075 |
| 976 | Mettl22 | 155 | 0.34673305 | 9.88E-06 | 0.00135654 | 0.25162587 |
| 728 | Hck | 156 | 0.345629 | 9.91E-06 | 0.00135654 | 0.25231632 |
| 538 | Esam | 155 | -0.3457181 | 1.05E-05 | 0.00143418 | 0.26819135 |
| 849 | Kremen1 | 154 | -0.3463063 | 1.09E-05 | 0.00146326 | 0.27635013 |
| 833 | Kif15 | 156 | 0.34417133 | 1.09E-05 | 0.00146326 | 0.27655583 |
| 1404 | Slc10a4 | 159 | 0.34053459 | 1.12E-05 | 0.00150457 | 0.2858686 |
| 848 | Krcc1 | 156 | 0.3434767 | 1.13E-05 | 0.00151239 | 0.28886614 |
| 219 | Bcan | 160 | -0.3388731 | 1.17E-05 | 0.00154594 | 0.2977498 |
| 1214 | Plscr1 | 156 | 0.3429595 | 1.17E-05 | 0.00154594 | 0.2983666 |
| 1434 | Slc3a1 | 158 | 0.34075903 | 1.18E-05 | 0.00154599 | 0.30066719 |
| 286 | Ccl22 | 155 | 0.34384702 | 1.18E-05 | 0.00154599 | 0.30146807 |
| 1321 | Riok2 | 154 | 0.34473305 | 1.20E-05 | 0.00155508 | 0.30479586 |
| 133 | Amph | 160 | -0.33819 | 1.22E-05 | 0.001578 | 0.31086504 |
| 287 | Ccnb1 | 155 | 0.34268844 | 1.27E-05 | 0.00163631 | 0.32398979 |
| 824 | Kcnmb4os2 | 159 | -0.3382798 | 1.29E-05 | 0.00165028 | 0.32943433 |
| 502 | Ehmt2 | 159 | -0.3382497 | 1.30E-05 | 0.00165028 | 0.33005552 |
| 603 | Frg1 | 158 | 0.33860138 | 1.35E-05 | 0.00171211 | 0.34413415 |
| 303 | Cdc20 | 156 | 0.34045393 | 1.37E-05 | 0.00172635 | 0.34872251 |
| 721 | Hba-a1 | 160 | -0.3360844 | 1.39E-05 | 0.00174783 | 0.35480931 |
| 464 | Dmrtc1c2 | 160 | 0.3352056 | 1.47E-05 | 0.00183202 | 0.37483475 |
| 1499 | Stambpl1 | 155 | 0.34029858 | 1.47E-05 | 0.00183202 | 0.37556337 |
| 1088 | Nr2c1 | 159 | 0.33570116 | 1.52E-05 | 0.0018783 | 0.38692887 |
| 65 | Aaas | 156 | 0.338597 | 1.54E-05 | 0.00188943 | 0.39111127 |
| 1038 | Mzt2 | 155 | 0.33938583 | 1.56E-05 | 0.00190978 | 0.39723327 |
| 1423 | Slc25a5 | 160 | -0.3339135 | 1.60E-05 | 0.00194366 | 0.40622472 |
| 769 | Ifi202b | 153 | 0.34091181 | 1.62E-05 | 0.00196033 | 0.41166939 |
| 703 | Gsta3 | 155 | 0.33854997 | 1.64E-05 | 0.00197913 | 0.41810833 |
| 1634 | Txndc5 | 156 | 0.33745393 | 1.65E-05 | 0.00197913 | 0.41957564 |
| 1746 | Zfx | 158 | -0.3352756 | 1.66E-05 | 0.00198339 | 0.42292892 |
| 1069 | Ngb | 160 | 0.33320603 | 1.67E-05 | 0.00198339 | 0.42444591 |
| 449 | Dera | 155 | 0.33783001 | 1.72E-05 | 0.00201347 | 0.43691436 |
| 103 | Agap3 | 160 | -0.3327282 | 1.72E-05 | 0.00201347 | 0.43718748 |
| 1108 | Nxph1 | 160 | -0.3326799 | 1.72E-05 | 0.00201347 | 0.43849538 |
| 1464 | Snhg7 | 160 | -0.332652 | 1.72E-05 | 0.00201347 | 0.43925179 |
| 266 | Cartpt | 158 | 0.33454854 | 1.74E-05 | 0.00201347 | 0.44228662 |
| 61 | A830019P07Rik | 156 | -0.3365683 | 1.74E-05 | 0.00201347 | 0.44296243 |
| 1005 | Mrfap1 | 155 | -0.3372501 | 1.78E-05 | 0.00204763 | 0.45264045 |

| | | | | | | |
|---|---|---|---|---|---|---|
| 1130 | Onecut1 | 158 | -0.3341025 | 1.79E-05 | 0.00204763 | 0.45457348 |
| 1019 | Mtif2 | 151 | 0.34129513 | 1.80E-05 | 0.00205369 | 0.45797231 |
| 367 | Cltc | 159 | -0.3328698 | 1.81E-05 | 0.00205541 | 0.4609203 |
| 161 | Ar | 160 | 0.33181775 | 1.82E-05 | 0.00205541 | 0.46246808 |
| 218 | BC051665 | 155 | 0.33671668 | 1.84E-05 | 0.00206072 | 0.46757425 |
| 466 | Dnah12 | 154 | -0.3376281 | 1.85E-05 | 0.00206072 | 0.47133669 |
| 1318 | Rin2 | 156 | 0.33546693 | 1.86E-05 | 0.00206072 | 0.47376119 |
| 666 | Gm953 | 160 | -0.3313738 | 1.87E-05 | 0.00206072 | 0.47529102 |
| 914 | LOC545133 | 159 | -0.3323517 | 1.87E-05 | 0.00206072 | 0.47582994 |
| 946 | Magt1 | 155 | 0.33637855 | 1.87E-05 | 0.00206072 | 0.47728078 |
| 363 | Clic4 | 160 | -0.3312785 | 1.88E-05 | 0.00206072 | 0.47808678 |
| 1388 | Sfxn2 | 155 | 0.33600079 | 1.92E-05 | 0.00209592 | 0.4883492 |
| 265 | Cartpt | 160 | 0.3303326 | 1.99E-05 | 0.00216117 | 0.50668998 |
| 435 | Dbr1 | 160 | -0.3302945 | 1.99E-05 | 0.00216117 | 0.50787494 |
| 1032 | Myg1 | 155 | 0.33510936 | 2.02E-05 | 0.00218405 | 0.51543569 |
| 779 | Il17b | 152 | 0.33806444 | 2.05E-05 | 0.00219276 | 0.52140727 |
| 130 | Alppl2 | 157 | 0.33281534 | 2.05E-05 | 0.00219276 | 0.52313285 |
| 1474 | Son | 160 | -0.3297822 | 2.06E-05 | 0.00219276 | 0.52407032 |
| 250 | C6 | 158 | 0.33155897 | 2.09E-05 | 0.0022126 | 0.53102448 |
| 892 | LOC432692 | 159 | -0.3302197 | 2.13E-05 | 0.00224938 | 0.54210142 |
| 354 | Cisd2 | 152 | 0.33671641 | 2.22E-05 | 0.00233544 | 0.56517618 |
| 1597 | Tmem38b | 157 | 0.33097364 | 2.30E-05 | 0.0024038 | 0.5848386 |
| 66 | Aadac | 156 | 0.33194046 | 2.30E-05 | 0.0024038 | 0.58652789 |
| 1010 | Mrps23 | 156 | 0.33144013 | 2.37E-05 | 0.00246455 | 0.6044403 |
| 1187 | Pglyrp2 | 155 | 0.33241208 | 2.38E-05 | 0.00246455 | 0.60627835 |
| 24 | 2610305D13Rik | 159 | -0.3281774 | 2.41E-05 | 0.0024845 | 0.61367146 |
| 1432 | Slc38a2 | 156 | -0.3308047 | 2.47E-05 | 0.00253199 | 0.62793286 |
| 1711 | Ywhae | 160 | -0.3264423 | 2.52E-05 | 0.00257906 | 0.64218614 |
| 1742 | Zfp819 | 152 | 0.33447791 | 2.54E-05 | 0.00258236 | 0.64558891 |
| 1577 | Tmbim6 | 160 | -0.3259787 | 2.59E-05 | 0.00263124 | 0.66044012 |
| 148 | Aplf | 158 | 0.32775568 | 2.62E-05 | 0.00265187 | 0.66827144 |
| 945 | Magel2 | 160 | 0.32536421 | 2.69E-05 | 0.00270904 | 0.68538755 |
| 72 | Abhd4 | 152 | 0.33324482 | 2.73E-05 | 0.00273373 | 0.69436853 |
| 319 | Cdr2 | 160 | -0.3247281 | 2.80E-05 | 0.00278463 | 0.71214628 |
| 948 | Map1b | 158 | -0.326665 | 2.80E-05 | 0.00278463 | 0.71341101 |
| 1201 | Piwil4 | 156 | -0.3286149 | 2.81E-05 | 0.00278463 | 0.71564902 |
| 75 | Acacb | 160 | -0.3245211 | 2.83E-05 | 0.00279483 | 0.72106531 |
| 1696 | Wrn | 158 | -0.3259243 | 2.93E-05 | 0.00287866 | 0.74568922 |
| 1610 | Tpi1 | 157 | -0.3268568 | 2.94E-05 | 0.00287866 | 0.74845267 |
| 315 | Cdhr4 | 158 | -0.3251614 | 3.06E-05 | 0.0029899 | 0.78036405 |
| 1512 | Sult1c1 | 156 | 0.32694305 | 3.10E-05 | 0.00300957 | 0.7902504 |
| 203 | Atp6v0a4 | 156 | 0.32685809 | 3.12E-05 | 0.00300957 | 0.79423021 |
| 1390 | Sgk1 | 158 | -0.324859 | 3.12E-05 | 0.00300957 | 0.79452537 |
| 1278 | Rad23b | 155 | -0.3275842 | 3.17E-05 | 0.0030477 | 0.80763991 |
| 916 | LOC545666 | 159 | -0.323468 | 3.20E-05 | 0.00306042 | 0.81407232 |

| | | | | | | |
|---|---|---|---|---|---|---|
| 238 | Bpifa1 | 156 | 0.32625132 | 3.23E-05 | 0.00308151 | 0.82320716 |
| 1186 | Pgk1 | 159 | -0.323193 | 3.25E-05 | 0.00308151 | 0.82750128 |
| 1124 | Olfr536 | 158 | -0.324075 | 3.27E-05 | 0.00308151 | 0.83237397 |
| 1046 | Naprt | 156 | 0.32602067 | 3.28E-05 | 0.00308151 | 0.83448054 |
| 1004 | Mrc1 | 160 | -0.3220264 | 3.29E-05 | 0.00308151 | 0.83714115 |
| 714 | H2-T23 | 154 | 0.32790279 | 3.30E-05 | 0.00308151 | 0.84151653 |
| 127 | Alkbh6 | 160 | -0.3219179 | 3.31E-05 | 0.00308151 | 0.84256874 |
| 1144 | P2ry10 | 155 | 0.3268289 | 3.32E-05 | 0.00308151 | 0.84433452 |
| 1176 | Pdyn | 156 | 0.32573211 | 3.33E-05 | 0.00308651 | 0.8487895 |
| 1734 | Zfp608 | 155 | -0.3265879 | 3.36E-05 | 0.00310279 | 0.85637076 |
| 1374 | Sema3f | 160 | -0.3212331 | 3.45E-05 | 0.00316821 | 0.8775936 |
| 803 | Itm2a | 160 | -0.3210787 | 3.48E-05 | 0.00318589 | 0.88567853 |
| 1127 | Olfr70 | 156 | 0.32482423 | 3.52E-05 | 0.00320906 | 0.89532756 |
| 498 | Efcab14 | 159 | -0.3217619 | 3.54E-05 | 0.00321713 | 0.90079704 |
| 366 | Cln3 | 156 | 0.32438833 | 3.61E-05 | 0.00326874 | 0.91851518 |
| 1479 | Sox7 | 158 | -0.3219126 | 3.71E-05 | 0.00335131 | 0.94574332 |
| 1304 | Rfx5 | 158 | 0.3218645 | 3.72E-05 | 0.00335131 | 0.94842115 |
| 1205 | Pla2g7 | 160 | -0.3198429 | 3.74E-05 | 0.00335548 | 0.9529574 |
| 246 | C1ql4 | 160 | 0.31976346 | 3.76E-05 | 0.00335944 | 0.95744001 |
| 1097 | Nts | 159 | 0.32060016 | 3.79E-05 | 0.00337323 | 0.96474307 |
| 878 | Lingo2 | 160 | -0.3194246 | 3.84E-05 | 0.00340346 | 0.97679352 |
| 713 | H2-T22 | 155 | 0.32411172 | 3.89E-05 | 0.00343652 | 0.98971708 |
| 121 | Aldh1l2 | 154 | 0.3250546 | 3.90E-05 | 0.00343787 | 0.99354453 |
| 1535 | Taok2 | 157 | -0.3218108 | 3.96E-05 | 0.00347523 | 1 |
| 657 | Gm454 | 158 | -0.3204303 | 4.05E-05 | 0.00354528 | 1 |
| 917 | LOC545780 | 158 | 0.32032161 | 4.08E-05 | 0.00355049 | 1 |
| 202 | Atp6v0a1 | 160 | -0.3183556 | 4.09E-05 | 0.00355049 | 1 |
| 641 | Ghitm | 160 | -0.3182454 | 4.11E-05 | 0.00355968 | 1 |
| 712 | H2-M3 | 155 | 0.3230921 | 4.12E-05 | 0.00355968 | 1 |
| 894 | LOC432762 | 159 | 0.31863852 | 4.25E-05 | 0.0036571 | 1 |
| 513 | Elovl1 | 160 | -0.3173613 | 4.33E-05 | 0.00368481 | 1 |
| 1392 | Sgpp1 | 156 | 0.32112732 | 4.36E-05 | 0.00368481 | 1 |
| 1148 | Pacrgl | 156 | 0.32105951 | 4.38E-05 | 0.00368481 | 1 |
| 709 | Gys1 | 154 | 0.32304983 | 4.38E-05 | 0.00368481 | 1 |
| 152 | App | 160 | -0.3171586 | 4.38E-05 | 0.00368481 | 1 |
| 1409 | Slc16a4 | 160 | -0.3171264 | 4.39E-05 | 0.00368481 | 1 |
| 491 | Eaf2 | 156 | 0.32086676 | 4.43E-05 | 0.00368481 | 1 |
| 1140 | Otx2 | 160 | -0.3169665 | 4.43E-05 | 0.00368481 | 1 |
| 1252 | Prpf31 | 115 | 0.37101987 | 4.47E-05 | 0.00368481 | 1 |
| 9 | 1810022K09Rik | 159 | -0.3177713 | 4.47E-05 | 0.00368481 | 1 |
| 925 | Lrp8 | 160 | -0.3167753 | 4.48E-05 | 0.00368481 | 1 |
| 95 | Adamts3 | 160 | -0.3167581 | 4.49E-05 | 0.00368481 | 1 |
| 339 | Chad | 156 | 0.32058876 | 4.50E-05 | 0.00368481 | 1 |
| 616 | Gabrg1 | 159 | 0.31760955 | 4.51E-05 | 0.00368481 | 1 |
| 997 | Mon1b | 158 | -0.3185663 | 4.52E-05 | 0.00368481 | 1 |

| | | | | | | |
|---|---|---|---|---|---|---|
| 1412 | Slc17a2 | 155 | 0.32149782 | 4.52E-05 | 0.00368481 | 1 |
| 1076 | Nipsnap3b | 152 | 0.32444798 | 4.54E-05 | 0.00368481 | 1 |
| 1050 | Nbl1 | 160 | -0.3164923 | 4.56E-05 | 0.00368481 | 1 |
| 1625 | Ttc28 | 159 | -0.3174436 | 4.56E-05 | 0.00368481 | 1 |
| 1680 | Vwa1 | 160 | -0.316209 | 4.63E-05 | 0.00372747 | 1 |
| 1028 | Musk | 156 | -0.3200583 | 4.64E-05 | 0.00372747 | 1 |
| 227 | Bend6 | 155 | -0.320935 | 4.67E-05 | 0.00374042 | 1 |
| 493 | Ebf1 | 158 | -0.3177683 | 4.73E-05 | 0.00376674 | 1 |
| 1454 | Smim3 | 158 | 0.31775897 | 4.73E-05 | 0.00376674 | 1 |
| 1132 | Oprm1 | 157 | -0.3186006 | 4.77E-05 | 0.00376782 | 1 |
| 1325 | Rlbp1 | 152 | 0.32358022 | 4.77E-05 | 0.00376782 | 1 |
| 622 | Gal | 155 | 0.3205368 | 4.78E-05 | 0.00376782 | 1 |
| 596 | Fndc8 | 159 | -0.3165375 | 4.80E-05 | 0.00377634 | 1 |
| 528 | Epb4.1l4a | 158 | -0.3174332 | 4.82E-05 | 0.00377945 | 1 |
| 474 | Dopey1 | 158 | -0.3173005 | 4.86E-05 | 0.0037892 | 1 |
| 403 | Crabp2 | 155 | 0.32022395 | 4.87E-05 | 0.0037892 | 1 |
| 1098 | Nts | 160 | 0.3148103 | 5.03E-05 | 0.00390172 | 1 |
| 1417 | Slc22a20 | 158 | 0.3164072 | 5.12E-05 | 0.0039499 | 1 |
| 499 | Efemp2 | 160 | -0.3143831 | 5.15E-05 | 0.0039499 | 1 |
| 886 | LOC432444 | 159 | -0.315281 | 5.17E-05 | 0.0039499 | 1 |
| 59 | A330041J22Rik | 158 | -0.3162339 | 5.17E-05 | 0.0039499 | 1 |
| 1408 | Slc16a10 | 154 | 0.32014992 | 5.17E-05 | 0.0039499 | 1 |
| 1248 | Prl2c2 | 156 | 0.3181428 | 5.18E-05 | 0.0039499 | 1 |
| 1430 | Slc35b2 | 159 | 0.31513268 | 5.21E-05 | 0.00396199 | 1 |
| 1524 | Sypl | 160 | -0.3140501 | 5.25E-05 | 0.00398049 | 1 |
| 1410 | Slc16a4 | 156 | 0.31715447 | 5.48E-05 | 0.00414257 | 1 |
| 1168 | Pde7b | 159 | 0.31402863 | 5.55E-05 | 0.00418498 | 1 |
| 206 | Atp6v1g2 | 158 | -0.3149229 | 5.57E-05 | 0.00418754 | 1 |
| 1209 | Plekhh3 | 156 | 0.31670682 | 5.62E-05 | 0.00421231 | 1 |
| 263 | Car9 | 158 | -0.3144676 | 5.72E-05 | 0.0042731 | 1 |
| 680 | Gpr149 | 158 | 0.31436873 | 5.75E-05 | 0.0042848 | 1 |
| 490 | E2f6 | 160 | -0.3123796 | 5.78E-05 | 0.0042941 | 1 |
| 729 | Hdgf | 160 | -0.3121477 | 5.86E-05 | 0.00433913 | 1 |
| 1736 | Zfp639 | 145 | -0.3271101 | 5.92E-05 | 0.00437117 | 1 |
| 1372 | Sds | 149 | 0.32266682 | 5.99E-05 | 0.00440537 | 1 |
| 681 | Gpr151 | 156 | -0.3153304 | 6.08E-05 | 0.00446362 | 1 |
| 885 | LOC381597 | 159 | -0.3123016 | 6.13E-05 | 0.00448816 | 1 |
| 1740 | Zfp763 | 152 | 0.31870645 | 6.29E-05 | 0.00458852 | 1 |
| 234 | Bmp5 | 160 | -0.3107524 | 6.35E-05 | 0.0046202 | 1 |
| 1606 | Tor1b | 156 | 0.31433888 | 6.43E-05 | 0.00466781 | 1 |
| 468 | Dnah6 | 159 | -0.3113105 | 6.49E-05 | 0.00467041 | 1 |
| 718 | Haus5 | 158 | -0.3122566 | 6.49E-05 | 0.00467041 | 1 |
| 1580 | Tmem106c | 155 | 0.31500574 | 6.55E-05 | 0.00467041 | 1 |
| 27 | 2810043G22Rik | 156 | -0.3139423 | 6.58E-05 | 0.00467041 | 1 |
| 839 | Klhdc7a | 158 | -0.3120122 | 6.58E-05 | 0.00467041 | 1 |

| | | | | | | |
|---|---|---|---|---|---|---|
| 855 | Lair1 | 155 | 0.31488465 | 6.59E-05 | 0.00467041 | 1 |
| 1041 | Nadsyn1 | 155 | 0.31487944 | 6.59E-05 | 0.00467041 | 1 |
| 87 | Acp2 | 156 | 0.31386504 | 6.61E-05 | 0.00467041 | 1 |
| 1365 | Scaf11 | 156 | -0.3137997 | 6.63E-05 | 0.00467041 | 1 |
| 1648 | Ufsp1 | 159 | 0.31087431 | 6.65E-05 | 0.00467041 | 1 |
| 1024 | Mucl1 | 155 | 0.31465314 | 6.68E-05 | 0.00467041 | 1 |
| 132 | Amd1 | 158 | -0.311721 | 6.69E-05 | 0.00467041 | 1 |
| 1190 | Pgr15l | 160 | 0.30981926 | 6.70E-05 | 0.00467041 | 1 |
| 425 | Cyhr1 | 160 | -0.3098139 | 6.70E-05 | 0.00467041 | 1 |
| 1439 | Slc5a11 | 151 | 0.31855068 | 6.71E-05 | 0.00467041 | 1 |
| 1380 | Serpinb12 | 159 | -0.3104776 | 6.81E-05 | 0.00471811 | 1 |
| 722 | Hbb | 147 | -0.3224106 | 6.82E-05 | 0.00471811 | 1 |
| 1719 | Zc3h12c | 156 | -0.3132569 | 6.84E-05 | 0.00471982 | 1 |
| 584 | Fbxo8 | 155 | 0.31402379 | 6.92E-05 | 0.00475184 | 1 |
| 1639 | Uba2 | 159 | -0.3101768 | 6.92E-05 | 0.00475184 | 1 |
| 1339 | Rpp25 | 160 | -0.3087661 | 7.11E-05 | 0.00486838 | 1 |
| 311 | Cdh5 | 158 | -0.3105732 | 7.14E-05 | 0.00487558 | 1 |
| 1556 | Tdgf1 | 155 | 0.31324509 | 7.23E-05 | 0.00492123 | 1 |
| 503 | Ei24 | 160 | -0.3083296 | 7.29E-05 | 0.00492601 | 1 |
| 268 | Cbln2 | 160 | -0.3082921 | 7.31E-05 | 0.00492601 | 1 |
| 1246 | Prkab1 | 152 | 0.31597549 | 7.32E-05 | 0.00492601 | 1 |
| 1271 | Qpct | 156 | 0.31202599 | 7.33E-05 | 0.00492601 | 1 |
| 458 | Diras1 | 160 | -0.3082031 | 7.34E-05 | 0.00492601 | 1 |
| 1260 | Psmc6 | 158 | -0.3100616 | 7.35E-05 | 0.00492601 | 1 |
| 1249 | Prl4a1 | 156 | 0.31164699 | 7.49E-05 | 0.00498492 | 1 |
| 1354 | Rwdd2b | 155 | 0.3125863 | 7.50E-05 | 0.00498492 | 1 |
| 774 | Igf2bp3 | 155 | 0.31255648 | 7.51E-05 | 0.00498492 | 1 |
| 1438 | Slc51a | 155 | 0.31254341 | 7.52E-05 | 0.00498492 | 1 |
| 1502 | Stat5b | 156 | 0.31149303 | 7.55E-05 | 0.00499486 | 1 |
| 762 | Htr6 | 147 | 0.32048241 | 7.58E-05 | 0.0049991 | 1 |
| 821 | Kcnj15 | 158 | 0.30944242 | 7.61E-05 | 0.00500863 | 1 |
| 1693 | Wls | 160 | -0.3073202 | 7.72E-05 | 0.00506479 | 1 |
| 959 | Mb | 159 | 0.30819177 | 7.75E-05 | 0.00506479 | 1 |
| 830 | Kdf1 | 156 | 0.31101392 | 7.76E-05 | 0.00506479 | 1 |
| 1086 | Npr1 | 159 | -0.3079474 | 7.85E-05 | 0.00511456 | 1 |
| 137 | Antxr1 | 158 | -0.308833 | 7.88E-05 | 0.00511709 | 1 |
| 1588 | Tmem161b | 156 | 0.31052964 | 7.97E-05 | 0.00516394 | 1 |
| 1023 | Muc13 | 156 | 0.31026691 | 8.09E-05 | 0.00522015 | 1 |
| 782 | Il4 | 155 | 0.3112025 | 8.10E-05 | 0.00522015 | 1 |
| 1114 | Obox5 | 152 | 0.31411236 | 8.12E-05 | 0.00522015 | 1 |
| 1712 | Ywhag | 160 | -0.3062119 | 8.22E-05 | 0.00526098 | 1 |
| 387 | Col9a1 | 160 | -0.3062008 | 8.22E-05 | 0.00526098 | 1 |
| 1020 | Mtss1 | 160 | -0.3061002 | 8.27E-05 | 0.00527742 | 1 |
| 177 | Armc4 | 159 | -0.3069592 | 8.30E-05 | 0.00527742 | 1 |
| 675 | Gpnmb | 160 | -0.306012 | 8.31E-05 | 0.00527742 | 1 |

| | | | | | | |
|---|---|---|---|---|---|---|
| 1652 | Ung | 156 | 0.30956954 | 8.41E-05 | 0.00532572 | 1 |
| 1514 | Susd2 | 153 | -0.3123722 | 8.46E-05 | 0.00534574 | 1 |
| 533 | Epha10 | 160 | -0.3055708 | 8.52E-05 | 0.0053698 | 1 |
| 1120 | Olfr1387 | 158 | -0.3072889 | 8.59E-05 | 0.00540018 | 1 |
| 1620 | Tspan1 | 155 | 0.310065 | 8.63E-05 | 0.00541161 | 1 |
| 1270 | Pyurf | 146 | 0.31902287 | 8.68E-05 | 0.00543324 | 1 |
| 151 | Apoc2 | 156 | 0.30882247 | 8.76E-05 | 0.00546971 | 1 |
| 1564 | Tex22 | 159 | 0.30586528 | 8.83E-05 | 0.00549541 | 1 |
| 1533 | Tal1 | 160 | -0.3048007 | 8.90E-05 | 0.00552223 | 1 |
| 996 | Mob3b | 160 | -0.3047656 | 8.91E-05 | 0.00552223 | 1 |
| 1669 | Vdac3 | 160 | -0.3043782 | 9.11E-05 | 0.00562961 | 1 |
| 857 | Lao1 | 152 | 0.31161437 | 9.31E-05 | 0.0057317 | 1 |
| 870 | Lgals1 | 160 | -0.30397 | 9.32E-05 | 0.0057317 | 1 |
| 90 | Actn2 | 156 | -0.3076667 | 9.34E-05 | 0.00573273 | 1 |
| 1671 | Vmac | 158 | -0.3052974 | 9.60E-05 | 0.0058669 | 1 |
| 578 | Fbp2 | 155 | 0.30811651 | 9.61E-05 | 0.0058669 | 1 |
| 477 | Dppa4 | 156 | 0.30705677 | 9.66E-05 | 0.00588639 | 1 |
| 999 | Mpg | 156 | 0.306922 | 9.73E-05 | 0.00591362 | 1 |
| 191 | Atf5 | 159 | -0.3040618 | 9.76E-05 | 0.00591362 | 1 |
| 530 | Epgn | 160 | -0.3031091 | 9.78E-05 | 0.00591362 | 1 |
| 758 | Hsp90b1 | 156 | -0.3064638 | 9.98E-05 | 0.00602411 | 1 |
| 434 | Dand5 | 159 | -0.3035104 | 0.00010063 | 0.00604617 | 1 |
| 617 | Gabrp | 160 | 0.302583 | 0.00010067 | 0.00604617 | 1 |
| 1290 | Rbm19 | 129 | 0.3355203 | 0.0001015 | 0.0060734 | 1 |
| 874 | Lhx6 | 153 | 0.30894202 | 0.0001021 | 0.0060734 | 1 |
| 1217 | Plxnb3 | 155 | -0.307005 | 0.00010212 | 0.0060734 | 1 |
| 716 | Hadha | 151 | -0.310838 | 0.00010251 | 0.0060734 | 1 |
| 106 | Agtrap | 157 | -0.3050324 | 0.00010254 | 0.0060734 | 1 |
| 1614 | Trex2 | 159 | -0.3031675 | 0.00010256 | 0.0060734 | 1 |
| 1241 | Pramel4 | 159 | -0.3031143 | 0.00010286 | 0.00607719 | 1 |
| 1489 | Spryd4 | 155 | 0.30659321 | 0.00010445 | 0.00615657 | 1 |
| 1282 | Ralgps2 | 156 | 0.30557211 | 0.00010484 | 0.0061657 | 1 |
| 7 | 1700094D03Rik | 150 | -0.3111973 | 0.00010611 | 0.00622555 | 1 |
| 1286 | Rax | 159 | 0.30239684 | 0.00010702 | 0.0062649 | 1 |
| 1035 | Myoz1 | 149 | 0.31195283 | 0.00010755 | 0.00628117 | 1 |
| 942 | Mafg | 156 | -0.3050648 | 0.0001078 | 0.0062814 | 1 |
| 1320 | Riok1 | 155 | 0.30591439 | 0.00010839 | 0.0063016 | 1 |
| 1468 | Snx20 | 158 | 0.30286793 | 0.00010973 | 0.00636468 | 1 |
| 767 | Id3 | 160 | -0.300972 | 0.00011008 | 0.00637037 | 1 |
| 1653 | Usb1 | 156 | 0.3045034 | 0.00011116 | 0.00641843 | 1 |
| 1582 | Tmem123 | 155 | 0.30528234 | 0.00011219 | 0.00645581 | 1 |
| 1342 | Rps21 | 155 | 0.30526237 | 0.00011231 | 0.00645581 | 1 |
| 1243 | Prcp | 160 | -0.3005458 | 0.0001127 | 0.00646306 | 1 |
| 689 | Gpt | 156 | 0.30420255 | 0.000113 | 0.00646306 | 1 |
| 1147 | Pabpc4 | 156 | 0.3041699 | 0.0001132 | 0.00646306 | 1 |

| | | | | | | |
|---|---|---|---|---|---|---|
| 1670 | Vipr2 | 156 | 0.30404004 | 0.000114 | 0.00649444 | 1 |
| 290 | Ccnl2 | 160 | -0.3000613 | 0.00011575 | 0.00657034 | 1 |
| 1181 | Peli1 | 156 | 0.30374489 | 0.00011585 | 0.00657034 | 1 |
| 18 | 2310039H08Rik | 158 | 0.30145357 | 0.00011857 | 0.00670971 | 1 |
| 888 | LOC432463 | 159 | -0.3003916 | 0.0001195 | 0.00674729 | 1 |
| 131 | Amacr | 155 | 0.30404086 | 0.00012002 | 0.00676123 | 1 |
| 198 | Atp5a1 | 160 | -0.2988177 | 0.00012393 | 0.0069576 | 1 |
| 518 | Emc3 | 156 | 0.30248724 | 0.00012405 | 0.0069576 | 1 |
| 1683 | Vwf | 159 | -0.2996108 | 0.00012472 | 0.00697997 | 1 |
| 253 | Caap1 | 158 | 0.30044696 | 0.00012527 | 0.00699542 | 1 |
| 691 | Grap2 | 156 | 0.30214225 | 0.00012639 | 0.00704235 | 1 |
| 313 | Cdhr4 | 159 | -0.2993058 | 0.00012682 | 0.00705075 | 1 |
| 755 | Hsd3b7 | 155 | 0.30265791 | 0.00012933 | 0.0071749 | 1 |
| 1743 | Zfp91 | 159 | -0.2986483 | 0.00013144 | 0.00727211 | 1 |
| 99 | Adi1 | 155 | 0.30232398 | 0.00013168 | 0.00727211 | 1 |
| 898 | LOC433071 | 155 | -0.3022519 | 0.00013219 | 0.00727211 | 1 |
| 1328 | Rnd2 | 160 | -0.2976136 | 0.00013237 | 0.00727211 | 1 |
| 1695 | Wrap73 | 160 | 0.29759422 | 0.00013251 | 0.00727211 | 1 |
| 686 | Gpr82 | 153 | -0.3037931 | 0.0001348 | 0.00733263 | 1 |
| 1729 | Zfp185 | 158 | -0.2990983 | 0.0001348 | 0.00733263 | 1 |
| 525 | Enpp6 | 160 | -0.2972596 | 0.00013495 | 0.00733263 | 1 |
| 840 | Klhl10 | 159 | 0.29815892 | 0.00013499 | 0.00733263 | 1 |
| 975 | Mettl21a | 155 | 0.3018537 | 0.00013505 | 0.00733263 | 1 |
| 1155 | Papola | 160 | -0.2970879 | 0.00013622 | 0.00737075 | 1 |
| 927 | Lrrcc1 | 156 | -0.3007147 | 0.00013652 | 0.00737075 | 1 |
| 912 | LOC434786 | 159 | -0.2979377 | 0.00013662 | 0.00737075 | 1 |
| 1215 | Pltp | 156 | -0.3005855 | 0.00013747 | 0.00738912 | 1 |
| 462 | Dlx6os1 | 158 | 0.29872613 | 0.00013754 | 0.00738912 | 1 |
| 465 | Dmxl1 | 159 | -0.2976258 | 0.00013896 | 0.00743359 | 1 |
| 1301 | Rfk | 152 | 0.30417407 | 0.00013906 | 0.00743359 | 1 |
| 104 | Ago2 | 157 | -0.2994186 | 0.00013925 | 0.00743359 | 1 |
| 1718 | Zc2hc1b | 160 | 0.29663271 | 0.00013964 | 0.00743879 | 1 |
| 447 | Dennd4c | 158 | -0.2983048 | 0.00014072 | 0.00748058 | 1 |
| 1112 | Oasl2 | 156 | 0.30003015 | 0.00014164 | 0.00751408 | 1 |
| 357 | Clcc1 | 155 | 0.30080836 | 0.00014285 | 0.00756221 | 1 |
| 335 | Cfap43 | 160 | -0.2961084 | 0.00014367 | 0.00759023 | 1 |
| 1236 | Ppp1r16a | 159 | -0.2968997 | 0.00014453 | 0.00761987 | 1 |
| 497 | Efcab12 | 160 | -0.2958842 | 0.00014543 | 0.0076514 | 1 |
| 1602 | Tnfrsf17 | 156 | 0.29945927 | 0.00014605 | 0.00766805 | 1 |
| 602 | Fpr-rs4 | 159 | -0.2966356 | 0.00014661 | 0.00768182 | 1 |
| 724 | Hbb-b1 | 158 | -0.2973842 | 0.00014788 | 0.00772357 | 1 |
| 934 | Lyve1 | 155 | 0.30012108 | 0.00014819 | 0.00772357 | 1 |
| 837 | Klf7 | 160 | -0.2954947 | 0.00014853 | 0.00772357 | 1 |
| 1440 | Slc5a2 | 156 | 0.29913331 | 0.00014862 | 0.00772357 | 1 |
| 1471 | Snx8 | 158 | 0.29712987 | 0.00014992 | 0.00775305 | 1 |

| | | | | | | |
|---|---|---|---|---|---|---|
| 220 | Bcar3 | 160 | -0.2953073 | 0.00015005 | 0.00775305 | 1 |
| 1219 | Pml | 155 | 0.29988137 | 0.0001501 | 0.00775305 | 1 |
| 1303 | Rfx1 | 159 | 0.29604251 | 0.00015138 | 0.00780337 | 1 |
| 979 | Mgat1 | 155 | 0.29957589 | 0.00015257 | 0.0078486 | 1 |
| 176 | Armc3 | 159 | -0.2957794 | 0.00015355 | 0.00788295 | 1 |
| 1102 | Nudt1 | 157 | 0.29721452 | 0.00015676 | 0.00802152 | 1 |
| 736 | Hibadh | 156 | 0.29812248 | 0.00015688 | 0.00802152 | 1 |
| 1122 | Olfr371 | 159 | -0.2948825 | 0.00016114 | 0.00822296 | 1 |
| 599 | Foxj1 | 156 | -0.2975365 | 0.00016186 | 0.00822569 | 1 |
| 1594 | Tmem229b | 158 | -0.2956583 | 0.00016225 | 0.00822569 | 1 |
| 1637 | Uaca | 158 | -0.2956104 | 0.00016267 | 0.00822569 | 1 |
| 674 | Gpc3 | 160 | 0.2938104 | 0.00016268 | 0.00822569 | 1 |
| 224 | Bcl2l10 | 157 | 0.29650593 | 0.00016281 | 0.00822569 | 1 |
| 1137 | Osbpl7 | 126 | -0.3296751 | 0.00016347 | 0.00824271 | 1 |
| 1608 | Tox | 159 | -0.2943789 | 0.00016555 | 0.00830495 | 1 |
| 1550 | Tcerg1l | 160 | 0.29347922 | 0.0001656 | 0.00830495 | 1 |
| 1175 | Pdxk | 160 | -0.2934702 | 0.00016568 | 0.00830495 | 1 |
| 1593 | Tmem229a | 159 | -0.2942749 | 0.00016648 | 0.00832858 | 1 |
| 1298 | Rdh7 | 152 | 0.30072272 | 0.0001669 | 0.00833332 | 1 |
| 606 | Fstl5 | 159 | 0.29404519 | 0.00016854 | 0.00839765 | 1 |
| 1507 | Stx3 | 158 | -0.2949126 | 0.00016885 | 0.00839765 | 1 |
| 662 | Gm5607 | 160 | 0.29304245 | 0.00016953 | 0.00841518 | 1 |
| 1531 | Sytl5 | 160 | 0.29292813 | 0.00017058 | 0.00844342 | 1 |
| 507 | Eif4a3 | 160 | -0.2929075 | 0.00017077 | 0.00844342 | 1 |
| 889 | LOC432560 | 158 | 0.29450174 | 0.00017259 | 0.00851724 | 1 |
| 123 | Aldh9a1 | 155 | 0.29697824 | 0.00017512 | 0.00862535 | 1 |
| 778 | Igsf3 | 160 | -0.2922677 | 0.00017672 | 0.00864305 | 1 |
| 1563 | Tex19.1 | 155 | 0.29680517 | 0.00017673 | 0.00864305 | 1 |
| 1167 | Pde4dip | 160 | -0.2922571 | 0.00017682 | 0.00864305 | 1 |
| 279 | Ccdc23 | 156 | 0.29586893 | 0.00017684 | 0.00864305 | 1 |
| 815 | Kcna2 | 147 | -0.3044708 | 0.00017725 | 0.00864632 | 1 |
| 306 | Cdca7 | 160 | -0.2921175 | 0.00017815 | 0.00866601 | 1 |
| 1061 | Nefl | 160 | -0.2920382 | 0.0001789 | 0.00866601 | 1 |
| 1592 | Tmem219 | 152 | 0.29937528 | 0.00017912 | 0.00866601 | 1 |
| 164 | Arg1 | 152 | 0.2993546 | 0.00017931 | 0.00866601 | 1 |
| 496 | Ecel1 | 160 | 0.29199168 | 0.00017935 | 0.00866601 | 1 |
| 178 | Arr3 | 155 | 0.2963525 | 0.000181 | 0.00872913 | 1 |
| 1395 | Shh | 160 | 0.29178294 | 0.00018136 | 0.00873011 | 1 |
| 259 | Calcr | 160 | 0.29168987 | 0.00018227 | 0.00875421 | 1 |
| 1335 | Rpap2 | 156 | 0.29526769 | 0.00018255 | 0.00875421 | 1 |
| 798 | Isl1 | 155 | 0.29611106 | 0.00018332 | 0.00877439 | 1 |
| 107 | Agxt2 | 159 | -0.2922911 | 0.00018509 | 0.00884242 | 1 |
| 1203 | Pla2g10 | 149 | 0.301515 | 0.00018637 | 0.00888707 | 1 |
| 1665 | Vat1 | 152 | 0.29850507 | 0.00018744 | 0.00889743 | 1 |
| 1749 | Zim1 | 160 | 0.29115807 | 0.00018751 | 0.00889743 | 1 |

| | | | | | | |
|---|---|---|---|---|---|---|
| 1182 | Peli3 | 156 | 0.2947472 | 0.00018763 | 0.00889743 | 1 |
| 54 | 9030407P20Rik | 158 | -0.2928189 | 0.00018874 | 0.00893333 | 1 |
| 1308 | Rgs5 | 160 | -0.2909576 | 0.00018953 | 0.0089402 | 1 |
| 783 | Il4ra | 160 | -0.2909514 | 0.00018959 | 0.0089402 | 1 |
| 1717 | Zbtb4 | 149 | -0.300857 | 0.00019281 | 0.00907322 | 1 |
| 922 | Lpar1 | 159 | -0.291491 | 0.00019312 | 0.00907322 | 1 |
| 212 | B630019K06Rik | 159 | 0.2914189 | 0.00019386 | 0.00909121 | 1 |
| 273 | Ccdc108 | 156 | -0.2940258 | 0.0001949 | 0.00909267 | 1 |
| 1391 | Sgpl1 | 156 | -0.2940205 | 0.00019495 | 0.00909267 | 1 |
| 551 | Fabp7 | 160 | -0.290426 | 0.00019497 | 0.00909267 | 1 |
| 862 | Ldlrad2 | 158 | -0.2921443 | 0.0001956 | 0.00909881 | 1 |
| 169 | Arhgap5 | 160 | -0.2903445 | 0.00019581 | 0.00909881 | 1 |
| 921 | Lox | 155 | 0.29481579 | 0.00019622 | 0.00910134 | 1 |
| 594 | Fibp | 152 | 0.29746557 | 0.00019786 | 0.00916044 | 1 |
| 46 | 5430435G22Rik | 156 | 0.29367162 | 0.00019856 | 0.00917622 | 1 |
| 1009 | Mrpl4 | 156 | -0.2935193 | 0.00020015 | 0.00923312 | 1 |
| 261 | Calm1 | 160 | -0.2898541 | 0.00020098 | 0.00924744 | 1 |
| 68 | Aar2 | 158 | 0.29161106 | 0.00020119 | 0.00924744 | 1 |
| 1420 | Slc25a24 | 156 | 0.29328311 | 0.00020265 | 0.00928596 | 1 |
| 1081 | Nmb | 158 | 0.29142763 | 0.00020314 | 0.00928596 | 1 |
| 568 | Fam89b | 157 | 0.29228012 | 0.00020365 | 0.00928596 | 1 |
| 1317 | Rhpn1 | 159 | -0.2904772 | 0.00020377 | 0.00928596 | 1 |
| 69 | Abca13 | 159 | -0.2904701 | 0.00020385 | 0.00928596 | 1 |
| 1083 | Nop9 | 160 | 0.28953166 | 0.00020444 | 0.00929634 | 1 |
| 122 | Aldh3b1 | 158 | 0.29121368 | 0.00020545 | 0.00932464 | 1 |
| 974 | Mettl14 | 152 | 0.2966914 | 0.00020596 | 0.00932464 | 1 |
| 1506 | Strbp | 158 | -0.2911476 | 0.00020616 | 0.00932464 | 1 |
| 1488 | Sphkap | 160 | -0.2892702 | 0.00020729 | 0.00935911 | 1 |
| 660 | Gm5126 | 159 | -0.2900814 | 0.00020808 | 0.00937793 | 1 |
| 725 | Hbb-b2 | 158 | -0.2907871 | 0.00021011 | 0.0094529 | 1 |
| 457 | Dip2a | 160 | -0.2887905 | 0.00021262 | 0.00954885 | 1 |
| 398 | Cpe | 160 | -0.2886857 | 0.0002138 | 0.00957011 | 1 |
| 759 | Htatsf1 | 158 | -0.2904521 | 0.00021385 | 0.00957011 | 1 |
| 846 | Kmt2a | 160 | -0.2885512 | 0.00021533 | 0.00961938 | 1 |
| 321 | Cds1 | 160 | -0.2883582 | 0.00021753 | 0.00967998 | 1 |
| 995 | Mob3b | 155 | 0.29282411 | 0.00021773 | 0.00967998 | 1 |
| 1568 | Thbs2 | 160 | -0.2883328 | 0.00021782 | 0.00967998 | 1 |
| 1399 | Sigirr | 158 | 0.29004909 | 0.00021842 | 0.00968956 | 1 |
| 486 | Dynlrb2 | 160 | -0.2881599 | 0.00021982 | 0.00973465 | 1 |
| 626 | Galnt1 | 153 | 0.29424952 | 0.00022259 | 0.00983533 | 1 |
| 983 | Mipol1 | 156 | -0.2914636 | 0.00022286 | 0.00983533 | 1 |
| 760 | Htr2c | 160 | -0.2877985 | 0.00022404 | 0.00986256 | 1 |
| 439 | Dclre1b | 156 | 0.29134411 | 0.00022425 | 0.00986256 | 1 |
| 1523 | Syp | 159 | -0.2885103 | 0.00022601 | 0.00990968 | 1 |
| 1194 | Pidd1 | 152 | -0.2948802 | 0.00022614 | 0.00990968 | 1 |

| | | | | | | |
|---|---|---|---|---|---|---|
| 764 | Hyal1 | 152 | 0.29485018 | 0.00022649 | 0.00990968 | 1 |
| 392 | Coq7 | 155 | 0.29194525 | 0.0002279 | 0.00994151 | 1 |
| 1674 | Vps13b | 159 | -0.2882971 | 0.00022856 | 0.00994151 | 1 |
| 715 | Hacl1 | 156 | 0.29095734 | 0.00022881 | 0.00994151 | 1 |
| 1356 | Rxrg | 160 | -0.2873712 | 0.00022914 | 0.00994151 | 1 |
| 60 | A630043P06 | 160 | -0.287337 | 0.00022955 | 0.00994151 | 1 |
| 356 | Clasp1 | 159 | -0.2881872 | 0.00022988 | 0.00994151 | 1 |
| 40 | 4930588G17Rik | 159 | -0.2881568 | 0.00023024 | 0.00994151 | 1 |
| 1382 | Serpinb9e | 155 | 0.2917196 | 0.00023058 | 0.00994151 | 1 |
| 1104 | Nwd1 | 160 | -0.2872389 | 0.00023073 | 0.00994151 | 1 |
| 520 | Enkur | 159 | -0.2879166 | 0.00023316 | 0.01002105 | 1 |
| 557 | Fam187a | 159 | -0.2878994 | 0.00023337 | 0.01002105 | 1 |
| 652 | Gm1335 | 160 | -0.2869749 | 0.00023396 | 0.01002944 | 1 |
| 71 | Abcc4 | 160 | -0.2867242 | 0.00023705 | 0.01014511 | 1 |
| 612 | G6pc | 152 | 0.29387011 | 0.00023818 | 0.01017632 | 1 |
| 351 | Ciao1 | 160 | 0.28653509 | 0.00023942 | 0.01019204 | 1 |
| 1311 | Rhebl1 | 156 | 0.29007352 | 0.00023955 | 0.01019204 | 1 |
| 615 | Gabbr2 | 155 | -0.2909657 | 0.00023975 | 0.01019204 | 1 |
| 210 | B3gnt8 | 155 | 0.29084982 | 0.00024119 | 0.01023617 | 1 |
| 92 | Adamts1 | 158 | 0.28806009 | 0.00024235 | 0.01026801 | 1 |
| 226 | Becn1 | 157 | -0.288861 | 0.00024348 | 0.01028731 | 1 |
| 1366 | Scamp4 | 156 | 0.28974946 | 0.00024361 | 0.01028731 | 1 |
| 1753 | Zscan26 | 159 | -0.2870118 | 0.00024445 | 0.01030579 | 1 |
| 1158 | Pcdh8 | 159 | 0.28688322 | 0.0002461 | 0.01035579 | 1 |
| 1329 | Rnf11 | 160 | -0.285982 | 0.00024645 | 0.01035579 | 1 |
| 1373 | Sectm1a | 153 | 0.29221029 | 0.00024722 | 0.0103709 | 1 |
| 784 | Impa2 | 154 | 0.29095428 | 0.00025147 | 0.01053216 | 1 |
| 643 | Gja1 | 160 | -0.2853265 | 0.00025503 | 0.01064159 | 1 |
| 318 | Cdk8 | 160 | -0.285324 | 0.00025507 | 0.01064159 | 1 |
| 935 | Lyve1 | 160 | -0.2853035 | 0.00025534 | 0.01064159 | 1 |
| 522 | Enpp2 | 160 | -0.285165 | 0.00025719 | 0.01070121 | 1 |
| 1351 | Rsu1 | 158 | 0.28684909 | 0.00025808 | 0.01072073 | 1 |
| 345 | Chrna10 | 155 | 0.28946419 | 0.00025904 | 0.01073052 | 1 |
| 1033 | Myl6b | 155 | 0.289455 | 0.00025916 | 0.01073052 | 1 |
| 1579 | Tmem106b | 160 | -0.2848846 | 0.00026098 | 0.0107882 | 1 |
| 362 | Clic1 | 160 | -0.2847254 | 0.00026315 | 0.01085381 | 1 |
| 1493 | Sri | 160 | -0.2847059 | 0.00026342 | 0.01085381 | 1 |
| 235 | Bmp7 | 160 | -0.2846002 | 0.00026487 | 0.01089308 | 1 |
| 640 | Ggt1 | 155 | 0.28898579 | 0.00026548 | 0.01089308 | 1 |
| 113 | AK129341 | 156 | -0.2880706 | 0.00026565 | 0.01089308 | 1 |
| 1446 | Slc9a8 | 158 | 0.28621696 | 0.00026667 | 0.01091704 | 1 |
| 1283 | Ran | 160 | -0.2844301 | 0.00026722 | 0.01091704 | 1 |
| 400 | Cpt1b | 159 | 0.28526116 | 0.00026775 | 0.01091704 | 1 |
| 186 | Asns | 160 | -0.2843777 | 0.00026795 | 0.01091704 | 1 |
| 1452 | Smg6 | 156 | -0.2874413 | 0.00027438 | 0.01110393 | 1 |

| | | | | | | |
|---|---|---|---|---|---|---|
| 669 | Gnat1 | 160 | -0.2839024 | 0.00027465 | 0.01110393 | 1 |
| 991 | Mmp3 | 155 | 0.28829542 | 0.00027504 | 0.01110393 | 1 |
| 1220 | Pnoc | 156 | 0.28739457 | 0.00027504 | 0.01110393 | 1 |
| 1043 | Nans | 158 | 0.28560211 | 0.00027527 | 0.01110393 | 1 |
| 1701 | Xlr4a | 156 | -0.2873447 | 0.00027575 | 0.01110393 | 1 |
| 1315 | Rhox5 | 158 | -0.285552 | 0.00027599 | 0.01110393 | 1 |
| 453 | Dhcr7 | 156 | 0.2872726 | 0.00027677 | 0.01110393 | 1 |
| 891 | LOC432648 | 156 | -0.2872648 | 0.00027688 | 0.01110393 | 1 |
| 1450 | Smc5 | 154 | 0.28907209 | 0.0002769 | 0.01110393 | 1 |
| 1710 | Ywhab | 158 | 0.28541029 | 0.00027801 | 0.01111445 | 1 |
| 162 | Araf | 157 | 0.28629193 | 0.00027804 | 0.01111445 | 1 |
| 1459 | Snap25 | 160 | -0.2835967 | 0.00027904 | 0.01113714 | 1 |
| 820 | Kcnip1 | 160 | -0.2835386 | 0.00027988 | 0.01115325 | 1 |
| 197 | Atp13a5 | 160 | -0.2833343 | 0.00028286 | 0.01125428 | 1 |
| 1500 | Stard10 | 160 | -0.2832135 | 0.00028463 | 0.01130724 | 1 |
| 601 | Fpgs | 156 | 0.28647419 | 0.00028832 | 0.01143577 | 1 |
| 692 | Grin2d | 160 | -0.2827346 | 0.00029177 | 0.01155471 | 1 |
| 1566 | Tgfbi | 158 | -0.2843794 | 0.00029316 | 0.01159164 | 1 |
| 1368 | Scgb2b27 | 158 | -0.2841621 | 0.00029645 | 0.01167253 | 1 |
| 646 | Gldc | 160 | -0.2823741 | 0.00029725 | 0.01167253 | 1 |
| 1739 | Zfp758 | 158 | 0.28409279 | 0.00029751 | 0.01167253 | 1 |
| 707 | Gtf2h2 | 152 | 0.28947767 | 0.00029778 | 0.01167253 | 1 |
| 281 | Ccdc60 | 159 | -0.2830994 | 0.00029938 | 0.01167253 | 1 |
| 1305 | Rgcc | 159 | -0.2830854 | 0.00029959 | 0.01167253 | 1 |
| 221 | Bcar3 | 158 | -0.2839415 | 0.00029982 | 0.01167253 | 1 |
| 754 | Hsd3b5 | 156 | 0.28565868 | 0.00030058 | 0.01167253 | 1 |
| 1703 | Xxylt1 | 158 | 0.28388855 | 0.00030064 | 0.01167253 | 1 |
| 1326 | Rln3 | 160 | -0.2821074 | 0.00030137 | 0.01167253 | 1 |
| 1560 | Tenc1 | 156 | -0.2855889 | 0.00030165 | 0.01167253 | 1 |
| 1117 | Ogt | 160 | -0.2820768 | 0.00030185 | 0.01167253 | 1 |
| 1143 | P2rx3 | 160 | 0.28206513 | 0.00030203 | 0.01167253 | 1 |
| 1730 | Zfp263 | 156 | 0.28554296 | 0.00030236 | 0.01167253 | 1 |
| 309 | Cdh24 | 156 | -0.2855361 | 0.00030246 | 0.01167253 | 1 |
| 1094 | Nt5dc1 | 159 | 0.28289494 | 0.00030254 | 0.01167253 | 1 |
| 807 | Jade1 | 152 | 0.28902567 | 0.00030464 | 0.01173498 | 1 |
| 792 | Irx1 | 153 | -0.2880767 | 0.00030508 | 0.01173498 | 1 |
| 195 | Atl3 | 160 | -0.2814222 | 0.00031219 | 0.01199045 | 1 |
| 470 | Dnajc24 | 150 | 0.29029687 | 0.00031387 | 0.01203683 | 1 |
| 1183 | Penk | 159 | 0.28195153 | 0.00031754 | 0.01214331 | 1 |
| 1139 | Ostn | 160 | -0.2810878 | 0.0003176 | 0.01214331 | 1 |
| 379 | Cntrl | 156 | -0.2844818 | 0.00031912 | 0.0121832 | 1 |
| 985 | Mitf | 156 | -0.2841752 | 0.00032413 | 0.01235565 | 1 |
| 1055 | Ndrg2 | 158 | 0.28230679 | 0.00032596 | 0.01240685 | 1 |
| 426 | Cyp1b1 | 155 | 0.28490348 | 0.00032681 | 0.0124207 | 1 |
| 632 | Gata2 | 160 | -0.2803464 | 0.00032991 | 0.01251987 | 1 |

| | | | | | | |
|---|---|---|---|---|---|---|
| 334 | Cfap43 | 159 | -0.2811093 | 0.00033152 | 0.01252877 | 1 |
| 1429 | Slc35a2 | 160 | 0.28025084 | 0.00033153 | 0.01252877 | 1 |
| 1559 | Tekt2 | 143 | 0.2959887 | 0.00033168 | 0.01252877 | 1 |
| 572 | Fasl | 152 | 0.28730319 | 0.00033215 | 0.01252877 | 1 |
| 544 | Exosc3 | 155 | 0.28455537 | 0.0003326 | 0.01252877 | 1 |
| 1478 | Sox14 | 160 | -0.2801545 | 0.00033317 | 0.01253141 | 1 |
| 1273 | R3hcc1 | 158 | -0.281787 | 0.0003347 | 0.0125705 | 1 |
| 899 | LOC433088 | 160 | 0.27999432 | 0.00033591 | 0.01259733 | 1 |
| 607 | Ftcd | 156 | 0.28332235 | 0.00033843 | 0.0126731 | 1 |
| 232 | Blcap | 155 | 0.28412878 | 0.00033984 | 0.0127072 | 1 |
| 1692 | Wls | 158 | -0.281381 | 0.00034168 | 0.01275536 | 1 |
| 285 | Ccer1 | 157 | -0.2822271 | 0.00034213 | 0.01275536 | 1 |
| 637 | Gemin4 | 154 | 0.28470591 | 0.00034534 | 0.01284348 | 1 |
| 1465 | Snrnp200 | 159 | -0.2802988 | 0.0003455 | 0.01284348 | 1 |
| 1227 | Pparg | 160 | -0.2793979 | 0.0003463 | 0.01285453 | 1 |
| 804 | Itm2b | 158 | -0.2810397 | 0.00034765 | 0.01288587 | 1 |
| 209 | Axin2 | 160 | -0.2792711 | 0.00034855 | 0.01290033 | 1 |
| 1600 | Tmsb15l | 154 | -0.2844082 | 0.00035054 | 0.01293879 | 1 |
| 1105 | Nwd2 | 159 | -0.2800104 | 0.0003506 | 0.01293879 | 1 |
| 476 | Dpm3 | 157 | 0.28169901 | 0.00035139 | 0.01294912 | 1 |
| 185 | Ascc1 | 158 | -0.2806862 | 0.00035394 | 0.01302408 | 1 |
| 1393 | Sh3gl2 | 159 | -0.2794487 | 0.00036075 | 0.01325562 | 1 |
| 1146 | P3h3 | 159 | 0.27937694 | 0.00036207 | 0.01326927 | 1 |
| 1193 | Phlda3 | 155 | -0.2828449 | 0.00036249 | 0.01326927 | 1 |
| 1470 | Snx31 | 158 | -0.2802038 | 0.00036269 | 0.01326927 | 1 |
| 4 | 1700026L06Rik | 159 | -0.2792643 | 0.00036414 | 0.01329597 | 1 |
| 770 | Ifit1 | 155 | 0.28273685 | 0.00036446 | 0.01329597 | 1 |
| 717 | Haus4 | 157 | 0.28094518 | 0.00036502 | 0.01329738 | 1 |
| 1189 | Pgr | 158 | 0.28002141 | 0.00036605 | 0.01331569 | 1 |
| 573 | Fasn | 159 | -0.2790909 | 0.00036735 | 0.01332598 | 1 |
| 1337 | Rpl13 | 160 | -0.2782377 | 0.00036737 | 0.01332598 | 1 |
| 208 | Axin2 | 160 | -0.2781749 | 0.00036855 | 0.01334958 | 1 |
| 772 | Ifne | 156 | -0.2815556 | 0.00036993 | 0.0133805 | 1 |
| 51 | 6820408C15Rik | 157 | -0.2805318 | 0.0003727 | 0.01346156 | 1 |
| 1444 | Slc7a11 | 153 | -0.284051 | 0.00037329 | 0.01346364 | 1 |
| 272 | Ccdc106 | 151 | 0.28584818 | 0.00037383 | 0.01346405 | 1 |
| 672 | Gorasp2 | 157 | -0.280317 | 0.00037675 | 0.01354523 | 1 |
| 1555 | Tctex1d1 | 159 | -0.2785713 | 0.00037714 | 0.01354523 | 1 |
| 679 | Gpr137b | 158 | 0.27936873 | 0.00037831 | 0.01356784 | 1 |
| 1629 | Ttll8 | 159 | -0.27843 | 0.00037985 | 0.01360348 | 1 |
| 1641 | Ube2q2 | 155 | 0.28188314 | 0.00038037 | 0.01360348 | 1 |
| 1154 | Panx3 | 156 | 0.28084963 | 0.00038326 | 0.01368771 | 1 |
| 1654 | Usp16 | 160 | -0.2773045 | 0.00038518 | 0.01370428 | 1 |
| 1617 | Trim8 | 156 | -0.280739 | 0.00038539 | 0.01370428 | 1 |
| 135 | Ankrd54 | 160 | -0.277285 | 0.00038556 | 0.01370428 | 1 |

| | | | | | | |
|---|---|---|---|---|---|---|
| 53 | 8430408G22Rik | 156 | 0.28068676 | 0.0003864 | 0.01370428 | 1 |
| 1662 | Uty | 151 | -0.2851775 | 0.00038642 | 0.01370428 | 1 |
| 109 | AI987944 | 159 | -0.2780187 | 0.00038782 | 0.0137154 | 1 |
| 740 | Hist1h4i | 158 | -0.2788647 | 0.00038803 | 0.0137154 | 1 |
| 231 | Birc5 | 153 | 0.28324494 | 0.00038853 | 0.0137154 | 1 |
| 1297 | Rdh14 | 160 | -0.277102 | 0.00038915 | 0.0137154 | 1 |
| 687 | Gpr83 | 156 | 0.28053097 | 0.00038942 | 0.0137154 | 1 |
| 869 | Letmd1 | 158 | 0.27872666 | 0.00039074 | 0.01374267 | 1 |
| 1369 | Scml1 | 158 | -0.2785904 | 0.00039342 | 0.01380471 | 1 |
| 1469 | Snx27 | 159 | -0.2777069 | 0.00039397 | 0.01380471 | 1 |
| 1520 | Syn3 | 160 | -0.2768507 | 0.00039413 | 0.01380471 | 1 |
| 1204 | Pla2g7 | 146 | -0.2892313 | 0.000399 | 0.0139564 | 1 |
| 196 | Atn1 | 158 | -0.2782613 | 0.00039998 | 0.01395666 | 1 |
| 960 | Mbd2 | 156 | -0.2799723 | 0.00040044 | 0.01395666 | 1 |
| 1636 | Tyms | 158 | 0.27822765 | 0.00040066 | 0.01395666 | 1 |
| 944 | Magee1 | 159 | -0.2773375 | 0.00040136 | 0.01396222 | 1 |
| 1750 | Zmiz1 | 158 | -0.2781319 | 0.00040259 | 0.01397631 | 1 |
| 115 | Akna | 158 | -0.2780887 | 0.00040346 | 0.01397631 | 1 |
| 933 | Ly6g5b | 160 | -0.2763714 | 0.00040378 | 0.01397631 | 1 |
| 738 | Hist1h2bc | 160 | -0.2763624 | 0.00040396 | 0.01397631 | 1 |
| 384 | Col4a5 | 160 | -0.2762942 | 0.00040536 | 0.01399097 | 1 |
| 730 | Hectd2 | 160 | -0.2762878 | 0.00040549 | 0.01399097 | 1 |
| 581 | Fbxo2 | 160 | -0.2761871 | 0.00040755 | 0.0140432 | 1 |
| 314 | Cdhr4 | 151 | -0.2840433 | 0.0004086 | 0.01405639 | 1 |
| 563 | Fam25c | 156 | -0.2795466 | 0.00040904 | 0.01405639 | 1 |
| 727 | Hc | 158 | -0.2777657 | 0.00041005 | 0.01406567 | 1 |
| 6 | 1700052N19Rik | 155 | 0.28032284 | 0.00041111 | 0.01406567 | 1 |
| 986 | Mknk1 | 159 | 0.27685793 | 0.00041116 | 0.01406567 | 1 |
| 85 | Acot2 | 158 | -0.2776942 | 0.00041152 | 0.01406567 | 1 |
| 288 | Ccnd3 | 155 | -0.2801454 | 0.00041475 | 0.01414783 | 1 |
| 575 | Fau | 160 | -0.2758125 | 0.00041532 | 0.01414783 | 1 |
| 39 | 4930548F15Rik | 159 | -0.2766446 | 0.00041559 | 0.01414783 | 1 |
| 1545 | Tbrg4 | 155 | 0.28003808 | 0.00041697 | 0.01417579 | 1 |
| 875 | Lhx8 | 160 | 0.2757028 | 0.00041762 | 0.0141789 | 1 |
| 1152 | Palm2 | 156 | -0.2788868 | 0.00042269 | 0.01433222 | 1 |
| 1659 | Utp14b | 160 | -0.2753528 | 0.00042503 | 0.01438559 | 1 |
| 956 | Masp2 | 156 | 0.27875853 | 0.0004254 | 0.01438559 | 1 |
| 207 | Avp | 160 | 0.27523461 | 0.00042756 | 0.01442469 | 1 |
| 440 | Dctn1 | 156 | -0.2786216 | 0.0004283 | 0.01442469 | 1 |
| 1657 | Usp31 | 158 | -0.2768943 | 0.00042832 | 0.01442469 | 1 |
| 139 | Anxa2 | 160 | -0.2751684 | 0.00042899 | 0.01442469 | 1 |
| 1495 | Sry | 156 | -0.2785706 | 0.00042939 | 0.01442469 | 1 |
| 1741 | Zfp790 | 160 | -0.2751118 | 0.00043021 | 0.01443328 | 1 |
| 166 | Arhgap22 | 157 | 0.27754451 | 0.0004328 | 0.01450106 | 1 |
| 1213 | Plp2 | 158 | 0.27657797 | 0.00043514 | 0.0145484 | 1 |

| | | | | | | |
|---|---|---|---|---|---|---|
| 433 | Dalrd3 | 156 | -0.2782925 | 0.00043536 | 0.0145484 | 1 |
| 1302 | Rfx1 | 160 | -0.2748253 | 0.00043644 | 0.01456552 | 1 |
| 1673 | Vps11 | 155 | 0.2790748 | 0.00043734 | 0.0145733 | 1 |
| 140 | Anxa2 | 157 | -0.2772701 | 0.00043875 | 0.0145733 | 1 |
| 280 | Ccdc58 | 159 | -0.2755217 | 0.00043964 | 0.0145733 | 1 |
| 1694 | Wnt1 | 160 | -0.2746775 | 0.00043969 | 0.0145733 | 1 |
| 389 | Commd7 | 160 | -0.2746663 | 0.00043993 | 0.0145733 | 1 |
| 330 | Cep112 | 160 | -0.2746584 | 0.00044011 | 0.0145733 | 1 |
| 1044 | Nap1l3 | 159 | -0.2754425 | 0.00044138 | 0.01459025 | 1 |
| 239 | Bpifb1 | 160 | 0.27456863 | 0.00044209 | 0.01459025 | 1 |
| 230 | Bik | 158 | 0.27624874 | 0.00044234 | 0.01459025 | 1 |
| 971 | Meis1 | 136 | -0.2970658 | 0.00044474 | 0.01465069 | 1 |
| 1686 | Wdr63 | 158 | -0.2760351 | 0.00044707 | 0.01470819 | 1 |
| 395 | Cox7b | 160 | -0.2742569 | 0.00044905 | 0.01475422 | 1 |
| 1118 | Olfm1 | 160 | -0.2742153 | 0.00044998 | 0.01476038 | 1 |
| 1422 | Slc25a41 | 158 | -0.2758611 | 0.00045096 | 0.01476038 | 1 |
| 636 | Gclc | 155 | 0.27845346 | 0.00045097 | 0.01476038 | 1 |
| 668 | Gnao1 | 160 | -0.2740196 | 0.00045441 | 0.01485379 | 1 |
| 456 | Dido1 | 156 | -0.2773751 | 0.00045559 | 0.01487319 | 1 |
| 699 | Grm8 | 160 | -0.2738695 | 0.00045783 | 0.01492733 | 1 |
| 683 | Gpr165 | 159 | 0.2746577 | 0.00045901 | 0.0149466 | 1 |
| 591 | Fgd6 | 155 | 0.27802349 | 0.00046063 | 0.01498026 | 1 |
| 929 | Lsm4 | 155 | 0.27775033 | 0.00046687 | 0.01516368 | 1 |
| 492 | Ear6 | 159 | -0.2742751 | 0.00046784 | 0.01517581 | 1 |
| 624 | Gal3st2 | 160 | -0.2732903 | 0.00047126 | 0.01526749 | 1 |
| 828 | Kctd8 | 160 | -0.2732365 | 0.00047253 | 0.01527508 | 1 |
| 1462 | Snapc3 | 159 | -0.2740674 | 0.0004727 | 0.01527508 | 1 |
| 887 | LOC432456 | 160 | -0.2731815 | 0.00047383 | 0.01529022 | 1 |
| 1017 | Mthfd2 | 159 | 0.27399643 | 0.00047437 | 0.01529022 | 1 |
| 739 | Hist1h2bc | 146 | 0.2854659 | 0.00047832 | 0.01539806 | 1 |
| 111 | Aip | 156 | -0.2763617 | 0.00047895 | 0.01539886 | 1 |
| 1184 | Pex13 | 160 | -0.2729048 | 0.0004804 | 0.01542623 | 1 |
| 814 | Kcmf1 | 159 | -0.2736679 | 0.00048217 | 0.01546351 | 1 |
| 77 | Acads | 155 | 0.27692783 | 0.00048612 | 0.01557042 | 1 |
| 293 | Cct2 | 160 | -0.2725531 | 0.00048889 | 0.01563943 | 1 |
| 23 | 2610020H08Rik | 154 | 0.27758054 | 0.00049146 | 0.01570204 | 1 |
| 906 | LOC433740 | 155 | -0.276625 | 0.00049339 | 0.01572481 | 1 |
| 1125 | Olfr561 | 158 | -0.2740421 | 0.00049352 | 0.01572481 | 1 |
| 1436 | Slc48a1 | 159 | -0.2731782 | 0.00049402 | 0.01572481 | 1 |
| 1480 | Spa17 | 160 | -0.2722721 | 0.00049576 | 0.01576038 | 1 |
| 1590 | Tmem203 | 130 | 0.30116148 | 0.00049805 | 0.01581342 | 1 |
| 147 | Aph1b | 154 | -0.2770749 | 0.00050374 | 0.01597423 | 1 |
| 1676 | Vps51 | 159 | -0.2727117 | 0.00050557 | 0.01601212 | 1 |
| 1192 | Phgdh | 156 | -0.2750806 | 0.00051006 | 0.01613425 | 1 |
| 1267 | Ptpdc1 | 159 | -0.2724017 | 0.00051337 | 0.01621904 | 1 |

| | | | | | | |
|---|---|---|---|---|---|---|
| 1473 | Soga3 | 160 | -0.2715136 | 0.00051477 | 0.01624299 | 1 |
| 850 | Krt32 | 159 | -0.2722993 | 0.00051598 | 0.0162609 | 1 |
| 535 | Eqtn | 160 | -0.2714297 | 0.00051691 | 0.01627031 | 1 |
| 1211 | Plekhm2 | 157 | -0.2739156 | 0.00051783 | 0.01627895 | 1 |
| 1744 | Zfp952 | 156 | -0.2746918 | 0.00051986 | 0.01630356 | 1 |
| 1288 | Rbl2 | 159 | -0.2721146 | 0.0005207 | 0.01630356 | 1 |
| 1490 | Sptb | 159 | -0.2721049 | 0.00052095 | 0.01630356 | 1 |
| 467 | Dnah3 | 158 | -0.2729365 | 0.00052117 | 0.01630356 | 1 |
| 1157 | Pax3 | 158 | -0.2728116 | 0.00052438 | 0.016381 | 1 |
| 485 | Dus2 | 159 | -0.2719506 | 0.00052493 | 0.016381 | 1 |
| 28 | 2900092C05Rik | 160 | -0.2710672 | 0.00052627 | 0.01640254 | 1 |
| 684 | Gpr165 | 160 | 0.27102891 | 0.00052726 | 0.01641353 | 1 |
| 1517 | Svs3a | 159 | 0.27172641 | 0.00053077 | 0.01648384 | 1 |
| 510 | Eif5 | 160 | -0.270893 | 0.00053082 | 0.01648384 | 1 |
| 1030 | Mx1 | 155 | 0.27497982 | 0.00053467 | 0.01658313 | 1 |
| 1331 | Rnf138rt1 | 147 | 0.28188299 | 0.00054241 | 0.0167972 | 1 |
| 1295 | Rcc1 | 154 | 0.27553598 | 0.00054289 | 0.0167972 | 1 |
| 1163 | Pcsk1n | 159 | 0.27115442 | 0.00054592 | 0.01687058 | 1 |
| 604 | Frmd5 | 159 | -0.2711168 | 0.00054693 | 0.01688131 | 1 |
| 1542 | Tbc1d9 | 157 | -0.2726799 | 0.00055013 | 0.01693362 | 1 |
| 381 | Cog2 | 155 | 0.27437541 | 0.00055062 | 0.01693362 | 1 |
| 745 | Hnrnpk | 138 | -0.2903737 | 0.00055062 | 0.01693362 | 1 |
| 189 | Atf2 | 148 | -0.2805731 | 0.00055229 | 0.01696441 | 1 |
| 141 | Aox1 | 152 | 0.27687699 | 0.00055408 | 0.01699884 | 1 |
| 188 | Aspn | 155 | 0.27411276 | 0.00055768 | 0.01706181 | 1 |
| 1170 | Pdgfrb | 160 | -0.2698757 | 0.00055813 | 0.01706181 | 1 |
| 757 | Hsp90aa1 | 147 | -0.2812805 | 0.00055814 | 0.01706181 | 1 |
| 1702 | Xpo1 | 136 | 0.29204126 | 0.00056091 | 0.01712595 | 1 |
| 1164 | Pdcd2 | 158 | -0.2713781 | 0.00056258 | 0.0171498 | 1 |
| 1621 | Tspan15 | 160 | -0.2696977 | 0.00056304 | 0.0171498 | 1 |
| 62 | A830029E22Rik | 159 | -0.2704267 | 0.00056578 | 0.01721274 | 1 |
| 628 | Galnt13 | 160 | -0.2695325 | 0.00056763 | 0.01724849 | 1 |
| 1643 | Ublcp1 | 156 | 0.27284804 | 0.00056877 | 0.01726237 | 1 |
| 1370 | Scp2 | 153 | 0.27541613 | 0.00056967 | 0.01726513 | 1 |
| 1748 | Zg16 | 156 | 0.27278709 | 0.00057046 | 0.01726513 | 1 |
| 205 | Atp6v1c2 | 155 | 0.27362988 | 0.00057089 | 0.01726513 | 1 |
| 987 | Mllt1 | 137 | -0.2904941 | 0.00057411 | 0.0173418 | 1 |
| 811 | Jrkl | 158 | -0.270937 | 0.00057484 | 0.0173433 | 1 |
| 262 | Capza1 | 155 | 0.27341197 | 0.00057695 | 0.01738631 | 1 |
| 248 | C2cd5 | 159 | -0.2698839 | 0.00058103 | 0.01747316 | 1 |
| 1071 | Ngfr | 160 | 0.26905146 | 0.0005812 | 0.01747316 | 1 |
| 1277 | Rab39 | 156 | 0.27217089 | 0.00058777 | 0.01764977 | 1 |
| 977 | Mfsd6l | 153 | 0.27464781 | 0.00059112 | 0.01771576 | 1 |
| 992 | Mmp7 | 148 | 0.2791168 | 0.0005918 | 0.01771576 | 1 |
| 910 | LOC434271 | 159 | -0.2694993 | 0.00059206 | 0.01771576 | 1 |

| | | | | | | |
|---|---|---|---|---|---|---|
| 1018 | Mthfsd | 146 | -0.2808921 | 0.00059416 | 0.01775776 | 1 |
| 1751 | Zpld1 | 158 | -0.2702228 | 0.00059522 | 0.01776855 | 1 |
| 504 | Eif2ak3 | 160 | -0.2683493 | 0.00060155 | 0.01793659 | 1 |
| 998 | Morc3 | 160 | -0.2682505 | 0.00060446 | 0.0180024 | 1 |
| 1352 | Rtf1 | 158 | -0.2697206 | 0.00060994 | 0.0181442 | 1 |
| 89 | Acta2 | 158 | -0.2696845 | 0.00061101 | 0.01815491 | 1 |
| 1498 | St8sia4 | 150 | 0.27658738 | 0.00061199 | 0.01816298 | 1 |
| 1453 | Smim11 | 158 | 0.26959837 | 0.00061357 | 0.01817615 | 1 |
| 1530 | Sytl5 | 160 | 0.26793481 | 0.00061387 | 0.01817615 | 1 |
| 1441 | Slc6a17 | 157 | -0.2703779 | 0.00061533 | 0.0181982 | 1 |
| 1327 | Rnasel | 159 | 0.26853139 | 0.00062067 | 0.01830528 | 1 |
| 371 | Cmtr1 | 153 | 0.27362069 | 0.00062095 | 0.01830528 | 1 |
| 1371 | Sdr16c5 | 156 | -0.2710304 | 0.0006211 | 0.01830528 | 1 |
| 1733 | Zfp384 | 158 | -0.2691402 | 0.00062737 | 0.0184686 | 1 |
| 1364 | Sbno2 | 151 | 0.27508356 | 0.00062993 | 0.0185227 | 1 |
| 789 | Iqsec1 | 160 | -0.2673219 | 0.00063251 | 0.01856237 | 1 |
| 296 | Cd247 | 154 | 0.27235844 | 0.00063274 | 0.01856237 | 1 |
| 228 | Bet1 | 152 | 0.27398724 | 0.00063628 | 0.01863881 | 1 |
| 1045 | Napb | 156 | -0.2705124 | 0.00063681 | 0.01863881 | 1 |
| 565 | Fam58b | 138 | 0.28716147 | 0.00063802 | 0.01865267 | 1 |
| 1411 | Slc16a5 | 156 | -0.270398 | 0.00064033 | 0.0186931 | 1 |
| 702 | Gsdmcl-ps | 158 | -0.2686829 | 0.00064143 | 0.0186931 | 1 |
| 1726 | Zdhhc9 | 160 | -0.2670288 | 0.0006416 | 0.0186931 | 1 |
| 96 | Adat1 | 156 | 0.27030035 | 0.00064335 | 0.01872247 | 1 |
| 777 | Ighg2a | 159 | -0.2676811 | 0.00064686 | 0.01880319 | 1 |
| 708 | Gucd1 | 147 | 0.27810144 | 0.00064832 | 0.01882002 | 1 |
| 167 | Arhgap29 | 160 | -0.2667959 | 0.00064892 | 0.01882002 | 1 |
| 1165 | Pdcd6ip | 159 | -0.2674909 | 0.00065286 | 0.01891275 | 1 |
| 294 | Cct2 | 160 | -0.2665972 | 0.00065522 | 0.01894411 | 1 |
| 1574 | Tlr6 | 158 | 0.26823665 | 0.00065543 | 0.01894411 | 1 |
| 1087 | Nptx2 | 159 | -0.2673352 | 0.0006578 | 0.01899121 | 1 |
| 1607 | Tor4a | 155 | 0.27063278 | 0.00065955 | 0.01902027 | 1 |
| 579 | Fbrs | 160 | -0.2664146 | 0.00066105 | 0.01904194 | 1 |
| 649 | Gm10549 | 160 | 0.26637012 | 0.00066248 | 0.0190616 | 1 |
| 52 | 6820408C15Rik | 160 | -0.2662862 | 0.00066519 | 0.01910179 | 1 |
| 936 | Lyz2 | 158 | -0.2679245 | 0.00066538 | 0.01910179 | 1 |
| 947 | Maoa | 160 | -0.2662374 | 0.00066677 | 0.01912005 | 1 |
| 419 | Cubn | 160 | -0.2662062 | 0.00066778 | 0.01912743 | 1 |
| 877 | Lig3 | 158 | -0.2677399 | 0.00067134 | 0.01917292 | 1 |
| 338 | Cfp | 157 | -0.2685477 | 0.00067215 | 0.01917292 | 1 |
| 980 | Mia | 159 | -0.2668786 | 0.00067251 | 0.01917292 | 1 |
| 546 | Eya1 | 160 | -0.2660261 | 0.00067364 | 0.01917292 | 1 |
| 391 | Copz2 | 156 | -0.2693417 | 0.00067367 | 0.01917292 | 1 |
| 1385 | Sf3b6 | 126 | 0.29891822 | 0.00067388 | 0.01917292 | 1 |
| 373 | Cngb1 | 155 | 0.27009731 | 0.00067667 | 0.01921167 | 1 |

| | | | | | | |
|---|---|---|---|---|---|---|
| 1557 | Tdrd3 | 156 | 0.26924634 | 0.00067675 | 0.01921167 | 1 |
| 1435 | Slc41a3 | 160 | -0.2659055 | 0.00067759 | 0.01921398 | 1 |
| 1738 | Zfp748 | 155 | 0.26994203 | 0.00068171 | 0.0193093 | 1 |
| 816 | Kcna5 | 158 | 0.26730972 | 0.0006854 | 0.01939234 | 1 |
| 661 | Gm520 | 157 | 0.26804049 | 0.00068874 | 0.01946508 | 1 |
| 1389 | Sfxn4 | 159 | -0.2662508 | 0.00069321 | 0.01955122 | 1 |
| 531 | Epha1 | 158 | -0.2670492 | 0.00069405 | 0.01955122 | 1 |
| 34 | 4921531P07Rik | 156 | -0.2687181 | 0.00069409 | 0.01955122 | 1 |
| 1293 | Rc3h1 | 159 | -0.2661264 | 0.00069739 | 0.01962237 | 1 |
| 867 | Lenep | 158 | 0.26688694 | 0.00069949 | 0.01963062 | 1 |
| 305 | Cdc5l | 159 | -0.2660477 | 0.00070004 | 0.01963062 | 1 |
| 1427 | Slc2a1 | 160 | -0.2652248 | 0.00070029 | 0.01963062 | 1 |
| 1210 | Plekhj1 | 158 | 0.26684915 | 0.00070076 | 0.01963062 | 1 |
| 375 | Cnnm2 | 160 | -0.2651236 | 0.00070373 | 0.019692 | 1 |
| 592 | Fgf12 | 160 | -0.2650666 | 0.00070567 | 0.0197247 | 1 |
| 211 | B3gnt9 | 158 | -0.2666052 | 0.00070903 | 0.01978307 | 1 |
| 901 | LOC433588 | 159 | -0.2657746 | 0.00070931 | 0.01978307 | 1 |
| 128 | Alox12e | 157 | 0.26736162 | 0.00071152 | 0.01982303 | 1 |
| 1421 | Slc25a32 | 155 | 0.26895217 | 0.00071465 | 0.01988399 | 1 |
| 574 | Fastkd5 | 155 | 0.26890242 | 0.00071635 | 0.01988399 | 1 |
| 300 | Cd96 | 156 | 0.26802975 | 0.0007173 | 0.01988399 | 1 |
| 1510 | Suco | 160 | -0.2647282 | 0.0007173 | 0.01988399 | 1 |
| 1727 | Zfand2a | 152 | 0.27144844 | 0.00071762 | 0.01988399 | 1 |
| 1110 | Oas1b | 154 | 0.26960584 | 0.00072145 | 0.0199684 | 1 |
| 1350 | Rsad2 | 154 | -0.2695727 | 0.00072258 | 0.01996867 | 1 |
| 168 | Arhgap4 | 156 | 0.2678473 | 0.00072357 | 0.01996867 | 1 |
| 1226 | Postn | 156 | -0.2678353 | 0.00072398 | 0.01996867 | 1 |
| 1232 | Ppm1a | 156 | -0.2678175 | 0.00072459 | 0.01996867 | 1 |
| 1006 | Mrgprb4 | 160 | -0.2644947 | 0.00072542 | 0.01996994 | 1 |
| 555 | Fam169b | 155 | 0.26856385 | 0.00072797 | 0.02001691 | 1 |
| 80 | Accs | 158 | 0.2660347 | 0.0007287 | 0.02001691 | 1 |
| 49 | 6030419C18Rik | 156 | -0.2676422 | 0.00073068 | 0.02004947 | 1 |
| 57 | 9330109E03Rik | 158 | -0.2658351 | 0.0007357 | 0.02015686 | 1 |
| 2 | 1600029I14Rik | 159 | -0.2649921 | 0.00073652 | 0.02015686 | 1 |
| 1123 | Olfr424 | 158 | -0.2657916 | 0.00073724 | 0.02015686 | 1 |
| 1658 | Usp54 | 159 | -0.2649377 | 0.00073844 | 0.02015686 | 1 |
| 928 | Lsm1 | 156 | -0.2674172 | 0.00073855 | 0.02015686 | 1 |
| 836 | Kifc2 | 160 | -0.2640379 | 0.00074156 | 0.02018885 | 1 |
| 1431 | Slc38a1 | 158 | 0.26566576 | 0.00074169 | 0.02018885 | 1 |
| 173 | Arl10 | 160 | 0.26401935 | 0.00074223 | 0.02018885 | 1 |
| 908 | LOC433773 | 158 | -0.2656321 | 0.00074289 | 0.02018885 | 1 |
| 542 | Etfdh | 159 | 0.26477451 | 0.00074425 | 0.02020427 | 1 |
| 690 | Gpx6 | 158 | 0.26555222 | 0.00074573 | 0.02022295 | 1 |
| 881 | LOC195357 | 158 | 0.26552293 | 0.00074678 | 0.02022976 | 1 |
| 1134 | Orc5 | 158 | 0.26539781 | 0.00075126 | 0.02030987 | 1 |

| | | | | | | |
|---|---|---|---|---|---|---|
| 1212 | Pllp | 160 | -0.2637659 | 0.00075133 | 0.02030987 | 1 |
| 82 | Acer2 | 158 | -0.2652332 | 0.0007572 | 0.0204467 | 1 |
| 180 | Arrdc3 | 158 | -0.2651652 | 0.00075966 | 0.02049145 | 1 |
| 1534 | Tal1 | 160 | -0.2634747 | 0.00076192 | 0.02053072 | 1 |
| 808 | Jakmip3 | 158 | -0.265017 | 0.00076505 | 0.02057624 | 1 |
| 445 | Ddx52 | 159 | 0.26419465 | 0.00076523 | 0.02057624 | 1 |
| 966 | Mdm1 | 156 | -0.2666328 | 0.0007666 | 0.02059145 | 1 |
| 243 | Btg3 | 156 | 0.26651093 | 0.00077105 | 0.02068904 | 1 |
| 1503 | Stau1 | 152 | 0.26972197 | 0.00077828 | 0.02086112 | 1 |
| 559 | Fam20a | 160 | -0.2629238 | 0.00078233 | 0.02094762 | 1 |
| 1224 | Pomc | 160 | 0.26281756 | 0.00078632 | 0.02103154 | 1 |
| 1047 | Narfl | 156 | 0.26607577 | 0.00078711 | 0.02103154 | 1 |
| 256 | Cadm3 | 160 | -0.2625691 | 0.00079573 | 0.02123951 | 1 |
| 488 | E230008N13Rik | 158 | -0.2640204 | 0.00080226 | 0.0213682 | 1 |
| 429 | Cypt1 | 159 | -0.2631903 | 0.00080285 | 0.0213682 | 1 |
| 83 | Acer3 | 160 | 0.26237717 | 0.00080307 | 0.0213682 | 1 |
| 1645 | Ubxn10 | 158 | -0.2639303 | 0.00080571 | 0.02141594 | 1 |
| 404 | Cradd | 159 | -0.2630723 | 0.00080738 | 0.02143799 | 1 |
| 163 | Arf6 | 155 | 0.26630458 | 0.0008101 | 0.02148491 | 1 |
| 1197 | Pik3cb | 157 | -0.2646186 | 0.00081083 | 0.02148491 | 1 |
| 1042 | Nanp | 156 | 0.26541272 | 0.00081219 | 0.02148644 | 1 |
| 1569 | Ticam2 | 158 | -0.2637516 | 0.00081258 | 0.02148644 | 1 |
| 879 | Lmbr1l | 156 | 0.26535181 | 0.00081453 | 0.02151572 | 1 |
| 1546 | Tbx2 | 158 | 0.26344447 | 0.00082451 | 0.02175687 | 1 |
| 756 | Hsf2 | 152 | 0.26837862 | 0.0008287 | 0.0218447 | 1 |
| 317 | Cdk5rap2 | 159 | -0.2625004 | 0.00082966 | 0.02184746 | 1 |
| 237 | Bpgm | 160 | -0.2616626 | 0.00083095 | 0.02185877 | 1 |
| 165 | Arhgap18 | 158 | 0.26312977 | 0.00083691 | 0.02199285 | 1 |
| 1244 | Prdm2 | 136 | -0.2831158 | 0.0008384 | 0.02200922 | 1 |
| 93 | Adamts18 | 152 | 0.2680596 | 0.00084111 | 0.02204388 | 1 |
| 1497 | St3gal6 | 160 | -0.2613992 | 0.00084145 | 0.02204388 | 1 |
| 1049 | Natd1 | 148 | 0.27147038 | 0.00084542 | 0.02212515 | 1 |
| 1387 | Sft2d2 | 157 | 0.26364433 | 0.00084907 | 0.02219773 | 1 |
| 825 | Kcnq1ot1 | 159 | -0.2619444 | 0.00085187 | 0.02220363 | 1 |
| 577 | Fbp1 | 154 | 0.2660745 | 0.00085199 | 0.02220363 | 1 |
| 1706 | Yipf5 | 158 | 0.26273831 | 0.00085257 | 0.02220363 | 1 |
| 1294 | Rcbtb2 | 152 | 0.26776337 | 0.00085278 | 0.02220363 | 1 |
| 460 | Dlx2 | 154 | 0.26581372 | 0.00086244 | 0.02243226 | 1 |
| 1687 | Wdr77 | 159 | -0.2615269 | 0.00086891 | 0.02255743 | 1 |
| 411 | Csdc2 | 158 | -0.2623341 | 0.00086902 | 0.02255743 | 1 |
| 406 | Creb3l3 | 155 | 0.26477625 | 0.0008704 | 0.02257004 | 1 |
| 1685 | Wdpcp | 157 | 0.26308637 | 0.0008717 | 0.02258084 | 1 |
| 222 | Bche | 160 | -0.2606091 | 0.00087368 | 0.02258708 | 1 |
| 1074 | Nhsl1 | 160 | -0.2606082 | 0.00087371 | 0.02258708 | 1 |
| 732 | Herpud1 | 158 | -0.2621531 | 0.00087649 | 0.02261645 | 1 |

| | | | | | | |
|---|---|---|---|---|---|---|
| 361 | Clec4a3 | 159 | -0.2613125 | 0.00087778 | 0.02261645 | 1 |
| 17 | 2310035C23Rik | 158 | -0.2621102 | 0.00087826 | 0.02261645 | 1 |
| 320 | Cds1 | 158 | -0.2621068 | 0.0008784 | 0.02261645 | 1 |
| 352 | Cidea | 160 | -0.2604101 | 0.00088197 | 0.02268536 | 1 |
| 1731 | Zfp353-ps | 158 | -0.2619847 | 0.00088348 | 0.02269142 | 1 |
| 1361 | Samd14 | 160 | -0.260362 | 0.00088399 | 0.02269142 | 1 |
| 41 | 4931408C20Rik | 160 | -0.2603102 | 0.00088616 | 0.02272436 | 1 |
| 1377 | Serpina1a | 152 | 0.26685991 | 0.0008893 | 0.02276963 | 1 |
| 590 | Fezf1 | 160 | 0.2602258 | 0.00088972 | 0.02276963 | 1 |
| 1075 | Ninj1 | 160 | -0.2601335 | 0.00089362 | 0.0228352 | 1 |
| 658 | Gm471 | 158 | -0.2617319 | 0.00089407 | 0.0228352 | 1 |
| 909 | LOC433775 | 159 | -0.2607116 | 0.00090308 | 0.02304173 | 1 |
| 926 | Lrrc40 | 153 | 0.26559803 | 0.00090624 | 0.02304173 | 1 |
| 407 | Crh | 157 | 0.26225031 | 0.00090665 | 0.02304173 | 1 |
| 274 | Ccdc109b | 160 | -0.2598155 | 0.00090719 | 0.02304173 | 1 |
| 952 | Mapk8 | 153 | -0.2655701 | 0.00090741 | 0.02304173 | 1 |
| 1651 | Unc80 | 159 | -0.2606061 | 0.00090759 | 0.02304173 | 1 |
| 1115 | Obox6 | 155 | 0.26383832 | 0.00090941 | 0.02306499 | 1 |
| 1585 | Tmem144 | 152 | 0.26630982 | 0.00091224 | 0.02311373 | 1 |
| 1540 | Tasp1 | 152 | 0.26618573 | 0.00091749 | 0.0232236 | 1 |
| 267 | Cbln2 | 146 | -0.2714452 | 0.00091924 | 0.02324486 | 1 |
| 883 | LOC240906 | 158 | -0.2611132 | 0.0009205 | 0.02325055 | 1 |
| 1322 | Ripk3 | 156 | 0.26273059 | 0.00092129 | 0.02325055 | 1 |
| 1284 | Ranbp2 | 157 | -0.261856 | 0.00092358 | 0.02328523 | 1 |
| 1259 | Psmc4 | 149 | -0.2686054 | 0.00092559 | 0.0233127 | 1 |
| 1538 | Tars | 158 | 0.26092915 | 0.0009285 | 0.02336301 | 1 |
| 964 | Mccc2 | 155 | 0.2631937 | 0.00093715 | 0.02353608 | 1 |
| 856 | Lamp1 | 157 | -0.2615334 | 0.00093765 | 0.02353608 | 1 |
| 621 | Gak | 157 | -0.2615088 | 0.00093873 | 0.02353608 | 1 |
| 1418 | Slc22a8 | 160 | -0.2590849 | 0.00093908 | 0.02353608 | 1 |
| 1401 | Sin3b | 159 | -0.2597811 | 0.00094358 | 0.02362572 | 1 |
| 623 | Gal | 160 | 0.25891193 | 0.00094677 | 0.0236757 | 1 |
| 344 | Chia1 | 152 | 0.26547517 | 0.00094808 | 0.0236757 | 1 |
| 3 | 1700019L03Rik | 159 | -0.2596736 | 0.00094837 | 0.0236757 | 1 |
| 611 | G3bp1 | 156 | 0.26205335 | 0.00095089 | 0.02371546 | 1 |
| 149 | Apoa1bp | 159 | -0.259557 | 0.00095358 | 0.02375933 | 1 |
| 536 | Erlin1 | 155 | 0.26274029 | 0.00095712 | 0.02381949 | 1 |
| 554 | Fam151b | 160 | -0.2586649 | 0.00095787 | 0.02381949 | 1 |
| 304 | Cdc42ep2 | 155 | 0.26264558 | 0.00096134 | 0.02388256 | 1 |
| 805 | Itpk1 | 160 | -0.2584551 | 0.00096739 | 0.02400927 | 1 |
| 1079 | Nkx1-1 | 156 | -0.2616624 | 0.00096837 | 0.02401038 | 1 |
| 1238 | Ppp2r3d | 160 | -0.2583457 | 0.00097238 | 0.02408634 | 1 |
| 676 | Gpr101 | 160 | 0.25829994 | 0.00097448 | 0.02411482 | 1 |
| 1057 | Ndufs1 | 160 | -0.2581608 | 0.00098088 | 0.02424968 | 1 |
| 1177 | Pef1 | 144 | -0.2717494 | 0.00098491 | 0.02429596 | 1 |

| | | | | | | |
|---|---|---|---|---|---|---|
| 1587 | Tmem160 | 159 | -0.2588376 | 0.00098635 | 0.02429596 | 1 |
| 665 | Gm816 | 152 | -0.2646118 | 0.00098651 | 0.02429596 | 1 |
| 831 | Khk | 154 | 0.26292034 | 0.00098657 | 0.02429596 | 1 |
| 650 | Gm1140 | 156 | -0.2611461 | 0.00099191 | 0.02440221 | 1 |
| 1483 | Spata18 | 159 | -0.2586712 | 0.00099407 | 0.02440221 | 1 |
| 1722 | Zcchc7 | 159 | -0.2586585 | 0.00099466 | 0.02440221 | 1 |
| 409 | Crnde | 158 | -0.2594591 | 0.00099472 | 0.02440221 | 1 |
| 201 | Atp5o | 160 | -0.2578339 | 0.00099607 | 0.02441196 | 1 |
| 1056 | Ndrg3 | 160 | -0.2577589 | 0.00099959 | 0.02447455 | 1 |
| 55 | 9130024F11Rik | 160 | -0.257657 | 0.00100438 | 0.02456835 | 1 |
| 479 | Drd3 | 160 | 0.25762787 | 0.00100576 | 0.0245784 | 1 |
| 102 | Adrb2 | 155 | 0.26156792 | 0.00101059 | 0.02466839 | 1 |
| 1107 | Nxn | 160 | -0.2575092 | 0.00101138 | 0.02466839 | 1 |
| 957 | Matn1 | 152 | 0.26403046 | 0.00101318 | 0.02468868 | 1 |
| 1547 | Tbx22 | 159 | -0.2580713 | 0.00102238 | 0.02488234 | 1 |
| 1048 | Nat2 | 155 | 0.26128256 | 0.00102401 | 0.02488234 | 1 |
| 1376 | Serpina10 | 152 | 0.26378294 | 0.00102474 | 0.02488234 | 1 |
| 682 | Gpr156 | 160 | -0.2572047 | 0.00102593 | 0.02488234 | 1 |
| 1089 | Nr4a2 | 159 | 0.25798901 | 0.00102632 | 0.02488234 | 1 |
| 539 | Esr1 | 160 | 0.25718253 | 0.00102699 | 0.02488234 | 1 |
| 22 | 2610008E11Rik | 159 | -0.2578209 | 0.00103441 | 0.02502811 | 1 |
| 1638 | Uap1l1 | 158 | -0.2586 | 0.00103539 | 0.02502811 | 1 |
| 432 | Dab2 | 139 | -0.2753424 | 0.00103596 | 0.02502811 | 1 |
| 1723 | Zcchc8 | 158 | 0.2585031 | 0.00104007 | 0.02510368 | 1 |
| 1174 | Pdlim4 | 159 | -0.2574905 | 0.00105049 | 0.02533113 | 1 |
| 785 | Ina | 152 | -0.2631296 | 0.00105582 | 0.02543569 | 1 |
| 1145 | P2ry2 | 160 | -0.2564603 | 0.0010623 | 0.02555402 | 1 |
| 826 | Kctd10 | 155 | -0.2604585 | 0.00106369 | 0.02555402 | 1 |
| 1160 | Pcdhb20 | 160 | -0.2564312 | 0.00106375 | 0.02555402 | 1 |
| 12 | 1810065E05Rik | 156 | 0.25959955 | 0.00106559 | 0.02557405 | 1 |
| 1624 | Ttc16 | 150 | -0.2644951 | 0.00107267 | 0.02571973 | 1 |
| 586 | Fbxw17 | 155 | 0.26014767 | 0.00107902 | 0.02584772 | 1 |
| 1755 | Zufsp | 156 | -0.2592513 | 0.00108285 | 0.02591517 | 1 |
| 1101 | Nudcd2 | 153 | 0.26159265 | 0.0010898 | 0.02605705 | 1 |
| 937 | Mab21l1 | 160 | -0.2558292 | 0.00109406 | 0.02612631 | 1 |
| 413 | Cstf2 | 160 | -0.2557625 | 0.00109747 | 0.02612631 | 1 |
| 1343 | Rps4x | 159 | -0.2565371 | 0.00109817 | 0.02612631 | 1 |
| 1496 | Ssna1 | 158 | -0.2573239 | 0.00109862 | 0.02612631 | 1 |
| 1584 | Tmem132a | 159 | -0.2565212 | 0.00109898 | 0.02612631 | 1 |
| 1034 | Myo7a | 158 | -0.2573136 | 0.00109915 | 0.02612631 | 1 |
| 670 | Gnb3 | 158 | 0.25727344 | 0.0011012 | 0.02612631 | 1 |
| 1095 | Nt5dc2 | 156 | 0.25886727 | 0.00110219 | 0.02612631 | 1 |
| 483 | Dtd1 | 160 | -0.2556663 | 0.0011024 | 0.02612631 | 1 |
| 1266 | Ptn | 160 | -0.2556554 | 0.00110296 | 0.02612631 | 1 |
| 21 | 2610005L07Rik | 158 | -0.2571809 | 0.00110592 | 0.02617215 | 1 |

| | | | | | | |
|---|---|---|---|---|---|---|
| 370 | Cml3 | 155 | -0.2594859 | 0.00111233 | 0.02629937 | 1 |
| 526 | Entpd1 | 155 | 0.25945549 | 0.00111388 | 0.02631165 | 1 |
| 1063 | Nek2 | 160 | -0.2553673 | 0.00111786 | 0.02638098 | 1 |
| 1195 | Pifo | 159 | -0.2561084 | 0.00112025 | 0.02641302 | 1 |
| 1255 | Psap | 160 | -0.2552914 | 0.00112181 | 0.02642147 | 1 |
| 402 | Cr1l | 158 | -0.256856 | 0.00112268 | 0.02642147 | 1 |
| 1096 | Ntrk1 | 159 | 0.25601201 | 0.00112527 | 0.02643841 | 1 |
| 1554 | Tcta | 156 | 0.25841288 | 0.00112548 | 0.02643841 | 1 |
| 1344 | Rps5 | 160 | 0.25512847 | 0.00113034 | 0.0265059 | 1 |
| 1604 | Tnk1 | 155 | 0.25913384 | 0.00113044 | 0.0265059 | 1 |
| 1064 | Nek5 | 155 | 0.25907425 | 0.00113353 | 0.02655392 | 1 |
| 765 | Hyou1 | 158 | -0.2566009 | 0.001136 | 0.02658563 | 1 |
| 1586 | Tmem14c | 154 | 0.25981904 | 0.00113764 | 0.02658563 | 1 |
| 1126 | Olfr646 | 158 | -0.2565626 | 0.00113801 | 0.02658563 | 1 |
| 1 | 1110032A03Rik | 151 | 0.26228104 | 0.00114061 | 0.02662197 | 1 |
| 1196 | Pigf | 155 | 0.25889355 | 0.00114295 | 0.026652 | 1 |
| 245 | C130036L24Rik | 158 | 0.25644167 | 0.00114439 | 0.02666119 | 1 |
| 136 | Ankrd6 | 160 | -0.2548278 | 0.00114625 | 0.02668015 | 1 |
| 298 | Cd5 | 155 | 0.25878914 | 0.00114842 | 0.02669378 | 1 |
| 324 | Cela1 | 155 | 0.25876534 | 0.00114967 | 0.02669378 | 1 |
| 1332 | Rnf170 | 156 | 0.25794409 | 0.00114998 | 0.02669378 | 1 |
| 446 | Ddx59 | 159 | -0.2555058 | 0.00115198 | 0.02671581 | 1 |
| 1059 | Necab3 | 158 | -0.2562287 | 0.0011557 | 0.0267548 | 1 |
| 98 | Adh7 | 156 | 0.25783481 | 0.00115576 | 0.0267548 | 1 |
| 441 | Dcxr | 155 | 0.25857488 | 0.00115973 | 0.02678199 | 1 |
| 706 | Gtf2h1 | 156 | 0.25775709 | 0.00115989 | 0.02678199 | 1 |
| 154 | Appl2 | 160 | -0.2545692 | 0.00116009 | 0.02678199 | 1 |
| 749 | Hoxd9 | 160 | -0.2545133 | 0.0011631 | 0.02682345 | 1 |
| 1109 | Nxph1 | 160 | -0.2544968 | 0.00116399 | 0.02682345 | 1 |
| 1477 | Sorl1 | 160 | -0.2544212 | 0.00116808 | 0.02689325 | 1 |
| 776 | Igfbp5 | 160 | -0.2543544 | 0.0011717 | 0.02695233 | 1 |
| 677 | Gpr108 | 160 | -0.2543033 | 0.00117448 | 0.02697183 | 1 |
| 35 | 4921531P07Rik | 160 | -0.2542998 | 0.00117467 | 0.02697183 | 1 |
| 1720 | Zcchc17 | 159 | -0.2550205 | 0.00117812 | 0.0270127 | 1 |
| 529 | Epb4.1l5 | 154 | 0.2590439 | 0.00117857 | 0.0270127 | 1 |
| 915 | LOC545466 | 158 | 0.25575041 | 0.00118146 | 0.02705452 | 1 |
| 1525 | Syt1 | 160 | -0.2540748 | 0.00118698 | 0.02715646 | 1 |
| 270 | Cbx2 | 156 | 0.25713828 | 0.00119323 | 0.02727516 | 1 |
| 1091 | Nrn1 | 160 | -0.2538808 | 0.00119768 | 0.02735229 | 1 |
| 1595 | Tmem248 | 155 | 0.25767636 | 0.00120829 | 0.02756975 | 1 |
| 558 | Fam206a | 158 | 0.25522447 | 0.00121039 | 0.02759307 | 1 |
| 1347 | Rpsa | 159 | -0.2543436 | 0.00121549 | 0.02766685 | 1 |
| 181 | Arsg | 156 | -0.2567284 | 0.0012158 | 0.02766685 | 1 |
| 1475 | Sorbs3 | 160 | -0.2534922 | 0.00121939 | 0.02770543 | 1 |
| 896 | LOC432906 | 159 | -0.2542527 | 0.0012206 | 0.02770543 | 1 |

| | | | | | | |
|---|---|---|---|---|---|---|
| 1375 | 5-Sep | 72 | 0.37375617 | 0.00122076 | 0.02770543 | 1 |
| 835 | Kif9 | 158 | -0.254965 | 0.0012249 | 0.02777469 | 1 |
| 900 | LOC433525 | 160 | -0.2533364 | 0.0012282 | 0.02782465 | 1 |
| 129 | Alpk2 | 160 | -0.2533125 | 0.00122956 | 0.0278306 | 1 |
| 255 | Cacnb2 | 158 | -0.2548262 | 0.00123274 | 0.02787781 | 1 |
| 1263 | Pthlh | 158 | -0.2547309 | 0.00123814 | 0.02797508 | 1 |
| 79 | Acbd3 | 158 | -0.2546844 | 0.00124078 | 0.02799083 | 1 |
| 385 | Col5a3 | 160 | -0.2531112 | 0.00124103 | 0.02799083 | 1 |
| 43 | 4933434E20Rik | 158 | -0.2546189 | 0.00124451 | 0.02801899 | 1 |
| 741 | Hmgcs2 | 154 | 0.25782936 | 0.00124542 | 0.02801899 | 1 |
| 74 | Ablim1 | 158 | -0.2546002 | 0.00124558 | 0.02801899 | 1 |
| 1562 | Tex15 | 158 | -0.2545789 | 0.0012468 | 0.0280216 | 1 |
| 1754 | Zscan4a | 152 | 0.25942639 | 0.00124894 | 0.02802282 | 1 |
| 292 | Ccser2 | 158 | -0.2544818 | 0.00125236 | 0.02802282 | 1 |
| 1338 | Rplp0 | 160 | -0.2529087 | 0.00125267 | 0.02802282 | 1 |
| 1682 | Vwa5a | 155 | 0.25688299 | 0.0012527 | 0.02802282 | 1 |
| 455 | Dhx30 | 154 | 0.25769378 | 0.00125309 | 0.02802282 | 1 |
| 763 | Htra2 | 158 | -0.2544627 | 0.00125346 | 0.02802282 | 1 |
| 633 | Gata3 | 160 | -0.252816 | 0.00125803 | 0.02808169 | 1 |
| 1065 | Nek7 | 156 | -0.2559752 | 0.00125829 | 0.02808169 | 1 |
| 801 | Itga8 | 160 | -0.2527834 | 0.00125992 | 0.02809345 | 1 |
| 1359 | S100b | 160 | -0.2526594 | 0.00126714 | 0.0282296 | 1 |
| 597 | Folr2 | 158 | 0.25420211 | 0.00126851 | 0.02823535 | 1 |
| 376 | Cnst | 153 | -0.2582272 | 0.00126972 | 0.02823767 | 1 |
| 1688 | Wdtc1 | 156 | -0.2557423 | 0.00127171 | 0.0282572 | 1 |
| 1396 | Shh | 156 | 0.25566456 | 0.00127621 | 0.02833257 | 1 |
| 1424 | Slc27a3 | 159 | -0.2532605 | 0.00127756 | 0.02833773 | 1 |
| 1247 | Prkcq | 159 | -0.2532184 | 0.00128003 | 0.02836785 | 1 |
| 312 | Cdh7 | 158 | -0.2539541 | 0.00128298 | 0.02840863 | 1 |
| 1349 | Rrp12 | 158 | 0.25393312 | 0.00128421 | 0.02841114 | 1 |
| 1615 | Trh | 155 | 0.25630703 | 0.00128587 | 0.02842306 | 1 |
| 1222 | Polk | 158 | -0.2538232 | 0.00129069 | 0.02850479 | 1 |
| 1027 | Mup5 | 160 | -0.2521994 | 0.00129424 | 0.02855849 | 1 |
| 882 | LOC231914 | 155 | 0.2561129 | 0.00129723 | 0.02859969 | 1 |
| 1253 | Prpsap2 | 160 | 0.25209806 | 0.00130028 | 0.02862985 | 1 |
| 1188 | Pgm3 | 158 | -0.2536516 | 0.00130085 | 0.02862985 | 1 |
| 1272 | Qsox1 | 158 | -0.2535979 | 0.00130404 | 0.02867537 | 1 |
| 170 | Arhgef2 | 160 | -0.2519784 | 0.00130745 | 0.02872556 | 1 |
| 1642 | Ube2t | 160 | 0.25175892 | 0.00132069 | 0.02896547 | 1 |
| 343 | Cherp | 158 | -0.2533199 | 0.00132069 | 0.02896547 | 1 |
| 1000 | Mpi | 159 | -0.2525177 | 0.00132178 | 0.02896547 | 1 |
| 415 | Ctf1 | 158 | 0.25327602 | 0.00132334 | 0.02897458 | 1 |
| 1567 | Tgm3 | 158 | -0.2532153 | 0.00132701 | 0.02901519 | 1 |
| 569 | Fanci | 159 | -0.2523966 | 0.00132913 | 0.02901519 | 1 |
| 1200 | Pitx1 | 159 | -0.2523931 | 0.00132933 | 0.02901519 | 1 |

| | | | | | | |
|---|---|---|---|---|---|---|
| 576 | Fbln1 | 160 | -0.2516098 | 0.00132975 | 0.02901519 | 1 |
| 332 | Cep57 | 158 | -0.2531377 | 0.00133171 | 0.02903304 | 1 |
| 1406 | Slc15a1 | 147 | -0.2621878 | 0.00133646 | 0.02911183 | 1 |
| 1348 | Rrm1 | 159 | -0.252211 | 0.00134044 | 0.02917357 | 1 |
| 1591 | Tmem216 | 158 | -0.2529224 | 0.00134484 | 0.02924419 | 1 |
| 1233 | Ppm1f | 147 | -0.2619561 | 0.00135018 | 0.02933531 | 1 |
| 1660 | Utp3 | 159 | -0.2519322 | 0.00135762 | 0.02945179 | 1 |
| 1084 | Npdc1 | 160 | -0.2511339 | 0.00135906 | 0.02945179 | 1 |
| 911 | LOC434781 | 157 | -0.2534764 | 0.00135934 | 0.02945179 | 1 |
| 1191 | Phf20l1 | 147 | -0.2617761 | 0.00136093 | 0.02945179 | 1 |
| 73 | Abl1 | 159 | -0.2518725 | 0.00136132 | 0.02945179 | 1 |
| 342 | Chd2 | 160 | -0.2509992 | 0.00136747 | 0.0295454 | 1 |
| 895 | LOC432871 | 149 | -0.25992 | 0.00136851 | 0.0295454 | 1 |
| 1521 | Syne1 | 160 | -0.2509551 | 0.00137022 | 0.0295454 | 1 |
| 1709 | Ywhab | 160 | -0.250954 | 0.00137029 | 0.0295454 | 1 |
| 919 | LOC546041 | 159 | -0.2515637 | 0.00138063 | 0.0297432 | 1 |
| 845 | Klrb1b | 160 | -0.250733 | 0.00138421 | 0.02976033 | 1 |
| 258 | Calcoco1 | 160 | -0.2507218 | 0.00138492 | 0.02976033 | 1 |
| 1319 | Ring1 | 157 | -0.253056 | 0.0013855 | 0.02976033 | 1 |
| 444 | Ddx50 | 158 | -0.252258 | 0.0013861 | 0.02976033 | 1 |
| 1581 | Tmem109 | 145 | -0.2630309 | 0.00139166 | 0.02985449 | 1 |
| 1231 | Ppm1a | 160 | -0.2505962 | 0.00139289 | 0.02985559 | 1 |
| 1316 | Rhox9 | 158 | -0.2519948 | 0.00140276 | 0.02997825 | 1 |
| 172 | Arl10 | 159 | 0.25121336 | 0.00140284 | 0.02997825 | 1 |
| 322 | Cds2 | 160 | -0.2504389 | 0.00140293 | 0.02997825 | 1 |
| 1501 | Stat3 | 157 | 0.25277373 | 0.00140332 | 0.02997825 | 1 |
| 1491 | Sptb | 160 | -0.2503997 | 0.00140544 | 0.02999835 | 1 |
| 1261 | Ptchd1 | 160 | -0.2503235 | 0.00141033 | 0.03007418 | 1 |
| 1466 | Sntn | 159 | -0.2510658 | 0.00141228 | 0.03007418 | 1 |
| 560 | Fam214b | 158 | -0.2518419 | 0.00141253 | 0.03007418 | 1 |
| 1103 | Nudt2 | 155 | -0.2541859 | 0.00141506 | 0.03009086 | 1 |
| 802 | Itih3 | 156 | 0.25337428 | 0.00141568 | 0.03009086 | 1 |
| 430 | Cysltr1 | 155 | -0.254156 | 0.00141697 | 0.0300932 | 1 |
| 731 | Helb | 158 | -0.2516897 | 0.00142231 | 0.03015085 | 1 |
| 134 | Ank | 159 | -0.2508967 | 0.00142319 | 0.03015085 | 1 |
| 487 | Dyx1c1 | 156 | -0.2532561 | 0.00142324 | 0.03015085 | 1 |
| 383 | Col3a1 | 158 | -0.2514802 | 0.00143588 | 0.03036475 | 1 |
| 399 | Cpe | 155 | -0.2538464 | 0.00143681 | 0.03036475 | 1 |
| 1543 | Tbl2 | 160 | 0.2499139 | 0.00143691 | 0.03036475 | 1 |
| 822 | Kcnj16 | 160 | 0.24987131 | 0.0014397 | 0.03037759 | 1 |
| 685 | Gpr179 | 159 | -0.2506397 | 0.00143991 | 0.03037759 | 1 |
| 454 | Dhfr | 156 | 0.25297515 | 0.00144136 | 0.03038315 | 1 |
| 656 | Gm3579 | 156 | -0.2528515 | 0.0014494 | 0.03052735 | 1 |
| 1737 | Zfp692 | 159 | -0.2504511 | 0.00145228 | 0.03055362 | 1 |
| 1663 | Vamp3 | 159 | -0.2504358 | 0.00145329 | 0.03055362 | 1 |

| | | | | | | |
|---|---|---|---|---|---|---|
| 1225 | Pomt2 | 156 | 0.25277732 | 0.00145425 | 0.03055362 | 1 |
| 610 | Fzd9 | 160 | -0.2495463 | 0.00146116 | 0.03067348 | 1 |
| 1725 | Zdhhc4 | 154 | 0.2542605 | 0.00146239 | 0.03067412 | 1 |
| 420 | Cul3 | 160 | -0.2494765 | 0.0014658 | 0.03072032 | 1 |
| 1180 | Peg3 | 159 | -0.250184 | 0.00146998 | 0.03078252 | 1 |
| 240 | Brd3 | 154 | 0.25407766 | 0.00147439 | 0.03080729 | 1 |
| 247 | C2cd2 | 156 | -0.252451 | 0.00147573 | 0.03080729 | 1 |
| 1153 | Pank2 | 147 | -0.2599156 | 0.00147666 | 0.03080729 | 1 |
| 943 | Maged2 | 155 | 0.25321151 | 0.0014783 | 0.03080729 | 1 |
| 832 | Kif12 | 156 | 0.25240109 | 0.00147905 | 0.03080729 | 1 |
| 1572 | Tlcd2 | 160 | -0.2492698 | 0.00147963 | 0.03080729 | 1 |
| 421 | Cwc22 | 156 | -0.2523642 | 0.0014815 | 0.03080729 | 1 |
| 809 | Jarid2 | 156 | 0.25236366 | 0.00148154 | 0.03080729 | 1 |
| 819 | Kcne2 | 160 | -0.2492339 | 0.00148205 | 0.03080729 | 1 |
| 251 | C87436 | 154 | 0.25392572 | 0.00148442 | 0.03083143 | 1 |
| 100 | Adipor2 | 159 | -0.249948 | 0.00148578 | 0.03083451 | 1 |
| 63 | A830039N20Rik | 160 | -0.2491498 | 0.00148772 | 0.03084963 | 1 |
| 1609 | Tpcn1 | 146 | -0.2605051 | 0.00149499 | 0.03097508 | 1 |
| 1054 | Nde1 | 160 | -0.2489608 | 0.00150054 | 0.03106491 | 1 |
| 336 | Cfap69 | 146 | -0.2603573 | 0.00150463 | 0.03112423 | 1 |
| 566 | Fam65b | 160 | -0.2488803 | 0.00150603 | 0.03112794 | 1 |
| 1245 | Prelp | 155 | -0.2526518 | 0.00151577 | 0.03130388 | 1 |
| 1360 | Samd11 | 145 | -0.261035 | 0.0015179 | 0.03132227 | 1 |
| 242 | Bspry | 156 | 0.25178434 | 0.00152053 | 0.03135134 | 1 |
| 851 | Krt71 | 158 | -0.2501796 | 0.00152276 | 0.03136547 | 1 |
| 418 | Cubn | 156 | -0.2517381 | 0.00152368 | 0.03136547 | 1 |
| 1133 | Orai2 | 159 | -0.2492583 | 0.00153286 | 0.03152881 | 1 |
| 1285 | Rap1b | 158 | -0.2500149 | 0.0015341 | 0.03152893 | 1 |
| 630 | Gas6 | 160 | -0.2483889 | 0.00153994 | 0.03162347 | 1 |
| 859 | Laptm4b | 160 | -0.2482941 | 0.00154657 | 0.03173396 | 1 |
| 326 | Cenpa | 155 | 0.25209759 | 0.00155374 | 0.03181802 | 1 |
| 1012 | Mrrf | 156 | -0.251291 | 0.00155448 | 0.03181802 | 1 |
| 978 | Mfsd7a | 160 | -0.2481791 | 0.00155463 | 0.03181802 | 1 |
| 1715 | Zbp1 | 152 | 0.25449136 | 0.00155649 | 0.03181802 | 1 |
| 583 | Fbxo7 | 160 | -0.2481467 | 0.00155691 | 0.03181802 | 1 |
| 1067 | Nfatc1 | 156 | 0.25123168 | 0.00155861 | 0.03182713 | 1 |
| 744 | Hnrnph2 | 160 | -0.2480718 | 0.0015622 | 0.03187487 | 1 |
| 11 | 1810030O07Rik | 146 | -0.2594534 | 0.00156487 | 0.03190388 | 1 |
| 1623 | Tsx | 156 | 0.25111227 | 0.00156695 | 0.03192065 | 1 |
| 450 | Desi1 | 160 | -0.2479594 | 0.00157016 | 0.03196045 | 1 |
| 365 | Clk2 | 159 | -0.2486924 | 0.00157249 | 0.03198229 | 1 |
| 1552 | Tcirg1 | 160 | 0.2479069 | 0.00157389 | 0.03198517 | 1 |
| 834 | Kif18a | 155 | 0.25169521 | 0.00158184 | 0.0321212 | 1 |
| 1324 | Ripply3 | 160 | 0.24773739 | 0.00158598 | 0.03217303 | 1 |
| 793 | Irx2 | 160 | -0.2477154 | 0.00158756 | 0.03217303 | 1 |

| | | | | | | |
|---|---|---|---|---|---|---|
| 1704 | Yif1b | 151 | -0.2548602 | 0.00158818 | 0.03217303 | 1 |
| 838 | Klf8 | 158 | -0.2491847 | 0.00159243 | 0.03222085 | 1 |
| 16 | 2310033P09Rik | 158 | 0.24917576 | 0.00159307 | 0.03222085 | 1 |
| 199 | Atp5c1 | 158 | -0.2490606 | 0.00160133 | 0.03236206 | 1 |
| 1014 | Msantd2 | 157 | -0.249772 | 0.00160618 | 0.03243438 | 1 |
| 695 | Grina | 156 | -0.2504725 | 0.00161233 | 0.03252858 | 1 |
| 297 | Cd34 | 155 | -0.2512447 | 0.00161385 | 0.03252858 | 1 |
| 902 | LOC433602 | 157 | -0.249654 | 0.00161468 | 0.03252858 | 1 |
| 1051 | Ncald | 160 | -0.2472639 | 0.00162023 | 0.03261457 | 1 |
| 257 | Calca | 160 | -0.2471148 | 0.00163115 | 0.03280846 | 1 |
| 1565 | Tgfb1 | 160 | -0.2469798 | 0.00164109 | 0.03295524 | 1 |
| 76 | Acadl | 160 | -0.2469606 | 0.00164251 | 0.03295524 | 1 |
| 1011 | Mrps5 | 143 | 0.26098062 | 0.00164263 | 0.03295524 | 1 |
| 438 | Dcdc5 | 158 | -0.2484788 | 0.00164362 | 0.03295524 | 1 |
| 571 | Farsa | 159 | -0.247619 | 0.00165024 | 0.03306197 | 1 |
| 124 | Aldoa | 158 | -0.2483672 | 0.00165184 | 0.03306805 | 1 |
| 854 | Lag3 | 155 | -0.2506892 | 0.00165414 | 0.03308795 | 1 |
| 827 | Kctd3 | 138 | -0.2653733 | 0.00165705 | 0.03312015 | 1 |
| 930 | Ltf | 155 | 0.2505538 | 0.0016641 | 0.03321378 | 1 |
| 229 | Bglap3 | 153 | 0.25213882 | 0.0016656 | 0.03321378 | 1 |
| 1310 | Rhbdd2 | 155 | 0.25053281 | 0.00166565 | 0.03321378 | 1 |
| 215 | Bap1 | 159 | -0.2473924 | 0.0016671 | 0.03321686 | 1 |
| 1265 | Ptn | 158 | -0.2480887 | 0.00167254 | 0.03327359 | 1 |
| 547 | Ezr | 160 | -0.2465575 | 0.00167257 | 0.03327359 | 1 |
| 673 | Got2 | 159 | -0.2472028 | 0.00168132 | 0.03342171 | 1 |
| 359 | Cldn6 | 155 | 0.25028541 | 0.001684 | 0.03342894 | 1 |
| 1456 | Sms | 155 | 0.25028116 | 0.00168431 | 0.03342894 | 1 |
| 86 | Acox3 | 138 | -0.2649081 | 0.00168975 | 0.03351074 | 1 |
| 1223 | Polr3k | 160 | -0.2463043 | 0.0016917 | 0.03352322 | 1 |
| 204 | Atp6v0e | 158 | -0.2477744 | 0.00169619 | 0.0335407 | 1 |
| 10 | 1810024B03Rik | 158 | -0.247771 | 0.00169644 | 0.0335407 | 1 |
| 631 | Gata2 | 160 | -0.2462407 | 0.00169653 | 0.0335407 | 1 |
| 1575 | Tm4sf1 | 159 | -0.2469557 | 0.00170003 | 0.03355713 | 1 |
| 647 | Glrb | 160 | -0.2461773 | 0.00170137 | 0.03355713 | 1 |
| 517 | Emc2 | 158 | -0.2477041 | 0.00170152 | 0.03355713 | 1 |
| 33 | 4833427G06Rik | 157 | -0.2484541 | 0.00170341 | 0.03355713 | 1 |
| 307 | Cdh18 | 158 | -0.247672 | 0.00170395 | 0.03355713 | 1 |
| 155 | Aprt | 156 | 0.24915802 | 0.00170935 | 0.03363749 | 1 |
| 1699 | Wwp2 | 153 | -0.2514951 | 0.00171344 | 0.03369187 | 1 |
| 1166 | Pde3b | 160 | -0.245996 | 0.00171526 | 0.03370177 | 1 |
| 1230 | Ppil6 | 158 | -0.2474976 | 0.00171725 | 0.03371483 | 1 |
| 512 | Elmo2 | 158 | -0.2473861 | 0.00172581 | 0.03383861 | 1 |
| 750 | Hpse2 | 159 | -0.2466139 | 0.00172622 | 0.03383861 | 1 |
| 1346 | Rps9 | 160 | -0.2457917 | 0.00173104 | 0.03389908 | 1 |
| 1258 | Psmc3 | 159 | -0.2465395 | 0.00173196 | 0.03389908 | 1 |

| | | | | | | |
|---|---|---|---|---|---|---|
| 903 | LOC433610 | 159 | -0.246477 | 0.0017368 | 0.03396774 | 1 |
| 664 | Gm6878 | 158 | -0.247186 | 0.00174126 | 0.03402696 | 1 |
| 1428 | Slc35a1 | 160 | -0.2456443 | 0.00174251 | 0.03402696 | 1 |
| 1264 | Ptn | 160 | -0.2456152 | 0.00174478 | 0.03404323 | 1 |
| 642 | Gin1 | 155 | 0.24946709 | 0.00174601 | 0.03404323 | 1 |
| 50 | 6330581N18Rik | 159 | -0.246285 | 0.00175175 | 0.03412905 | 1 |
| 884 | LOC381076 | 160 | 0.24548332 | 0.00175511 | 0.03416055 | 1 |
| 308 | Cdh22 | 159 | -0.2462301 | 0.00175605 | 0.03416055 | 1 |
| 984 | Mir99ahg | 158 | -0.2469498 | 0.00175966 | 0.03420464 | 1 |
| 813 | Katnb1 | 147 | -0.2558348 | 0.00176257 | 0.03423508 | 1 |
| 923 | Lpar1 | 160 | -0.2451649 | 0.00178028 | 0.03453434 | 1 |
| 1415 | Slc20a2 | 159 | -0.2459176 | 0.00178069 | 0.03453434 | 1 |
| 1022 | Mtus2 | 160 | -0.2450888 | 0.00178634 | 0.03461595 | 1 |
| 842 | Klhl21 | 143 | 0.258995 | 0.00178762 | 0.03461595 | 1 |
| 941 | Mafg | 158 | 0.24651333 | 0.00179413 | 0.03468227 | 1 |
| 866 | Lefty2 | 156 | -0.2480571 | 0.00179467 | 0.03468227 | 1 |
| 972 | Mepce | 155 | 0.24882667 | 0.00179599 | 0.03468227 | 1 |
| 1002 | Mptx1 | 160 | 0.24495049 | 0.00179742 | 0.03468227 | 1 |
| 961 | Mbl2 | 156 | 0.2480169 | 0.00179786 | 0.03468227 | 1 |
| 5 | 1700036A12Rik | 156 | -0.2479171 | 0.00180579 | 0.03475963 | 1 |
| 333 | Cfap126 | 158 | -0.2463668 | 0.00180584 | 0.03475963 | 1 |
| 88 | Acta1 | 160 | -0.2448443 | 0.00180596 | 0.03475963 | 1 |
| 1576 | Tm4sf1 | 160 | -0.2447965 | 0.00180981 | 0.03478755 | 1 |
| 19 | 2310057J18Rik | 159 | 0.24554926 | 0.00181014 | 0.03478755 | 1 |
| 1173 | Pdia4 | 155 | -0.2485987 | 0.00181409 | 0.03482318 | 1 |
| 355 | Ckmt2 | 159 | -0.2454724 | 0.00181634 | 0.03482318 | 1 |
| 608 | Fxyd3 | 158 | -0.2462206 | 0.00181759 | 0.03482318 | 1 |
| 1529 | Syt7 | 160 | -0.2446828 | 0.00181902 | 0.03482318 | 1 |
| 1159 | Pcdhb16 | 160 | -0.244671 | 0.00181998 | 0.03482318 | 1 |
| 442 | Ddx19b | 160 | -0.2446682 | 0.0018202 | 0.03482318 | 1 |
| 1161 | Pcdhgb6 | 153 | -0.2500015 | 0.0018293 | 0.03497094 | 1 |
| 587 | Fcgr2b | 158 | -0.2457744 | 0.00185388 | 0.03541434 | 1 |
| 543 | Etl4 | 157 | -0.2464313 | 0.00186312 | 0.03555036 | 1 |
| 26 | 2810007J24Rik | 156 | 0.24719946 | 0.0018638 | 0.03555036 | 1 |
| 1492 | Sqrdl | 160 | -0.244113 | 0.00186579 | 0.03556177 | 1 |
| 1357 | S100b | 160 | -0.2440835 | 0.00186824 | 0.03556796 | 1 |
| 753 | Hsd17b7 | 159 | -0.2448243 | 0.0018694 | 0.03556796 | 1 |
| 105 | Agpat3 | 160 | -0.2440587 | 0.00187031 | 0.03556796 | 1 |
| 696 | Grip2 | 160 | -0.2439681 | 0.00187787 | 0.03566237 | 1 |
| 1007 | Mrpl12 | 158 | 0.24548148 | 0.00187807 | 0.03566237 | 1 |
| 143 | Ap3b1 | 156 | -0.2469949 | 0.00188064 | 0.03568446 | 1 |
| 1129 | Olfr984 | 159 | -0.2445592 | 0.00189151 | 0.03586399 | 1 |
| 1142 | Oxct1 | 158 | -0.245251 | 0.0018973 | 0.03594714 | 1 |
| 423 | Cxcl13 | 160 | -0.2436817 | 0.00190192 | 0.0359825 | 1 |
| 1207 | Plcxd2 | 160 | -0.2436808 | 0.001902 | 0.0359825 | 1 |

| | | | | | | |
|---|---|---|---|---|---|---|
| 1494 | Srsf11 | 159 | -0.2443879 | 0.00190592 | 0.03603 | 1 |
| 489 | E2f1 | 160 | -0.2435952 | 0.00190925 | 0.03606613 | 1 |
| 1448 | Sln | 160 | -0.2435756 | 0.00191091 | 0.03607064 | 1 |
| 1650 | Uhrf1 | 151 | 0.25052834 | 0.00191832 | 0.0361838 | 1 |
| 1487 | Spg21 | 160 | -0.2434467 | 0.00192188 | 0.03620863 | 1 |
| 1656 | Usp19 | 160 | -0.2434396 | 0.00192248 | 0.03620863 | 1 |
| 1281 | Rala | 156 | 0.24641612 | 0.00192905 | 0.03629682 | 1 |
| 1616 | Trim39 | 155 | 0.24718607 | 0.00193001 | 0.03629682 | 1 |
| 638 | Gfra2 | 158 | -0.2448043 | 0.00193508 | 0.03635122 | 1 |
| 377 | Cntn4 | 158 | -0.2447964 | 0.00193576 | 0.03635122 | 1 |
| 1518 | Sybu | 158 | -0.2447618 | 0.00193872 | 0.03637995 | 1 |
| 872 | Lhfpl2 | 158 | -0.2447376 | 0.00194079 | 0.03639192 | 1 |
| 405 | Crb1 | 159 | -0.243871 | 0.00195002 | 0.03653808 | 1 |
| 1707 | Yjefn3 | 158 | -0.244576 | 0.00195466 | 0.03659814 | 1 |
| 1570 | Timm50 | 155 | 0.24685426 | 0.0019582 | 0.03662462 | 1 |
| 192 | Atg4b | 159 | -0.2437457 | 0.00196084 | 0.03662462 | 1 |
| 114 | Akna | 158 | -0.2444961 | 0.00196155 | 0.03662462 | 1 |
| 585 | Fbxo9 | 160 | -0.2429827 | 0.00196183 | 0.03662462 | 1 |
| 1300 | Rest | 159 | -0.243705 | 0.00196437 | 0.03664525 | 1 |
| 963 | Mcat | 159 | -0.2436574 | 0.0019685 | 0.03668191 | 1 |
| 1149 | Pafah1b1 | 160 | -0.2428979 | 0.00196922 | 0.03668191 | 1 |
| 567 | Fam73b | 157 | -0.245127 | 0.0019732 | 0.03671258 | 1 |
| 1280 | Rag2 | 157 | -0.2451207 | 0.00197375 | 0.03671258 | 1 |
| 44 | 4933435E02Rik | 157 | -0.2450992 | 0.00197561 | 0.03672034 | 1 |
| 393 | Cox6c | 158 | -0.2442945 | 0.00197904 | 0.03674793 | 1 |
| 1677 | Vti1a | 160 | -0.2427748 | 0.00197998 | 0.03674793 | 1 |
| 701 | Gsc | 152 | -0.2488747 | 0.00198946 | 0.03689697 | 1 |
| 954 | Marcksl1 | 16 | -0.7115974 | 0.00199113 | 0.03690112 | 1 |
| 556 | Fam178b | 154 | 0.24722022 | 0.00199448 | 0.03693629 | 1 |
| 678 | Gpr133 | 160 | -0.2425848 | 0.00199669 | 0.03695043 | 1 |
| 1400 | Sin3a | 159 | -0.2433163 | 0.00199836 | 0.03695451 | 1 |
| 1013 | Mrs2 | 158 | -0.2440227 | 0.00200285 | 0.03701051 | 1 |
| 495 | Ecel1 | 134 | 0.26457866 | 0.00200648 | 0.0370441 | 1 |
| 431 | D10Jhu81e | 156 | 0.24550467 | 0.00200757 | 0.0370441 | 1 |
| 1250 | Prlr | 154 | 0.24704043 | 0.00201012 | 0.03705329 | 1 |
| 506 | Eif3j1 | 157 | -0.2446946 | 0.00201098 | 0.03705329 | 1 |
| 1486 | Speer1-ps1 | 159 | -0.2431491 | 0.00201315 | 0.03706643 | 1 |
| 1612 | Trap1a | 154 | 0.24698279 | 0.00201516 | 0.03707658 | 1 |
| 1031 | Mx2 | 160 | 0.24231492 | 0.00202066 | 0.03715098 | 1 |
| 1457 | Smu1 | 160 | -0.2422546 | 0.00202605 | 0.03716627 | 1 |
| 540 | Esyt3 | 160 | 0.24224614 | 0.00202681 | 0.03716627 | 1 |
| 471 | Dnajc4 | 158 | -0.2437481 | 0.00202716 | 0.03716627 | 1 |
| 125 | Alg1 | 136 | -0.2624079 | 0.00202733 | 0.03716627 | 1 |
| 1398 | Siah1b | 156 | -0.2451517 | 0.00203876 | 0.03731007 | 1 |
| 671 | Gon4l | 158 | -0.2436159 | 0.00203896 | 0.03731007 | 1 |

| | | | | | | |
|---|---|---|---|---|---|---|
| 648 | Gm10010 | 160 | 0.24210403 | 0.00203957 | 0.03731007 | 1 |
| 20 | 2610002J02Rik | 159 | -0.2428333 | 0.00204135 | 0.03731578 | 1 |
| 295 | Cct8 | 160 | -0.2420616 | 0.0020434 | 0.03732646 | 1 |
| 36 | 4922502D21Rik | 160 | 0.24203348 | 0.00204593 | 0.03734595 | 1 |
| 1447 | Slco2a1 | 159 | -0.2426996 | 0.00205339 | 0.03742946 | 1 |
| 1605 | Tnni3 | 158 | -0.2434445 | 0.00205435 | 0.03742946 | 1 |
| 940 | Maea | 160 | -0.2419342 | 0.00205492 | 0.03742946 | 1 |
| 598 | Foxb1 | 160 | -0.2418882 | 0.00205909 | 0.03747693 | 1 |
| 1355 | Rxfp3 | 156 | 0.24490886 | 0.00206047 | 0.03747693 | 1 |
| 982 | Mipep | 156 | -0.2448547 | 0.00206534 | 0.03751449 | 1 |
| 1276 | Rab34 | 158 | -0.2433214 | 0.00206548 | 0.03751449 | 1 |
| 428 | Cyp4v3 | 159 | -0.2425389 | 0.00206796 | 0.03753279 | 1 |
| 37 | 4930430A15Rik | 156 | -0.2447877 | 0.00207137 | 0.03756797 | 1 |
| 1100 | Nudc | 160 | -0.2416951 | 0.00207669 | 0.0376318 | 1 |
| 1016 | Msmo1 | 160 | -0.2416768 | 0.00207837 | 0.0376318 | 1 |
| 1511 | Sugp1 | 133 | -0.2646747 | 0.00207933 | 0.0376318 | 1 |
| 1509 | Suco | 158 | -0.2431068 | 0.00208499 | 0.03770755 | 1 |
| 81 | Ace | 160 | -0.2414857 | 0.00209594 | 0.03782903 | 1 |
| 1093 | Nt5c2 | 160 | -0.2414622 | 0.00209812 | 0.03782903 | 1 |
| 1618 | Trmt112 | 158 | -0.2429611 | 0.00209834 | 0.03782903 | 1 |
| 1558 | Tecta | 160 | -0.2414149 | 0.00210249 | 0.03782903 | 1 |
| 193 | Atg5 | 156 | -0.2444333 | 0.00210359 | 0.03782903 | 1 |
| 223 | Bcl10 | 157 | -0.2436588 | 0.00210416 | 0.03782903 | 1 |
| 244 | Btn1a1 | 152 | 0.24756375 | 0.00210511 | 0.03782903 | 1 |
| 1353 | Runx1 | 158 | -0.2428833 | 0.00210549 | 0.03782903 | 1 |
| 955 | Marco | 156 | 0.24441077 | 0.00210565 | 0.03782903 | 1 |
| 595 | Fmn2 | 159 | -0.2421161 | 0.00210672 | 0.03782903 | 1 |
| 1705 | Yipf1 | 160 | -0.2413549 | 0.00210805 | 0.03782903 | 1 |
| 705 | Gsto1 | 160 | -0.2413149 | 0.00211176 | 0.03786502 | 1 |
| 734 | Hhip | 160 | -0.2413013 | 0.00211303 | 0.03786502 | 1 |
| 278 | Ccdc177 | 156 | -0.2442738 | 0.00211823 | 0.03793151 | 1 |
| 1416 | Slc22a12 | 155 | 0.24499123 | 0.00212353 | 0.03799978 | 1 |
| 620 | Gad2 | 158 | 0.24265061 | 0.00212703 | 0.03803566 | 1 |
| 505 | Eif3g | 158 | -0.2426205 | 0.00212983 | 0.03804929 | 1 |
| 1386 | Sfrp2 | 158 | 0.24261032 | 0.00213078 | 0.03804929 | 1 |
| 880 | Lmo4 | 160 | -0.2410536 | 0.00213617 | 0.03811871 | 1 |
| 1752 | Zpr1 | 156 | 0.24397321 | 0.00214609 | 0.03826886 | 1 |
| 284 | Ccdc97 | 159 | -0.2416686 | 0.00214847 | 0.03828452 | 1 |
| 56 | 9130204K15Rik | 160 | -0.2409057 | 0.0021501 | 0.0382868 | 1 |
| 183 | Asb4 | 160 | 0.24088448 | 0.00215211 | 0.03829578 | 1 |
| 372 | Cndp2 | 159 | 0.24159575 | 0.00215534 | 0.03831003 | 1 |
| 291 | Ccrl2 | 156 | 0.24386795 | 0.00215592 | 0.03831003 | 1 |
| 1425 | Slc28a2 | 156 | 0.24381384 | 0.00216099 | 0.03837335 | 1 |
| 1274 | R74862 | 159 | -0.2414225 | 0.00217175 | 0.0385321 | 1 |
| 1414 | Slc20a1 | 160 | -0.240665 | 0.00217295 | 0.0385321 | 1 |

| | | | | | | |
|---|---|---|---|---|---|---|
| 549 | F3 | 152 | 0.24679031 | 0.00217618 | 0.0385624 | 1 |
| 1449 | Smc3 | 159 | -0.241295 | 0.0021839 | 0.03865563 | 1 |
| 301 | Cdadc1 | 159 | -0.241289 | 0.00218447 | 0.03865563 | 1 |
| 697 | Grm4 | 160 | -0.2404978 | 0.00218895 | 0.03870798 | 1 |
| 799 | Itfg3 | 160 | -0.2404723 | 0.0021914 | 0.0387244 | 1 |
| 989 | Mmp14 | 160 | -0.2403875 | 0.00219957 | 0.03884016 | 1 |
| 570 | Fars2 | 155 | 0.24416323 | 0.002201 | 0.03884016 | 1 |
| 1443 | Slc6a8 | 137 | -0.2594418 | 0.00220258 | 0.03884112 | 1 |
| 157 | Aqp4 | 160 | -0.2403189 | 0.00220619 | 0.03887777 | 1 |
| 553 | Fam131b | 158 | -0.2417963 | 0.00220785 | 0.03888007 | 1 |
| 1649 | Ugt2a2 | 158 | -0.2417713 | 0.00221025 | 0.03889552 | 1 |
| 1690 | Wfikkn2 | 158 | -0.2417035 | 0.0022168 | 0.03898379 | 1 |
| 110 | Aifm3 | 160 | -0.2401749 | 0.00222015 | 0.03901424 | 1 |
| 187 | Aspg | 160 | -0.2401601 | 0.00222159 | 0.03901424 | 1 |
| 1345 | Rps6ka6 | 152 | 0.24623855 | 0.0022282 | 0.03910324 | 1 |
| 200 | Atp5e | 160 | -0.2399851 | 0.00223868 | 0.03925529 | 1 |
| 254 | Cab39l | 160 | -0.2399722 | 0.00223994 | 0.03925529 | 1 |
| 480 | Drg2 | 158 | -0.2414282 | 0.00224352 | 0.03929088 | 1 |
| 853 | L3mbtl4 | 160 | -0.2399162 | 0.00224545 | 0.03929765 | 1 |
| 1716 | Zbtb11 | 159 | -0.2406309 | 0.00224819 | 0.0393186 | 1 |
| 1463 | Sncb | 158 | -0.2413426 | 0.00225189 | 0.03935625 | 1 |
| 1070 | Ngfr | 160 | 0.23979628 | 0.00225725 | 0.03942299 | 1 |
| 1472 | Sod3 | 158 | 0.24120662 | 0.00226525 | 0.03953544 | 1 |
| 349 | Chrnd | 160 | 0.23968747 | 0.00226802 | 0.03954121 | 1 |
| 1561 | Tex11 | 159 | -0.2404228 | 0.00226868 | 0.03954121 | 1 |
| 890 | LOC432635 | 157 | -0.2418579 | 0.00227556 | 0.03963121 | 1 |
| 775 | Igfbp5 | 159 | -0.2403268 | 0.00227819 | 0.03963121 | 1 |
| 1613 | Treml1 | 156 | -0.2425907 | 0.00227851 | 0.03963121 | 1 |
| 42 | 4933400C23Rik | 158 | -0.2409324 | 0.0022924 | 0.03984548 | 1 |
| 651 | Gm11538 | 158 | 0.24089024 | 0.00229659 | 0.03989119 | 1 |
| 516 | Emb | 160 | -0.2393631 | 0.00230039 | 0.03992983 | 1 |
| 422 | Cwf19l2 | 158 | -0.2408213 | 0.00230347 | 0.03995616 | 1 |
| 1666 | Vcam1 | 160 | -0.2392967 | 0.00230707 | 0.03999134 | 1 |
| 1202 | Pkia | 142 | -0.2537466 | 0.00231103 | 0.04003271 | 1 |
| 1661 | Uts2b | 160 | -0.2392319 | 0.0023136 | 0.04004996 | 1 |
| 1515 | Suv39h1 | 156 | -0.2422061 | 0.00231666 | 0.04007568 | 1 |
| 1644 | Ublcp1 | 160 | -0.2391605 | 0.00232082 | 0.0401007 | 1 |
| 1603 | Tnfsf14 | 155 | 0.24292891 | 0.00232125 | 0.0401007 | 1 |
| 1527 | Syt2 | 160 | -0.2391018 | 0.00232677 | 0.04015638 | 1 |
| 582 | Fbxo2 | 158 | 0.24054153 | 0.0023316 | 0.04015638 | 1 |
| 629 | Gar1 | 159 | -0.2397659 | 0.0023345 | 0.04015638 | 1 |
| 1598 | Tmem64 | 160 | -0.2390195 | 0.00233514 | 0.04015638 | 1 |
| 1068 | Nfil3 | 152 | 0.24513055 | 0.00233608 | 0.04015638 | 1 |
| 843 | Klk14 | 156 | -0.242011 | 0.00233622 | 0.04015638 | 1 |
| 214 | Baiap3 | 160 | 0.23899252 | 0.00233789 | 0.04015638 | 1 |

| | | | | | | |
|---|---|---|---|---|---|---|
| 710 | H2-Ab1 | 159 | 0.23971394 | 0.00233978 | 0.04015638 | 1 |
| 78 | Acat3 | 156 | -0.24196 | 0.00234136 | 0.04015638 | 1 |
| 938 | Macrod1 | 155 | 0.24272462 | 0.00234172 | 0.04015638 | 1 |
| 1451 | Smchd1 | 158 | -0.2404406 | 0.00234182 | 0.04015638 | 1 |
| 329 | Cenpq | 149 | 0.24744798 | 0.00234614 | 0.04020329 | 1 |
| 508 | Eif4a3 | 160 | 0.23889518 | 0.00234783 | 0.04020516 | 1 |
| 1698 | Wwc1 | 160 | -0.2387627 | 0.00236141 | 0.04041063 | 1 |
| 346 | Chrna2 | 160 | 0.23872353 | 0.00236545 | 0.04045253 | 1 |
| 347 | Chrna3 | 159 | -0.2394374 | 0.00236805 | 0.04046987 | 1 |
| 91 | Acvrl1 | 160 | -0.2386249 | 0.00237563 | 0.04057162 | 1 |
| 905 | LOC433701 | 158 | -0.2400945 | 0.00237719 | 0.04057162 | 1 |
| 1735 | Zfp609 | 157 | -0.240782 | 0.00238389 | 0.04065871 | 1 |
| 412 | Csnk1g3 | 156 | -0.2414672 | 0.00239156 | 0.0407434 | 1 |
| 1539 | Tas2r134 | 158 | 0.23995038 | 0.00239206 | 0.0407434 | 1 |
| 864 | Lef1 | 61 | 0.38175679 | 0.00239889 | 0.04083102 | 1 |
| 1528 | Syt3 | 160 | -0.2383864 | 0.00240041 | 0.04083102 | 1 |
| 1578 | Tmem101 | 158 | 0.23984164 | 0.00240333 | 0.04085346 | 1 |
| 252 | C87499 | 154 | 0.24285829 | 0.0024069 | 0.04088687 | 1 |
| 38 | 4930463O16Rik | 159 | -0.2390379 | 0.00240946 | 0.04090292 | 1 |
| 1299 | Reep6 | 152 | 0.24433121 | 0.00241682 | 0.04096701 | 1 |
| 667 | Gm973 | 158 | -0.2396909 | 0.00241904 | 0.04096701 | 1 |
| 1724 | Zcchc9 | 152 | 0.24430922 | 0.00241908 | 0.04096701 | 1 |
| 331 | Cep41 | 154 | 0.24273447 | 0.00241967 | 0.04096701 | 1 |
| 233 | Blzf1 | 159 | -0.2389082 | 0.00242304 | 0.04099683 | 1 |
| 1646 | Uck1 | 160 | -0.2381536 | 0.00242483 | 0.04099987 | 1 |
| 847 | Kpna2 | 160 | -0.2381084 | 0.00242959 | 0.04102977 | 1 |
| 1162 | Pcm1 | 160 | -0.2381062 | 0.00242982 | 0.04102977 | 1 |
| 523 | Enpp2 | 160 | -0.2380733 | 0.0024333 | 0.0410613 | 1 |
| 390 | Cops3 | 159 | -0.2387112 | 0.0024438 | 0.04121113 | 1 |
| 1306 | Rgma | 160 | -0.2379529 | 0.00244605 | 0.04122186 | 1 |
| 451 | Desi2 | 160 | -0.2379197 | 0.00244958 | 0.04125403 | 1 |
| 15 | 2310030G06Rik | 156 | 0.24088987 | 0.00245159 | 0.04126064 | 1 |
| 1026 | Mup4 | 156 | 0.2408483 | 0.00245597 | 0.04130698 | 1 |
| 737 | Hipk2 | 158 | -0.2393091 | 0.00245924 | 0.04133479 | 1 |
| 1279 | Rad51 | 155 | -0.241456 | 0.00247253 | 0.04153066 | 1 |
| 150 | Apobec3 | 158 | 0.23911676 | 0.00247973 | 0.04162411 | 1 |
| 1333 | Rnf217 | 159 | -0.2383319 | 0.00248423 | 0.04167219 | 1 |
| 32 | 4833403I15Rik | 152 | -0.2436649 | 0.00248606 | 0.04167544 | 1 |
| 1433 | Slc39a7 | 159 | -0.2382886 | 0.00248888 | 0.04169524 | 1 |
| 1085 | Npepl1 | 158 | -0.2390066 | 0.00249152 | 0.04171216 | 1 |
| 482 | Dstn | 160 | -0.2375063 | 0.0024939 | 0.04171965 | 1 |
| 416 | Ctnnb1 | 160 | -0.2374772 | 0.00249704 | 0.04171965 | 1 |
| 1330 | Rnf121 | 153 | 0.2427692 | 0.00249762 | 0.04171965 | 1 |
| 481 | Dsg4 | 160 | -0.2374531 | 0.00249965 | 0.04171965 | 1 |
| 1596 | Tmem25 | 160 | -0.237441 | 0.00250097 | 0.04171965 | 1 |

| | | | | | | |
|---|---|---|---|---|---|---|
| 1437 | Slc4a1 | 155 | 0.24118044 | 0.0025018 | 0.04171965 | 1 |
| 158 | Aqp4 | 156 | -0.2403992 | 0.00250369 | 0.04172384 | 1 |
| 1541 | Tbc1d16 | 159 | 0.2381321 | 0.00250576 | 0.04173102 | 1 |
| 1622 | Tspyl4 | 160 | -0.2372582 | 0.00252084 | 0.04195472 | 1 |
| 302 | Cdan1 | 159 | -0.2379559 | 0.0025249 | 0.04199477 | 1 |
| 663 | Gm597 | 159 | -0.2378484 | 0.00253664 | 0.04214842 | 1 |
| 437 | Dcbld1 | 155 | 0.24084883 | 0.00253745 | 0.04214842 | 1 |
| 84 | Acot11 | 158 | -0.2385467 | 0.00254134 | 0.04217823 | 1 |
| 1697 | Wscd1 | 158 | -0.2385356 | 0.00254255 | 0.04217823 | 1 |
| 374 | Cnnm2 | 156 | -0.2399917 | 0.00254771 | 0.04223629 | 1 |
| 1208 | Pld2 | 156 | 0.23992142 | 0.00255538 | 0.04232319 | 1 |
| 1522 | Synj2bp | 160 | -0.2369355 | 0.00255628 | 0.04232319 | 1 |
| 1460 | Snap25 | 159 | -0.2376071 | 0.00256317 | 0.04240973 | 1 |
| 795 | Irx4 | 160 | -0.236846 | 0.00256618 | 0.042432 | 1 |
| 1150 | Pafah2 | 155 | 0.24055083 | 0.00256987 | 0.04246538 | 1 |
| 327 | Cenpf | 155 | -0.2405328 | 0.00257184 | 0.04247042 | 1 |
| 1052 | Ncald | 160 | -0.2367058 | 0.00258177 | 0.0426068 | 1 |
| 1383 | Serpinf1 | 156 | -0.2396335 | 0.00258699 | 0.04266526 | 1 |
| 1292 | Rbmy | 158 | -0.2381149 | 0.00258893 | 0.04266965 | 1 |
| 1156 | Parvg | 152 | 0.24268841 | 0.00259076 | 0.04267219 | 1 |
| 29 | 3300005D01Rik | 156 | -0.2395815 | 0.00259274 | 0.04267713 | 1 |
| 1171 | Pdgfrb | 156 | -0.2395422 | 0.00259709 | 0.04272115 | 1 |
| 904 | LOC433666 | 158 | -0.2379985 | 0.0026019 | 0.04277257 | 1 |
| 1551 | Tcf21 | 158 | 0.23791557 | 0.00261118 | 0.0428974 | 1 |
| 742 | Hnrnpa3 | 159 | -0.2371471 | 0.00261444 | 0.0429234 | 1 |
| 654 | Gm205 | 159 | -0.2371116 | 0.00261844 | 0.04293193 | 1 |
| 179 | Arrdc3 | 160 | -0.2363539 | 0.00262129 | 0.04293193 | 1 |
| 981 | Miat | 146 | -0.2472557 | 0.00262297 | 0.04293193 | 1 |
| 1062 | Nek11 | 158 | -0.237809 | 0.00262314 | 0.04293193 | 1 |
| 1060 | Nedd9 | 160 | -0.2363274 | 0.00262429 | 0.04293193 | 1 |
| 1268 | Ptpn21 | 158 | -0.2377917 | 0.00262508 | 0.04293193 | 1 |
| 962 | Mc4r | 158 | 0.23768988 | 0.00263657 | 0.04309212 | 1 |
| 353 | Ciita | 158 | -0.2376733 | 0.00263845 | 0.04309518 | 1 |
| 1092 | Nrxn1 | 160 | -0.2361817 | 0.00264083 | 0.04309618 | 1 |
| 1128 | Olfr705 | 159 | -0.2368912 | 0.00264336 | 0.04309618 | 1 |
| 277 | Ccdc171 | 160 | -0.2361513 | 0.00264429 | 0.04309618 | 1 |
| 264 | Carhsp1 | 155 | 0.23987071 | 0.00264528 | 0.04309618 | 1 |
| 156 | Aqp2 | 160 | -0.2360783 | 0.00265262 | 0.04318819 | 1 |
| 1482 | Spata16 | 159 | 0.23677706 | 0.00265636 | 0.04322137 | 1 |
| 31 | 4732414G09Rik | 156 | 0.23893877 | 0.00266474 | 0.04333014 | 1 |
| 1312 | Rhno1 | 160 | 0.23578815 | 0.00268597 | 0.04363514 | 1 |
| 1082 | Nod2 | 160 | -0.235768 | 0.0026883 | 0.04363514 | 1 |
| 310 | Cdh24 | 160 | -0.235765 | 0.00268864 | 0.04363514 | 1 |
| 605 | Frs3 | 160 | -0.2357059 | 0.00269548 | 0.04365558 | 1 |
| 1635 | Tymp | 154 | -0.2401804 | 0.00269694 | 0.04365558 | 1 |

| | | | | | | |
|---|---|---|---|---|---|---|
| 509 | Eif4e2 | 158 | -0.2371553 | 0.0026976 | 0.04365558 | 1 |
| 588 | Fezf1 | 160 | 0.23567689 | 0.00269885 | 0.04365558 | 1 |
| 968 | Med19 | 158 | -0.2371421 | 0.00269913 | 0.04365558 | 1 |
| 973 | Mest | 152 | 0.24168744 | 0.00270222 | 0.04365558 | 1 |
| 378 | Cntnap3 | 159 | -0.2363702 | 0.00270315 | 0.04365558 | 1 |
| 1481 | Spag8 | 159 | -0.2363662 | 0.00270362 | 0.04365558 | 1 |
| 350 | Chst14 | 160 | -0.2355768 | 0.0027105 | 0.0437232 | 1 |
| 861 | Lcat | 160 | -0.2355704 | 0.00271124 | 0.0437232 | 1 |
| 1548 | Tceal8 | 158 | -0.2369909 | 0.00271663 | 0.04378245 | 1 |
| 1119 | Olfr1357 | 159 | 0.23621556 | 0.00272112 | 0.04382714 | 1 |
| 282 | Ccdc8 | 159 | 0.2361455 | 0.00272931 | 0.04393113 | 1 |
| 514 | Elovl3 | 156 | 0.23822584 | 0.00274672 | 0.04416335 | 1 |
| 275 | Ccdc115 | 155 | 0.23897878 | 0.0027472 | 0.04416335 | 1 |
| 1632 | Tulp3 | 158 | -0.2367126 | 0.00274911 | 0.04416607 | 1 |
| 1291 | Rbmx2 | 155 | 0.23894231 | 0.00275144 | 0.04417576 | 1 |
| 249 | C5ar1 | 157 | -0.2373274 | 0.00276413 | 0.04434861 | 1 |
| 1419 | Slc25a18 | 160 | -0.2351075 | 0.00276569 | 0.04434861 | 1 |
| 1379 | Serpinb10 | 155 | 0.23880429 | 0.00276755 | 0.04435051 | 1 |
| 524 | Enpp5 | 156 | -0.2379948 | 0.00277378 | 0.04442225 | 1 |
| 174 | Arl16 | 160 | -0.2350248 | 0.00277552 | 0.04442225 | 1 |
| 360 | Clec18a | 160 | -0.2349684 | 0.00278224 | 0.04450193 | 1 |
| 394 | Cox6c | 160 | -0.2348818 | 0.00279259 | 0.04462266 | 1 |
| 1627 | Ttll5 | 134 | -0.2563354 | 0.0027933 | 0.04462266 | 1 |
| 1544 | Tbl3 | 160 | -0.2347759 | 0.0028053 | 0.04477861 | 1 |
| 965 | Mcm4 | 155 | 0.23847291 | 0.00280658 | 0.04477861 | 1 |
| 748 | Hoxa9 | 10 | -0.8322527 | 0.00281442 | 0.04485483 | 1 |
| 1732 | Zfp354b | 159 | -0.2354238 | 0.00281488 | 0.04485483 | 1 |
| 142 | Ap1b1 | 160 | -0.2346348 | 0.0028223 | 0.04494503 | 1 |
| 625 | Galk1 | 154 | 0.23906512 | 0.0028268 | 0.04498853 | 1 |
| 436 | Dbx1 | 156 | -0.2375153 | 0.00283068 | 0.04502214 | 1 |
| 1185 | Pex19 | 140 | -0.250404 | 0.00284557 | 0.04523066 | 1 |
| 1728 | Zfp180 | 158 | -0.2358767 | 0.0028488 | 0.04524894 | 1 |
| 860 | Larp1b | 160 | -0.2343929 | 0.00285168 | 0.04524894 | 1 |
| 1262 | Ptgfrn | 138 | -0.2521286 | 0.00285205 | 0.04524894 | 1 |
| 472 | Dnali1 | 158 | -0.2357972 | 0.00285846 | 0.04532237 | 1 |
| 459 | Dlk1 | 160 | 0.23427202 | 0.00286647 | 0.04541193 | 1 |
| 613 | Gaa | 160 | 0.23426221 | 0.00286767 | 0.04541193 | 1 |
| 1504 | Stmn4 | 160 | -0.2342314 | 0.00287145 | 0.04544352 | 1 |
| 171 | Arid5b | 158 | -0.2356646 | 0.00287461 | 0.04546521 | 1 |
| 1619 | Tsc2 | 160 | -0.2341763 | 0.00287822 | 0.04549414 | 1 |
| 341 | Chchd3 | 160 | -0.2341385 | 0.00288287 | 0.04553938 | 1 |
| 1573 | Tle3 | 155 | -0.2378133 | 0.00288574 | 0.04555644 | 1 |
| 593 | Fgf13 | 160 | -0.2340754 | 0.00289066 | 0.04556108 | 1 |
| 823 | Kcnj5 | 160 | -0.2340507 | 0.00289372 | 0.04556108 | 1 |
| 1684 | Wac | 158 | -0.2355035 | 0.00289435 | 0.04556108 | 1 |

| | | | | | | |
|---|---|---|---|---|---|---|
| 1040 | N4bp2l1 | 147 | 0.2440378 | 0.00289462 | 0.04556108 | 1 |
| 950 | Map3k4 | 158 | -0.2354984 | 0.00289498 | 0.04556108 | 1 |
| 796 | Isca2 | 160 | -0.2340241 | 0.00289701 | 0.04556487 | 1 |
| 1037 | Myrfl | 159 | -0.2347085 | 0.00290209 | 0.04561649 | 1 |
| 48 | 5730455P16Rik | 158 | -0.2354056 | 0.00290642 | 0.04565117 | 1 |
| 752 | Hsd17b2 | 156 | 0.23687868 | 0.00290788 | 0.04565117 | 1 |
| 771 | Ifnar1 | 160 | -0.233914 | 0.00291067 | 0.04566052 | 1 |
| 589 | Fezf1 | 160 | 0.23390276 | 0.00291206 | 0.04566052 | 1 |
| 1206 | Plcl1 | 159 | -0.2345734 | 0.00291883 | 0.04573859 | 1 |
| 1536 | Taok3 | 158 | 0.23525567 | 0.00292497 | 0.04580658 | 1 |
| 1571 | Tktl2 | 157 | -0.2358831 | 0.00293875 | 0.04594459 | 1 |
| 655 | Gm2a | 160 | -0.2336866 | 0.00293906 | 0.04594459 | 1 |
| 924 | Lrp1 | 160 | -0.2336854 | 0.0029392 | 0.04594459 | 1 |
| 931 | Ltn1 | 160 | -0.2335597 | 0.002955 | 0.04616322 | 1 |
| 768 | Idh1 | 158 | 0.23492853 | 0.00296583 | 0.04629561 | 1 |
| 812 | Jund | 160 | -0.2334638 | 0.00296711 | 0.04629561 | 1 |
| 1640 | Ube2b | 160 | -0.2333455 | 0.0029821 | 0.0464545 | 1 |
| 688 | Gprc5b | 160 | -0.2333364 | 0.00298326 | 0.0464545 | 1 |
| 967 | Med1 | 159 | -0.2340553 | 0.00298385 | 0.0464545 | 1 |
| 410 | Cryba2 | 159 | -0.2340494 | 0.00298459 | 0.0464545 | 1 |
| 766 | Id1 | 158 | -0.2347421 | 0.00298934 | 0.0464814 | 1 |
| 1708 | Ythdf2 | 154 | 0.237727 | 0.00299005 | 0.0464814 | 1 |
| 25 | 2700081O15Rik | 160 | -0.2332667 | 0.00299212 | 0.0464814 | 1 |
| 1526 | Syt11 | 160 | -0.233255 | 0.00299362 | 0.0464814 | 1 |
| 719 | Hax1 | 160 | -0.2331411 | 0.00300817 | 0.0466789 | 1 |
| 348 | Chrna3 | 160 | -0.2331196 | 0.00301093 | 0.04669329 | 1 |
| 1672 | Vmn1r66 | 159 | -0.2337958 | 0.0030169 | 0.0467168 | 1 |
| 537 | Erlin2 | 155 | 0.2367526 | 0.00301728 | 0.0467168 | 1 |
| 417 | Ctps | 145 | -0.2446664 | 0.00301812 | 0.0467168 | 1 |
| 112 | Aire | 160 | 0.23305057 | 0.00301979 | 0.0467168 | 1 |
| 443 | Ddx42 | 156 | -0.235961 | 0.0030225 | 0.04672541 | 1 |
| 260 | Cald1 | 158 | 0.23445092 | 0.00302641 | 0.04672541 | 1 |
| 817 | Kcnab2 | 160 | -0.2329799 | 0.00302889 | 0.04672541 | 1 |
| 1136 | Ormdl3 | 156 | 0.23590682 | 0.00302939 | 0.04672541 | 1 |
| 1099 | Nubp1 | 160 | 0.23297495 | 0.00302952 | 0.04672541 | 1 |
| 519 | Emilin2 | 156 | 0.23585016 | 0.00303662 | 0.04678698 | 1 |
| 1611 | Tprkb | 159 | -0.2336378 | 0.00303719 | 0.04678698 | 1 |
| 920 | LOC546168 | 157 | -0.2350688 | 0.00304154 | 0.04682191 | 1 |
| 733 | Hhip | 152 | -0.2388422 | 0.00304313 | 0.04682191 | 1 |
| 184 | Asb8 | 160 | -0.2328084 | 0.00305106 | 0.04691553 | 1 |
| 1077 | Nisch | 160 | -0.232754 | 0.00305811 | 0.04699562 | 1 |
| 561 | Fam221b | 157 | 0.23491728 | 0.00306102 | 0.04701201 | 1 |
| 1549 | Tcerg1 | 160 | -0.2327071 | 0.00306421 | 0.04703256 | 1 |
| 463 | Dmd | 159 | -0.2333975 | 0.00306828 | 0.04706667 | 1 |
| 1151 | Pald1 | 160 | -0.2326565 | 0.0030708 | 0.04707704 | 1 |

| | | | | | | |
|---|---|---|---|---|---|---|
| 1323 | Ripk4 | 157 | -0.2347922 | 0.00307719 | 0.04711857 | 1 |
| 1628 | Ttll6 | 158 | -0.234057 | 0.00307721 | 0.04711857 | 1 |
| 1239 | Ppp3r1 | 159 | -0.2332569 | 0.00308661 | 0.04720753 | 1 |
| 534 | Epm2aip1 | 157 | 0.23471866 | 0.00308673 | 0.04720753 | 1 |
| 119 | Aldh18a1 | 152 | 0.23841509 | 0.00309753 | 0.04730755 | 1 |
| 401 | Cpt2 | 157 | -0.2346197 | 0.00309961 | 0.04730755 | 1 |
| 1713 | Ywhag | 138 | -0.2500218 | 0.00310149 | 0.04730755 | 1 |
| 194 | Atg9b | 156 | -0.2353447 | 0.00310176 | 0.04730755 | 1 |
| 635 | Gbp2 | 156 | -0.2353361 | 0.00310287 | 0.04730755 | 1 |
| 1476 | Sorcs2 | 160 | -0.2324001 | 0.00310442 | 0.04730755 | 1 |
| 788 | Iqch | 158 | -0.2337961 | 0.00311128 | 0.04737656 | 1 |
| 1461 | Snap91 | 160 | -0.232336 | 0.00311288 | 0.04737656 | 1 |
| 787 | Iqcg | 158 | -0.2337714 | 0.00311453 | 0.04737656 | 1 |
| 241 | Brs3 | 158 | 0.23375643 | 0.0031165 | 0.04737818 | 1 |
| 64 | A930012M21Rik | 158 | -0.2337176 | 0.0031216 | 0.04742743 | 1 |
| 913 | LOC436099 | 160 | -0.2322484 | 0.00312447 | 0.0474427 | 1 |
| 806 | Izumo3 | 159 | -0.232933 | 0.0031292 | 0.04748626 | 1 |
| 844 | Klk1b8 | 155 | 0.23581454 | 0.00313809 | 0.04753102 | 1 |
| 1668 | Vdac2 | 160 | -0.23213 | 0.0031402 | 0.04753102 | 1 |
| 1309 | Rgs9 | 158 | 0.23357419 | 0.00314053 | 0.04753102 | 1 |
| 1313 | Rhobtb2 | 158 | -0.2335665 | 0.00314156 | 0.04753102 | 1 |
| 1234 | Ppp1r13b | 159 | -0.2328391 | 0.00314164 | 0.04753102 | 1 |
| 1296 | Rdh10 | 156 | -0.2350241 | 0.00314372 | 0.04753102 | 1 |
| 126 | Alg9 | 155 | 0.23576023 | 0.00314522 | 0.04753102 | 1 |
| 145 | Apba2 | 160 | -0.2320007 | 0.00315745 | 0.04766468 | 1 |
| 1633 | Txndc11 | 156 | 0.23490695 | 0.00315918 | 0.04766468 | 1 |
| 328 | Cenpm | 156 | 0.23490321 | 0.00315968 | 0.04766468 | 1 |
| 424 | Cyc1 | 160 | -0.2319481 | 0.00316449 | 0.04769989 | 1 |
| 550 | F8a | 156 | -0.2348573 | 0.00316576 | 0.04769989 | 1 |
| 893 | LOC432742 | 159 | -0.2325788 | 0.00317637 | 0.04783152 | 1 |
| 1169 | Pdgfra | 159 | -0.2325365 | 0.00318205 | 0.0478887 | 1 |
| 1655 | Usp18 | 158 | -0.2332261 | 0.00318691 | 0.04793348 | 1 |
| 1445 | Slc7a6 | 159 | 0.23237233 | 0.00320418 | 0.04814294 | 1 |
| 1221 | Pofut2 | 160 | -0.2316505 | 0.00320461 | 0.04814294 | 1 |
| 545 | Exosc9 | 159 | 0.23230445 | 0.00321336 | 0.04823614 | 1 |
| 340 | Chaf1a | 160 | 0.23156494 | 0.00321623 | 0.04823614 | 1 |
| 1008 | Mrpl23 | 159 | -0.2322758 | 0.00321724 | 0.04823614 | 1 |
| 323 | Ceacam3 | 160 | 0.23154672 | 0.00321871 | 0.04823614 | 1 |
| 1681 | Vwa3a | 159 | -0.2322534 | 0.00322029 | 0.04823614 | 1 |
| 8 | 1700123I01Rik | 158 | 0.23292166 | 0.00322798 | 0.04830686 | 1 |
| 868 | Lepr | 158 | -0.2329156 | 0.0032288 | 0.04830686 | 1 |
| 515 | Elp2 | 156 | -0.2343208 | 0.00323757 | 0.04838482 | 1 |
| 1113 | Oat | 160 | -0.2314035 | 0.00323825 | 0.04838482 | 1 |
| 47 | 5530401A14Rik | 159 | -0.2321107 | 0.00323972 | 0.04838482 | 1 |
| 1745 | Zfpm2 | 160 | -0.2313336 | 0.00324784 | 0.04847772 | 1 |

| | | | | | | |
|---|---|---|---|---|---|---|
| 1121 | Olfr1494 | 159 | -0.232006 | 0.00325403 | 0.04854173 | 1 |
| 644 | Gjb2 | 160 | -0.2312266 | 0.00326255 | 0.04863642 | 1 |
| 1362 | Samhd1 | 158 | 0.232656 | 0.0032642 | 0.04863642 | 1 |
| 907 | LOC433761 | 152 | 0.23711326 | 0.0032688 | 0.04864896 | 1 |
| 94 | Adamts19 | 160 | 0.23118082 | 0.00326886 | 0.04864896 | 1 |
| 1235 | Ppp1r15a | 154 | -0.2354773 | 0.00328378 | 0.04881338 | 1 |
| 337 | Cfh | 157 | -0.2332308 | 0.00328559 | 0.04881338 | 1 |
| 1256 | Psenen | 157 | -0.2332249 | 0.00328641 | 0.04881338 | 1 |
| 1394 | Sh3gl2 | 160 | -0.2310456 | 0.00328758 | 0.04881338 | 1 |
| 743 | Hnrnpf | 156 | -0.2339346 | 0.00329018 | 0.04882356 | 1 |
| 619 | Gad1 | 158 | 0.23243876 | 0.0032941 | 0.04884924 | 1 |
| 639 | Gfral | 159 | -0.2317034 | 0.00329575 | 0.04884924 | 1 |
| 13 | 2310002L09Rik | 156 | 0.23387744 | 0.00329804 | 0.04885472 | 1 |
| 993 | Mmp8 | 159 | -0.2316338 | 0.00330541 | 0.04891516 | 1 |
| 645 | Gjd3 | 160 | -0.2309109 | 0.00330632 | 0.04891516 | 1 |
| 216 | BC048671 | 156 | 0.23380596 | 0.00330788 | 0.04891516 | 1 |
| 818 | Kcnd2 | 158 | -0.2322765 | 0.00331659 | 0.04896143 | 1 |
| 698 | Grm8 | 158 | -0.2322664 | 0.00331799 | 0.04896143 | 1 |
| 1314 | Rhot1 | 160 | -0.230824 | 0.00331845 | 0.04896143 | 1 |
| 452 | Dgcr8 | 157 | -0.232991 | 0.0033187 | 0.04896143 | 1 |
| 1630 | Ttyh3 | 158 | -0.2321392 | 0.00333572 | 0.04918401 | 1 |
| 1029 | Mut | 157 | -0.2328475 | 0.00333866 | 0.04919882 | 1 |
| 939 | Mad2l1 | 158 | -0.2320836 | 0.0033435 | 0.04922386 | 1 |
| 800 | Itga10 | 158 | 0.23207847 | 0.00334422 | 0.04922386 | 1 |
| 1667 | Vcpkmt | 159 | 0.23132967 | 0.00334794 | 0.0492501 | 1 |
| 500 | Efhc1 | 158 | -0.2320308 | 0.0033509 | 0.04926522 | 1 |
| 67 | Aagab | 159 | -0.2312268 | 0.00336243 | 0.0494052 | 1 |
| 289 | Ccni | 146 | -0.2411471 | 0.0033663 | 0.0494052 | 1 |
| 1485 | Spdef | 145 | -0.2419663 | 0.00336633 | 0.0494052 | 1 |
| 368 | Cluap1 | 157 | -0.2326366 | 0.00336818 | 0.0494052 | 1 |
| 236 | Bms1 | 157 | -0.2326033 | 0.00337287 | 0.04944541 | 1 |
| 397 | Cpa2 | 158 | 0.23178675 | 0.00338531 | 0.04955697 | 1 |
| 1513 | Sun2 | 160 | 0.23034544 | 0.00338602 | 0.04955697 | 1 |
| 781 | Il1rl2 | 160 | 0.2303434 | 0.00338631 | 0.04955697 | 1 |
| 1058 | Necab1 | 160 | -0.2302733 | 0.00339632 | 0.04967483 | 1 |
| 1106 | Nwd2 | 160 | -0.2302505 | 0.00339957 | 0.04968991 | 1 |
| 693 | Grin2d | 160 | -0.2302388 | 0.00340125 | 0.04968991 | 1 |
| 1199 | Pip4k2c | 160 | 0.23016096 | 0.0034124 | 0.04981638 | 1 |
| 562 | Fam222b | 138 | -0.2475725 | 0.0034161 | 0.04981638 | 1 |
| 1458 | Smyd2 | 158 | -0.231568 | 0.00341642 | 0.04981638 | 1 |
| 386 | Col9a1 | 152 | 0.23603022 | 0.00341774 | 0.04981638 | 1 |
| 217 | BC049702 | 158 | -0.2315226 | 0.00342291 | 0.04986324 | 1 |
| 600 | Foxk1 | 158 | -0.2314896 | 0.00342764 | 0.04988292 | 1 |
| 1015 | Msi2 | 160 | -0.2300512 | 0.00342818 | 0.04988292 | 1 |
| 1381 | Serpinb1b | 160 | -0.2300004 | 0.00343552 | 0.04992922 | 1 |

| | | | | | | |
|---|---|---|---|---|---|---|
| 97 | Adcy5 | 157 | 0.23215687 | 0.00343623 | 0.04992922 | 1 |
| 1747 | Zfyve19 | 158 | 0.23141321 | 0.00343859 | 0.04992922 | 1 |
| 951 | Map3k6 | 157 | -0.2321361 | 0.0034392 | 0.04992922 | 1 |
| 1251 | Prm1 | 158 | -0.2313642 | 0.00344563 | 0.049994 | 1 |
